# Practical scan-length considerations for mapping upper limb movements to the somatosensory/motor cortex at 7T

**DOI:** 10.1101/2022.09.11.507497

**Authors:** D Rangaprakash, Olivia E Rowe, Hyungeun Song, Samantha Gutierrez-Arango, Michael F Fernandez, Erica A Israel, Hugh M Herr, Robert L Barry

## Abstract

The relationship between motor cortex (M1) and upper limb movements has been investigated extensively using functional MRI (fMRI). While most research has focused on applications, very few studies have focused on practical aspects related to developing the fMRI protocol. Thus, the effect of scan length on M1 activations during various upper limb movements remains unclear. Scan length constraints are important for conducting motor experiments within a 60- or 90-min scan session. We targeted this gap by studying 7T fMRI activations in a male participant while performing eight different upper limb movements (of the fingers, wrist, and elbow) across 16 task runs (8 with the left arm, 8 with the right arm, 88 mins total fMRI duration). Standard activation analyses were performed (*Z*>3.1, *p*<0.01, cluster thresholded) independently for 14 different cases (2 runs through 8 runs, left and right arm) and compared. We found diminishing returns with higher number of runs (activations gradually plateaued with runs). We observed two clusters of movements, one with generally higher activation (more activated voxels and higher Z-stats) and the other with lower activation. To achieve similar statistical power, movements with lower activation required longer scanning (more runs). Based on these observations, we propose a ‘*one size does not fit all*’ practical protocol within a 60-, 75-, or 90-min scan session, wherein different number of runs are assigned for different movements. Our study could benefit researchers who are designing upper limb fMRI experiments.

## 1. Introduction

Somatotopic mapping typically refers to the organized representation of body movements in the primary motor cortex (M1), and body sensations in the primary somatosensory cortex (S1). Mapping bodily movements/sensations, including the upper limb, to different areas of the highly organized M1/S1 has been of interest for several decades, and more recently has sustained interest with critical advancements through basic neuroscience and clinical psychiatry/neurology research. Recent advancements in motor somatotopy have been brought about largely through blood oxygenation level dependent (BOLD) functional magnetic resonance imaging (fMRI).

Somatosensory somatotopy broadly exhibits a one-to-one mapping between body locations and corresponding S1 activations [1]. A few studies have even demonstrated high-precision mapping of finger sensory stimulation with limited overlap between fingers [2] [3]. In contrast, motor somatotopy is not so straightforward. Since motor movements generally elicit somatosensory stimulation [4], neuronal activity in corresponding M1 and S1 regions is often tightly coupled. Hence, fMRI studies on motor somatotopy often report activations in both M1 and S1. So far, there have been several studies on mapping upper limb movements to M1/S1. Besle et al. [4] found largely overlapping cortical activations across each of the finger movements using an event-related fMRI task design. Roux et al.’s study using invasive electrostimulation found M1 representations of hand movement to overlap considerably with those of wrist and elbow movements [5]. Other studies have mirrored these observations with both event-related and block designs [6] [7] [8] [9] [10]. The current understanding is that rather than individual neurons controlling individual muscles, M1 neurons encode the building blocks of complex movements, which overlap across different limb movements. That is, there exists many-to-many mapping between M1 and body movements [5].

Despite these complexities in motor somatotopic mapping, precision motor somatotopy has been attempted more recently using longer scanning times, acquisition modifications (e.g., targeted spatial coverage), and advanced analysis techniques. Schellekens et al. [11] mapped digit representations in M1/S1 using high spatial resolution 7T fMRI with limited field of view, employing a Gaussian population receptive field model. Huber et al. [12] recently used sub-millimeter vascular space occupancy (VASO) fMRI (has poorer sensitivity but higher specificity compared to BOLD [13]) with a limited field of view to scan each participant for 84 hours at 7T. They found digit representations across M1 laminar columns with high spatial precision. For further reading on latest research, please refer to [5] [14] [15]. Taken together, significant advancements have been made in motor somatotopy and functional neuroimaging of upper limb movements. Thorough studies and specialized experimental designs have revealed motor mapping in M1 and have improved our understanding of the relationship between M1 activations and corresponding limb movements.

Despite this progress, there have been very few published studies with the primary goal of developing a protocol to image upper limb movements. Most studies have aimed to either advance neuroscientific understanding of motor movements or identify clinical markers of diseases involving motor impairments. Studies suggesting a practical protocol for imaging upper limb movements under various constraints and choices have been limited. Nevertheless, recent works have studied the impact of fMRI spatial resolution [16], spatial smoothing [17], task design [18], and statistical analysis [19] on M1 activations. However, the effect of scan length (different number of runs) on activation for individual motor tasks has not been assessed and compared previously. This factor becomes important for determining the scanning protocol of new studies under realistic time, cost, and patient compliance (e.g., muscle fatigue) constraints. Scan lengths varied largely across prior studies, as did the precision of activations. Firstly, for a given upper limb movement, it remains unclear as to how long one would need to scan to obtain asymptotically robust activations in M1/S1. Secondly, the optimal scan length to distinguish different upper limb movements has yet to be clarified. In this prospective study, we address this gap by presenting and comparing BOLD activations across eight different upper limb movements across different number of runs (2-8 runs, 11-44 min, performed with both left and right arms). Based on our observations, we provide recommendations on a practical protocol to image upper limb movements within a 60-, 75-, or 90-min MRI session using a routine fMRI sequence with standard acquisition parameters and whole brain coverage at 7T.

It is notable that although our study has just one participant, we performed extensive analyses using 16 runs of fMRI data involving 8 different movements, generating extensive supplementary data (40 pages) and a separately published public data resource in Mendeley Data. Thus, we consider this to be a full-fledged research study addressing an important gap in the field and not merely a case report in the traditional sense.

## 2. Methods

### 2.1. FMRI data acquisition

This study involved one male participant (age range 21-30, left-handed) who had previous fMRI experience. The protocol and procedures were approved by the Mass General Brigham Institutional Review Board (protocol #2016P002391). Informed consent was obtained prior to data acquisition. While the participant performed motor tasks, BOLD fMRI data acquisition was performed in a Siemens Magnetom Terra 7T MRI scanner (Siemens Healthcare, Erlangen, Germany) with the following parameters: 2D echo planar imaging (EPI) sequence, TR = 1190 ms, TE = 22 ms, flip angle = 60°, voxel size = 1.5×1.5×1.5 mm^3^, in-plane field of view = 190×190 mm, acquisition matrix = 128×128, 92 slices (no slice gap) with whole-brain coverage, acceleration factor = 12 (simultaneous multi-slice [SMS] slice acceleration factor = 4, GRAPPA in-plane acceleration factor = 3), scan duration = 5 min 30 sec (278 volumes) per run. High-resolution T1-weighted anatomical images were acquired using a 3D magnetization-prepared rapid acquisition of gradient echo (MPRAGE) sequence: TR/TE = 2530/1.76 ms, inversion time = 1100 ms, flip angle = 7°, voxel size = 0.75×0.75×0.75 mm^3^, field of view = 240×240 mm, acquisition matrix = 320×320. A standard Nova Medical head coil (1-channel Tx, 32-channel Rx) was used.

### 2.2. Motor task design

The block design motor task (lasting 330 sec) consisted of 8 on-off blocks, with each on (and off) period lasting for 20.625 sec (20.625×2×8 = 330). Each of the 8 blocks required the participant to perform a specific upper limb movement for the ‘on’ period and rest during the ‘off’ period. The instructions were shown on a head-mount display and coded in MATLAB using the Psychophysics Toolbox extensions [20]. Each of the blocks involved a different movement as follows (presented in this order): (i) thumb up/down (abduction/adduction), (ii) thumb open/close (flexion/extension), (iii) thumb rotation, (iv) index finger open/close (flexion/extension), (v) digits 3-5 open/close (flexion/extension), (vi) wrist prone/supine, (vii) elbow open/close (flexion/extension), and (viii) wrist open/close (flexion/extension). These movements are visually presented in **Video 1**. Please note that the terms open/close and flexion/extension are used interchangeably (same with up/down and abduction/adduction). Irrespective of what they are called, the video specifies what movements are precisely associated with these terms.

This 330-sec task constituted one run. The participant performed 8 runs with the right hand and likewise 8 runs with the left, alternating between the two (R-L-R-L-…), for a total of 16 functional runs. A short break of approximately 1 min was provided between runs. Task compliance was ensured visually from the scanner console. In the days leading up to the scan, the participant was trained on how to execute the movements and how they would be presented in the scanner. This training was reviewed with the participant on the day of the scan. The aim of this experimental setup was to (a) compare activations in M1/S1 across different movements, and (b) compare activations across different number of runs for each movement.

**Video 1.**
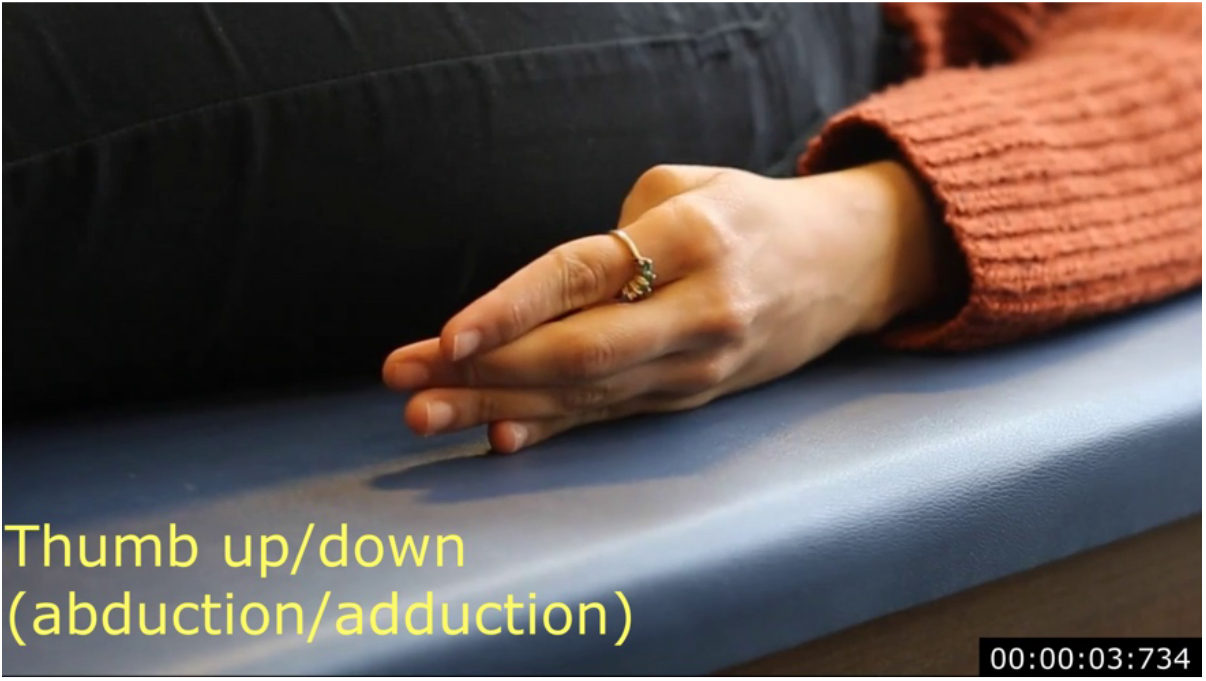
Visual illustration of the eight upper limb movements performed by the participant during the 20.625 second ‘ON’ periods of the task (shown here is the thumbnail for the purpose of presenting in the print version of the paper).

### 2.3. FMRI data pre-processing

Fourteen separate analyses were performed – 14 movements, 7 with left arm and 7 with right – and in each case with the (i) first 2 runs, (ii) first 3 runs, (iii) first 4 runs, (iv) first 5 runs, (v) first 6 runs, (vi) first 7 runs, and (vii) all runs. For the analysis of “all runs” left arm movements, the 8 runs were first concatenated across time to provide a single 3D+time image. At each voxel, the mean value of the fMRI time series was matched across all runs prior to concatenation (i.e., all runs were forced to have the same mean at a given voxel). Likewise, concatenation was performed for the other cases (2 runs, …, 7 runs), and this was repeated for the right arm movements. Further processing was performed on this concatenated image for each case.

Data were pre-processed in FSL v6.0.3 [21] using FMRI Expert Analysis Tool (FEAT) [22], with the following steps (**Supplemental Figures S1.1** and **S1.2** show FSL screenshots of these settings). Brain extraction was performed on structural images using BET to remove non-brain areas [23]. Motion correction was performed using MCFLIRT [24]. Spatial smoothing was performed using a 3-mm Gaussian kernel; a smaller kernel was chosen to preserve spatial specificity of different limb movements as much as possible. High-pass filtering was performed with a cut-off of 0.01 Hz. Two-step co-registration to the MNI space was performed, using the structural image as the intermediary [24]. Outputs of pre-processing steps (e.g., motion parameters and co-registrations) were visually inspected; no anomalies were found.

### 2.4. Activation analysis

First-level general linear model (GLM) activation analysis was performed in FEAT with 8 explanatory variables (EVs) – each of them corresponding to one of the eight limb movements. Standard and extended motion parameters were used as regressors of no interest (**Supplemental Figures S1.3** and **S1.4** show the task designs). Activations were limited to M1 and S1 using M1/S1 masks defined in the Harvard-Oxford atlas. Non-parametric statistics were performed with a voxel-level threshold of *Z*>3.1 and cluster threshold of *p*<0.01. The contrasts were set up to capture the activations corresponding to the 8 movements. The Results section presents and compares these activation maps across different limb movements and runs. Please refer to Supplemental Information section S1 for screenshots of pre-processing and activation analysis steps.

## 3. Results

Supplemental Information sections S2 and S4 provide all activation maps generated by FSL (thresholded Z-statistic maps of contrasts) for various limb movements and various runs respectively, for both left and right limbs. Along with those detailed comparisons, 3D maps and overlays generated using Mango (http://ric.uthscsa.edu/mango/) are also presented in sections S3 and S5. All thresholded Z-statistic NIfTI images are publicly available [25]. All images are presented with the radiological convention. Here we present summary statistics and similarity metrics of activations across different movements and numbers of runs.

### 3.1. Comparison of different upper limb movements

A comparison of activations across different limb movements (**Figure 1**) showed interesting patterns with both left and right arm movements. Thumb open/close and index finger open/close resulted in significantly fewer activated voxels compared to all other movements (*p*<0.02), with both left and right arm movements (**Fig. 1a**). The number of activated voxels between thumb open/close and index finger open/close did not differ (*p*>0.08). Elbow had significantly higher number of activated voxels compared to all other movements (*p*<0.02). The remaining five movements did not differ among themselves (*p*>0.06). A similar pattern was observed with Z-stats (**Fig. 1b**); thumb open/close and index did not differ (*p*>0.74) but were lower than the rest of the movements (*p*<10^−6^). Elbow had higher Z-stat than rest of the movements (*p*<10^−309^). The remaining five movements mostly differed amongst themselves, but to a smaller degree (*p*<0.54, median(*p*)=10^−4^). The bar graph, shown in **Fig. 1c**, focuses on highly significant voxels (95^th^ percentile and the median of the top 5 percentile voxels) because these are voxels corresponding to the activation peaks, which are of interest in functional neuroimaging. This bar graph of top percentile Z-stat values reflected a similar pattern. Although the 5^th^ percentile (bottom) did not differ, the 95^th^ percentile (top) Z-stat across all activated voxels was lowest for thumb open/close and index open/close, and highest for elbow and intermediate for the rest of the movements. All these observations were more consistent with the left arm movements.

**Figure 1.**
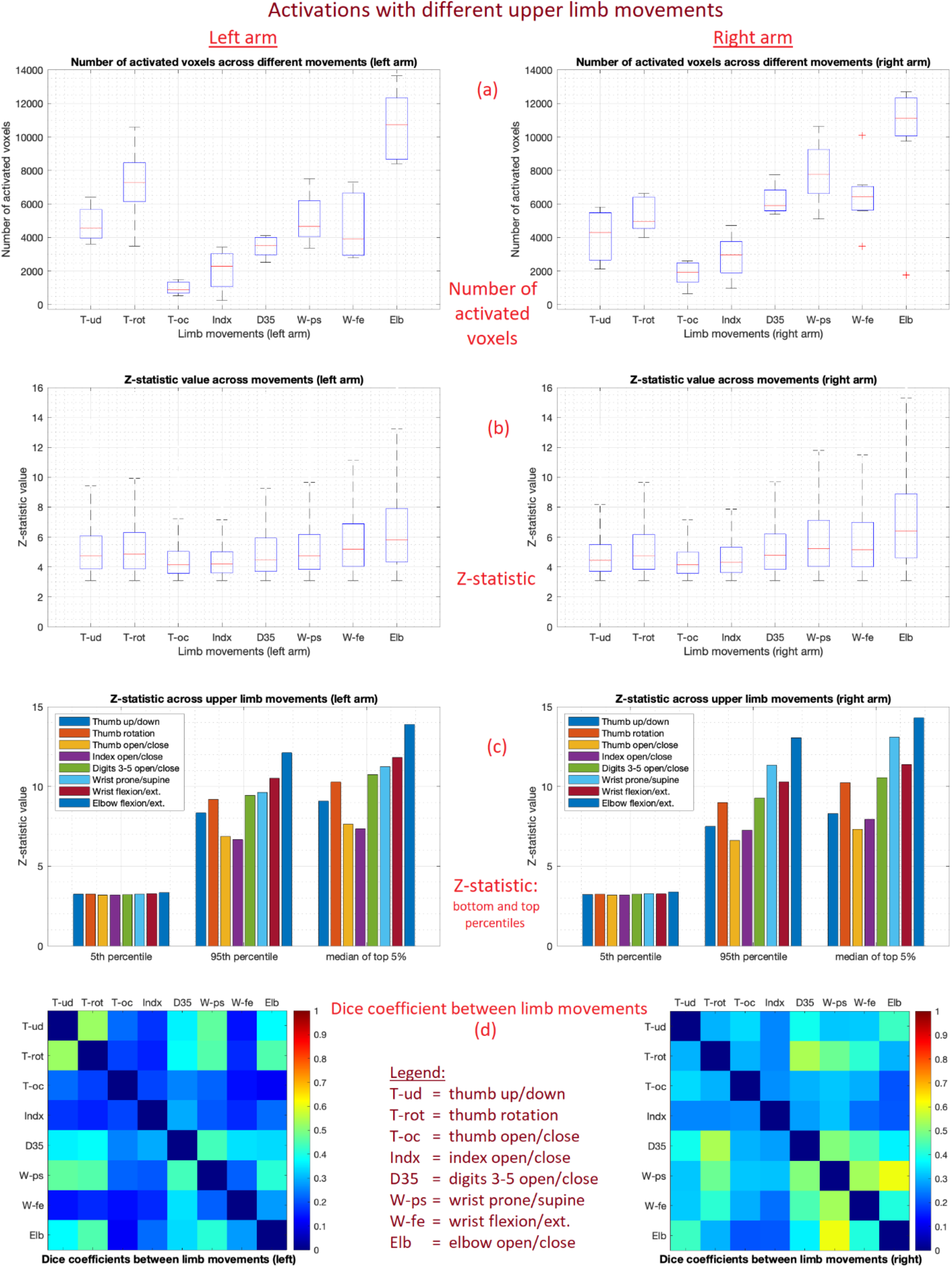
Activations with different upper limb movements (taken across different number of runs). (a) Boxplot of number of activated voxels; (b) boxplot of Z-stats in activated voxels; (c) bar graph of Z-stat’s 5^th^ percentile, 95^th^ percentile and the median of top 5% values; (d) dice coefficient (similarity) between different limb movements’ activations.

Finally, similarity in activation maps (or activated voxels) between different movements was quantified using the dice coefficient (*D*_*c*_) (**Fig. 1d**), which ranges from 0 to 1 where a value of 1 represents highest (perfect) similarity. Thumb open/close, index open/close and wrist flexion/extension had the most dissimilar activations among themselves (mean *D*_*c*_=0.21) as well as with rest of the movements (mean *D*_*c*_=0.27). These values were higher with the right arm (mean *D*_*c*_=0.31) compared to the left arm (mean *D*_*c*_=0.21). The remaining five movements had higher similarity among themselves (mean *D*_*c*_=0.42), and this was consistent across both the arms (mean *D*_*c*_: left=0.42, right=0.42).

Please refer to Supplemental Information sections S2 and S3 for activation maps and 3D maps, respectively, which reaffirm the observations made in Figure 1. Taken together, thumb open/close and index open/close showed the lowest number of activations with lower statistical power and less similarity in activation with the rest of the movements. Elbow had the highest number of activations with higher power. The remaining five movements had intermediate number of (and power of) activations.

### 3.2. Comparison of different number of runs

A comparison of activations across different number of runs (**Figure 2**) showed interesting patterns. The number of activated voxels did not differ among different runs (*p*>0.14) (**Fig. 2a**). However, significant increment in the Z-stat was observed (*p*<10^−20^) between every combination of more runs vs. fewer runs with the left arm, except where 5 runs had lower Z-stat than 3 runs and 4 runs (median *Z*: 8 runs = 5.32, 2 runs = 4.66) (**Fig. 2b**). Interestingly, with the right arm, 4 runs had the highest Z-stat (median *Z* = 5.66), with subsequently increasing number of runs resulting in lower Z-stats (2 and 3 runs also had lower Z-stats than 4 runs); all differences were significant (*p*<10^−11^). While our study was not designed to probe the reason for this observation, we suspect that this was perhaps due to a combination of handedness (difference between arms in habituation to movements) and task performance (harder to comply consistently with a greater number of runs, which might also vary with handedness).

**Figure 2.**
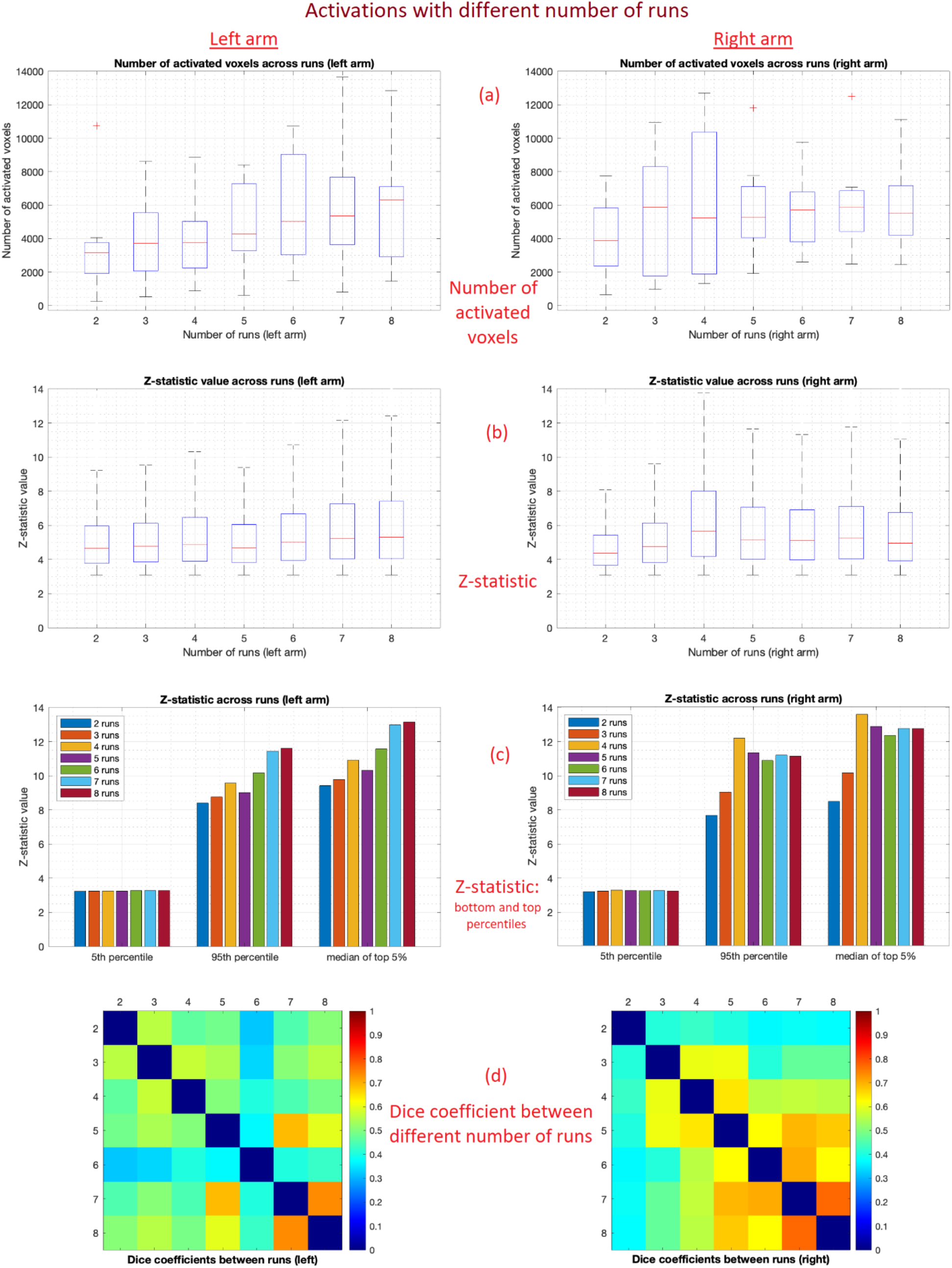
Activations with different number of runs (taken across different limb movements). (a) Boxplot of number of activated voxels; (b) boxplot of Z-stats in activated voxels; (c) bar graph of Z-stat’s 5^th^ percentile, 95^th^ percentile and the median of top 5% values; (d) dice coefficient (similarity) between activations with different number of runs.

Observing the bar graph of top percentile Z-stat values (**Fig. 2c**), there was a near-linear increase in 95^th^ percentile Z-stat across increasing number of runs with the left arm. This is expected given the higher statistical power with a greater number of runs. With the right arm, top Z-stats saturated at 4 runs. Finally, dice coefficients (**Fig. 2d**) revealed consistently similar activation patterns across runs with the left arm (mean *D*s_*c*_=0.54) except with 6 runs (mean *D*_*c*_=0.39). We suspect that task performance might have been a factor with run-6 due to the length of the experiment. With the right arm, runs 4 through 8 had similar activation patterns (mean *D*_*c*_=0.68), while 2 and 3 runs had poorer similarity with the rest (mean *D*_*c*_=0.48). This might be because subsequent runs beyond run 4 did not contribute much to improvement in estimating activation (as mentioned earlier), leading to similar activations with more runs.

Please refer to Supplemental Information sections S4 and S5 for activation maps and 3D maps, respectively, which reaffirm the observations made in Figure 2. In addition, those sections also reveal that tasks with larger activations (thumb up/down and rotation, digits 3-5, wrist prone/supine and elbow) showed remarkably better consistency of activation from 3-runs through 7-runs cases. Taken together, estimation of activation generally improves with a greater number of runs (as expected) but other factors such as task performance (exhaustion from performing the task, habituation) and handedness might play a role. Increasing number of runs might not always translate to proportional linear increase in the fidelity of estimated activations.

## 4. Discussion

Significant advancements have been made in the neuroscientific understanding of upper limb movements using fMRI. However, to the best of our knowledge, no studies have focused specifically on scan length considerations for achieving desired levels of activations in M1/S1 for upper limb movements, and subsequently their implications for the development of a practical imaging protocol that considers time, cost, and other subject specific constraints. We addressed this gap by estimating activations in M1/S1 during 8 different upper limb movements and comparing different scan lengths (2 to 8 runs, i.e., 11 mins to 44 mins). Our main findings were that (i) different movements had different strength of activations and detection powers, (ii) overlap in activations seemed to depend upon common muscle fibers between the movements, and (iii) increasing number of runs generally increased the statistical significance (Z-stat) but the gain was not proportionally higher with larger number of runs. We discuss these observations next.

i. Different movements had different detection powers (refer to **Fig. 1**). Specifically, elbow movement exhibited largest number of activated voxels and highest Z-stats. Thumb open/close and index open/close were the opposite, and the remaining five movements were in the intermediate category. To detect each of these movements with the same power, the weaker activations need a statistical boost (thumb open/close and index open/close), which can be realized through longer scanning (i.e., more runs or longer runs). Our findings with different number of runs showed higher Z-stats with greater number of runs. Hence, our recommendation for a practical protocol is to design the study with different number of runs for different movements depending on their detection power. A “one size does not fit all” approach is suggested. We recommend practical choices later in this section.
ii. Degree of activation appears to depend on the volume of muscle fibers engaged in a particular movement (bigger the muscle, larger/stronger the activation). Consequently, the overlap in activations between movements seemed to depend upon the degree of common muscle fibers employed for both movements (refer to Supplemental **Figures S3.1** to **S3.8**). Although this observation is grounded on quantitative measures of similarity (dice coefficient) presented by us (**Fig. 1d**), the inference on muscle fibers is qualitative and speculative. There is, however, support for this speculation [5]. We found, for instance, that wrist prone/supine had activations more similar to thumb up/down (*D*_*c*_=0.39) than wrist flexion/extension (*D*_*c*_=0.21) (refer to Supplemental **Figures S3.5** and **S3.7**). Hence, identifying wrist prone/supine activation that is different from thumb up/down with high precision would require more statistical power than with wrist flexion/extension. As another example, thumb up/down and rotation (that involve more muscles and tendons in common) resulted in similar activations (no. of activated voxels = 4438 and 6125, respectively), with considerable overlap (*D*_*c*_=0.53); however, thumb open/close had much fewer activated voxels (=1396) with lesser overlap with the other two (mean *D*_*c*_=0.21). Four of the movements had a similar profile with similar number of activated voxels with similar statistical power and having higher dice coefficients (i.e., similarity) – thumb up/down, thumb rotation, digits 3-5, and wrist prone/supine. All these movements involve a degree of common muscles. This observation could either be beneficial or problematic depending on the hypothesis being tested. We could expect it to be generally more challenging to distinguish between regions of activation arising from these movements because of the overlap. Our recommendation for a practical protocol is to design the study with higher power (and more runs) if the aim is to distinguish between movements that recruit similar muscles. If the aim is to study these movements independently, fewer runs would have the power to detect the activations (as few as 3 runs in our study).
iii. The Z-stat generally increased with more runs, but the gains were not proportionally higher with larger number of runs (refer to **Figures 2b** and **2c**). With the left arm, Z-stat increased considerably from 2 to 4 runs (4 vs. 2 runs T-statistic=23.7), but then the increment was smaller from 4 to 6 runs (T=11.1). With the right arm, Z-stat saw a considerable jump from 2 to 4 runs (T=98.8), but then reduced with a greater number of runs (4 vs. 6, T=–35.1). Since this study analyzes data from one healthy participant, confirmatory claims cannot be made without scanning a larger cohort or scanning the same person several times. Nevertheless, our findings suggest that increasing the number of runs does not proportionally increase detection power. Prior work suggests a less-than-linear increase in detection power with scan length or number of runs [26]. There might be several reasons for this, including task performance (boredom, exhaustion), habituation, and handedness. Future studies must further investigate this plateauing effect. Our recommendation for a practical protocol is that there are diminishing returns to increasing the number of runs – that is, adding more runs to the study becomes less justifiable upon a cost-benefit assessment.

As mentioned in the introduction, motor somatotopy is more sophisticated than sensory somatotopy. Prior research has determined that spatial location of activations due to different upper limb movements is variable across individuals, but relative similarity between them is preserved [27]. Referring to activation maps and 3D maps presented in Supplemental Information sections S2 through S5, one might wonder why some upper limb movements resulted in widespread activations outside traditionally considered upper limb areas of M1. Our activations were spatially variable across the span of M1, in addition to having considerable overlap across different movements (as in prior studies [6]). As per the latest research, however, there exists many-to-many mappings between M1 and body movements [5], i.e., for a given movement, various parts of M1 are typically engaged, even regions not typically in the ‘upper limb areas’, and this profile varies across individuals [27]. This supports our observation that M1 lacks a strong somatotopic organization. M1 neurons are tuned to all finger and wrist movements instead of individual movements [28]. In fact, cortical stimulation of M1 simultaneously moves multiple fingers [29]. Other studies support our observation as well; Plow et al. [8] reported similar finger and elbow activations with a 15-min fMRI motor task.

Based upon the insight obtained from our results, we propose a practical scanning protocol for three cases – scan sessions with a duration of 60 min, 75 min, and 90 min. Tasks with larger, more robust activations (thumb up/down and rotation, digits 3-5, wrist prone/supine, wrist flexion/extension, and elbow) showed remarkably visible consistency of activation for 3, 5 and 7 runs cases (see Supplemental Information sections S4 and S5). Tasks with weaker activations (thumb open/close, index finger) were more inconsistent across runs, and in some cases hardly showed activation with 2 runs. We suggest that at least 3 runs of scanning (3 blocks or 2 mins per movement) is required to robustly estimate activation with thumb open/close and index open/close movements. Note that each task block was 41.25 sec long in our study, and each run was 8 blocks long (5 min 30 sec). As mentioned earlier, ‘one size does not fit all’, so we propose different number of runs for different movements depending upon their detection power. We will assume that a 6-min anatomical run and a 10-min resting-state fMRI run will be part of the scanning protocol.

a. 60-min scan session:
  - Runtime for 6 movements with ‘good’ activations (2 runs): 2 left runs + 2 right runs = 4 × 4 min 8 sec = 16 min 32 sec
  - Runtime for 2 movements with ‘weaker’ activations (3 runs): 3 left runs + 3 right runs = 6 × 1 min 23 sec = 8 min 15 sec
  - Here, the first two runs could have all the 8 movements’ task blocks and the last run could have task blocks of only the two movements with weaker activations.
  - Total task fMRI time = 24 min 47 sec
  - Total scan time ≈ 10 (resting-state fMRI) + 25 (task fMRI) + 6 (anatomical) + 9 (breaks, interaction with participant, shimming) + 10 (participant set-up in scanner [loading], and removal of participant and cleaning of equipment [unloading]) ≈ 60 mins
b. 75-min scan session:
  - Runtime for 6 movements with ‘good’ activations (3 runs): 3 left runs + 3 right runs = 6 × 4 min 8 sec = 24 min 45 sec
  - Runtime for 2 movements with ‘weaker’ activations (5 runs): 5 left runs + 5 right runs = 10 × 1 min 23 sec = 13 min 45 sec
  - Total task fMRI time = 38 min 30 sec
  - Total scan time ≈ 10 (resting-state fMRI) + 38.5 (task fMRI) + 6 (anatomical) + 10.5 (breaks, interaction, shimming) + 10 (loading and unloading) ≈ 75 mins
c. 90-min scan session:
  - Runtime for 6 movements with ‘good’ activations (4 runs): 4 left runs + 4 right runs = 8 × 4 min 8 sec = 33 min 0 sec
  - Runtime for 2 movements with ‘weaker’ activations (7 runs): 7 left runs + 7 right runs = 14 × 1 min 23 sec = 19 min 15 sec
  - Total task fMRI time = 52 min 15 sec
  - Total scan time ≈ 10 (resting-state fMRI) + 52.25 (task fMRI) + 6 (anatomical) + 11.75 (breaks, interaction, shimming) + 10 (loading and unloading) ≈ 90 mins

These proposed protocols were, of course, tuned to our initial protocol with an on-off block design, 20.625-sec blocks and 5.5-min runs, and assuming all eight movements are to be included. Modifications could be made by researchers depending upon their requirements. Finally, the following points present the study limitations and suggest the scope for future work:

i. This is a pilot study; our conclusions were based upon extended scanning of one participant. Idiosyncrasies of the participant, degree of handedness, task performance and other such factors are more pronounced compared to population studies. Nonetheless, we do not expect the broad contours of our observations to differ substantially across healthy volunteers, but researchers replicating these results could expect variations in the minutiae; for instance, a different participant might need more runs to show stabler activations (yet we expect this to be consistent across movements for a given subject).
ii. Our results are specific to 7T imaging. We expect the conclusions to be broadly similar at 3T, but the numbers would, of course, be different due to differences in image signal-to-noise ratio [30] and functional contrast-to-noise ratio [31]. We therefore expect 7T imaging to provide at least a doubling of functional contrast compared to 3T, which can be traded for shorter scan times while retaining statistical power. As such, conducting similar experiments at 3T would require longer scans to produce similar results, or fewer tasks could be performed with an expectation of similar sensitivity within the same constraints of a 60- to 90-min session.
iii. The descending white matter tracts from M1 do not reach muscle fibers directly. They innervate neurons in the ventral horn of the spinal cord. Cervical spinal cord levels C5 through C8 are involved in upper limb movements [32], and functional neuroimaging of upper limb movements is not complete without the study of the cord. Fortunately, this is possible due to recent advancements in spinal cord functional imaging (especially at 7T) [33]. Future studies must consider the possibility of including spinal cord imaging in their acquisition protocol.
iv. Due to the similarities between the upper limb tasks being performed, we did not have a tight control over motor performance, which could have led to less robust activations due to co-occurrence of movements and task performance [34].
v. Although both muscle fibers and movement patterns find representations in M1 [35], fMRI is largely sensitive to the muscle fibers engaged rather than the pattern in which those movements were made [27]. As such, our study has common limitations of fMRI including limited spatial/temporal resolution, complication of negative BOLD responses in M1 [36], difficulty measuring inhibitory neurons [37], and so on. However, our primary aim was to propose a practical protocol for fMRI scanning at 7T, which we believe has been accomplished.

## 5. Conclusions

In this study, we found that increasing the scan length has diminishing returns for identifying M1/S1 activations with upper limb movements. Larger muscle movements were associated with larger/stronger activations; two movements were identified as having low activation and six movements as medium/high activation. A bare minimum of three runs in the former case and two runs in the latter were found to be necessary to identify reasonable activations (one on/off block of 41.25 seconds per movement in each run). We proposed a ‘*one size does not fit all*’ practical protocol wherein more runs of the low activation movements are performed to boost their statistical power relative to medium/high activation movements. Researchers can adopt and modify the proposed protocol depending on their hypothesis and experimental setup.

## CRediT (Contributor Roles Taxonomy) – author contributions

**D Rangaprakash:** Conceptualization, Methodology, Software, Data Analysis, Investigation, Visualization, Writing - Original Draft. **Olivia E Rowe**: Project Administration, Methodology, Investigation, Writing - Reviewing and Editing. **Hyungeun Song**: Data Acquisition, Investigation, Writing - Reviewing and Editing. **Samantha Gutierrez-Arango**: Data Acquisition, Investigation, Writing - Reviewing and Editing. **Michael F Fernandez**: Data Acquisition, Investigation, Writing - Reviewing and Editing. **Erica A Israel**: Data Acquisition, Investigation, Writing - Reviewing and Editing. **Hugh M Herr**: Funding Acquisition, Resources, Project Administration, Investigation, Writing - Reviewing and Editing. **Robert L Barry:** Funding Acquisition, Resources, Project Administration, Conceptualization, Methodology, Data Acquisition, Investigation, Writing - Reviewing and Editing, Supervision.

## Acknowledgments

Imaging was performed at the Athinoula A. Martinos Center for Biomedical Imaging at the Massachusetts General Hospital using resources provided by the Center for Functional Neuroimaging Technologies (P41EB015896) and the Center for Mesoscale Mapping (P41EB030006), Biotechnology Resource Grants supported by the National Institute of Biomedical Imaging and Bioengineering, National Institutes of Health (NIH). The NIH also provided support through a shared instrumentation grant S10OD023637 and grant R01EB027779 (R.L.B.). This research was also supported in part by the MGH/HST Athinoula A. Martinos Center for Biomedical Imaging. The content is solely the responsibility of the authors and does not necessarily represent the official views of the NIH.

## Data availability statement

All related study data including the thresholded Z-statistic NIfTI images have been made publicly available through Mendeley Data [25]. Any study data not available through this source could be obtained from the corresponding author upon reasonable request.

