## Supplemental Information for "Practical scan-length considerations for mapping upper limb movements to the somatosensory/motor cortex at 7T"

This document provides additional illustrations to supplement the information provided in the main text. The information provided here does not alter the conclusions in any way (i.e., the main text is self-sufficient), but it augments the information in main text by providing visuals that help in understanding the results better. This document renders more detailed information for replication and future studies. All the figures presented here, as well as all the thresholded Z-statistic NIfTI images are publicly available (DOI: 10.17632/ggyy848pxj.1), using which readers could regenerate and compare any of these maps in a software of their choice. All images must be viewed with the radiological convention.

While interpreting these maps, it must be noted that group-level (second level) maps are generally more robust and less noisy compared to subject-level (first level) maps. We performed a subject-level analysis, and our figures show first level maps. As such, it will appear noisier than figures seen commonly in fMRI activation papers.

Section S1 provides screenshots of fMRI pre-processing and activation analysis in FSL (contains Figures S1.1 through S1.4). Section S2 provides activation maps for different upper limb movements generated by FSL (contains Figures S2.1 through S2.11). Section S3 provides 3D overlay images for different upper limb movements generated using Mango (contains Figures S3.1 through S3.8). Section S4 provides activation maps for different number of runs generated by FSL (contains Figures S4.1 through S4.8). Section S5 provides 3D overlay images for different number of runs generated using Mango (contains Figures S5.1 through S5.8).

### S1. FMRI pre-processing and activation analysis screenshots

Pre-processing of fMRI data and activation analysis were performed in FSL v.6.0.3. The procedures were described in the main text. Here, we present screenshots of the setup in FSL FEAT, for the benefit of the readers (Figures S1.1 and S1.2). Additionally, the GLM design matrices for the cases of 2 runs through 8 runs are presented in Figures S1.3 and S1.4.

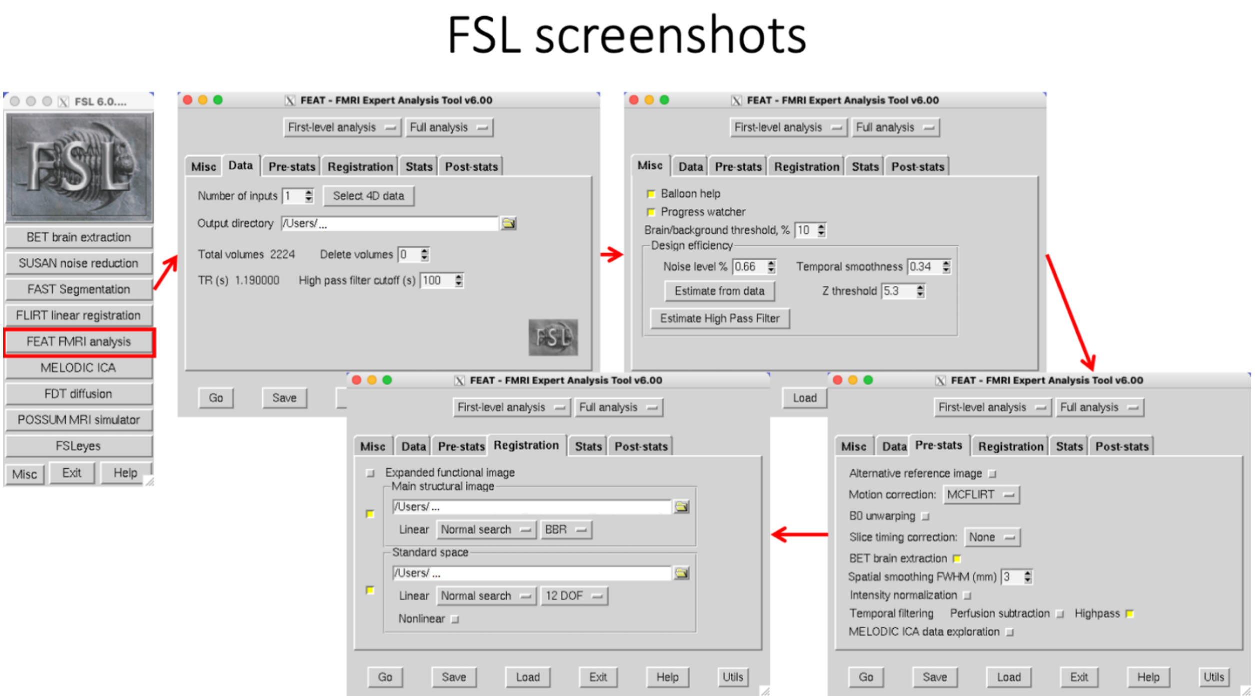


**Figure S1.1.** *Pre-processing setup in FSL FEAT*

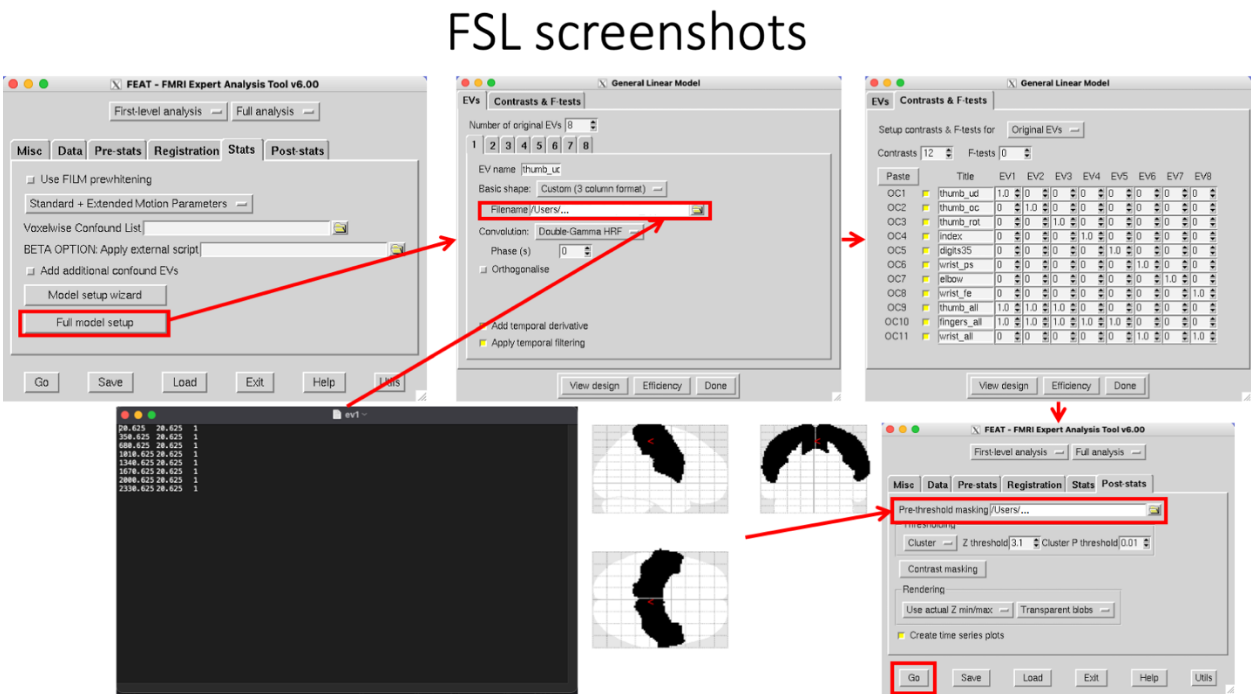


**Figure S1.2.** *GLM activation analysis setup in FSL FEAT. Figure also shows an example explanatory variable (EV) text file, and the Harvard-Oxford atlas' primary somatosensory/motor cortex mask used for pre-threshold contrast masking (illustration using SPM12).*

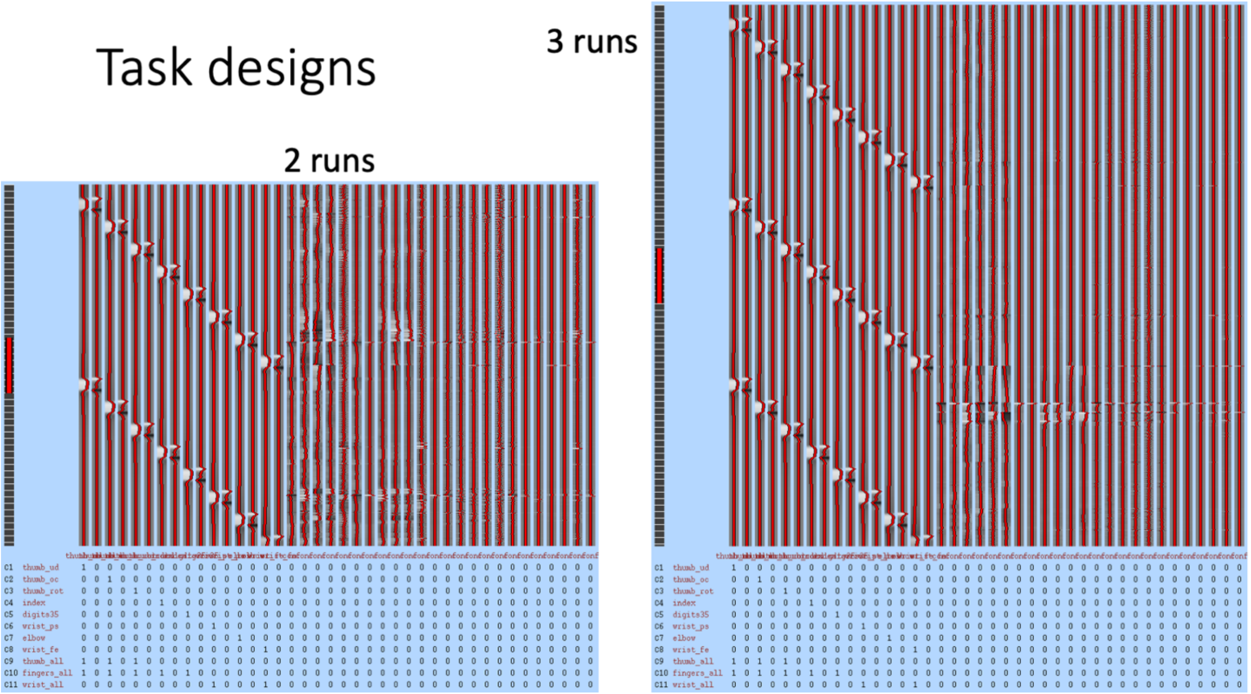


**Figure S1.3.** *GLM design matrices for the cases of 2 runs and 3 runs. Each EV (convolved with a canonical HRF) and its temporal derivative appear as the first 16 columns (corresponding to the 8 EVs). The remaining columns correspond to the 6 basic motion parameters (3 translations, 3 rotations) and 18 extended motion parameters (6 temporal derivatives and 12 squares of them all). These images were self-generated by FSL upon execution (file 'design.png' in the .feat directory).*

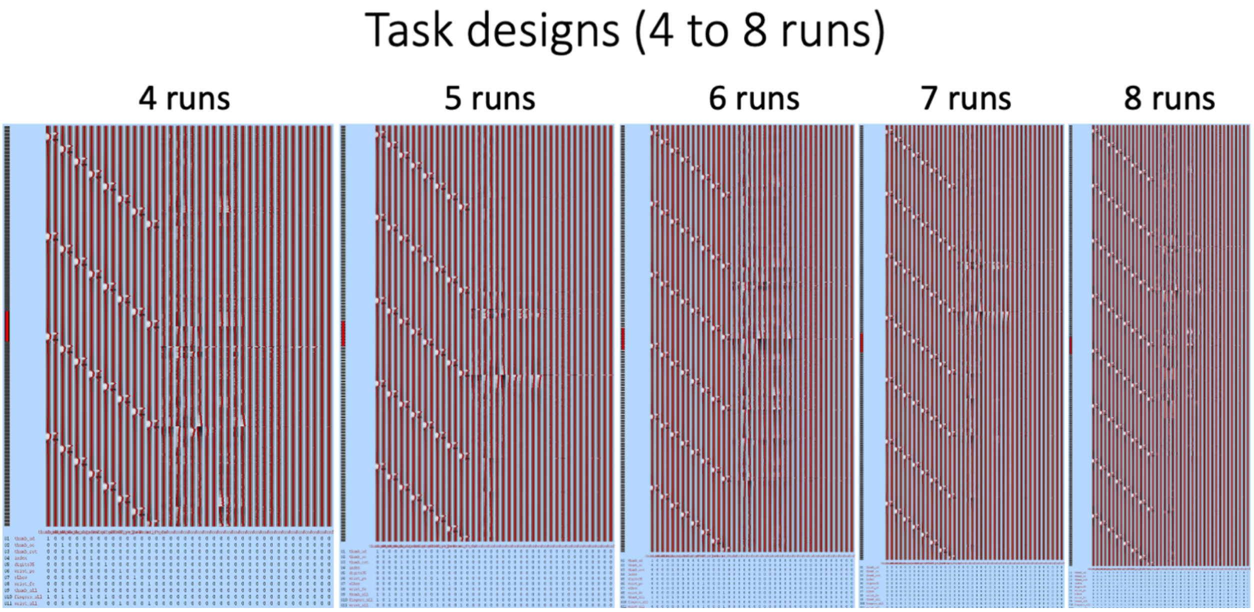


**Figure S1.4.** *GLM design matrices for the cases of 4 runs through 8 runs.*

### S2. Activation maps for different upper limb movements

GLM analysis in FSL outputs the axially sliced images of first-level activations (thresholded Z-stats), sliced along the *z*-direction in the participant's native space (one image per acquisition slice) (files '*rendered_thresh_zstat1.png*' and so on in the .feat folder). Here we present these images for all movements with both left and right arms (for the case of all 8 runs) (original images were cropped consistently using a MATLAB script). In each figure, the axial slices from top to bottom correspond to slices n=46 through n=85 in the participant's native functional space (n=1 is the bottommost slice). In the MNI space, this corresponds to *z*=0 through *z*=60 (in increments of 1.5mm, the voxel height).

Apart from the conclusions drawn in the main text, a few observations were as follows. The distinction between *all fingers* and *all wrist* cases was easily observable. *All fingers* and *all thumb* cases were also are similar, with *all fingers* having a few more activated voxels as expected. See Figure S1.2 for how *all thumb*, *all fingers* and *all wrist* contrasts were generated. Consistent patterns were observed with left and right arm movements in the corresponding hemispheres.

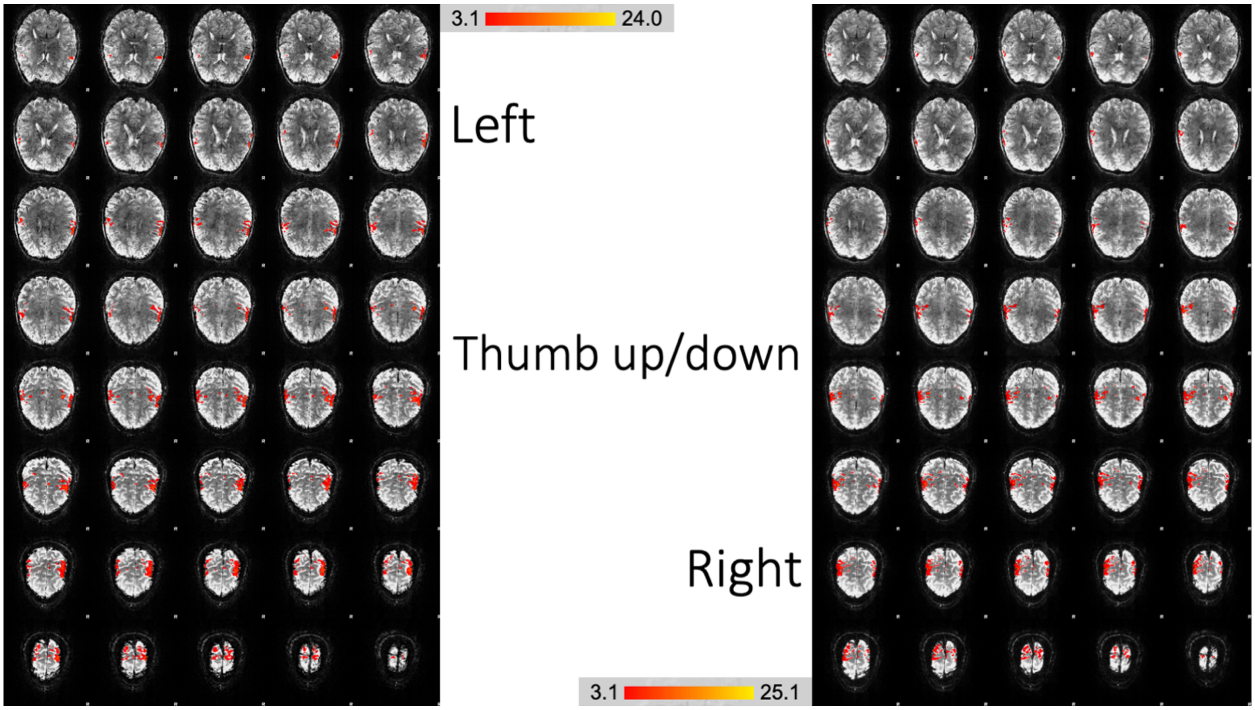


**Figure S2.1.** *BOLD activations due to thumb up/down movement*

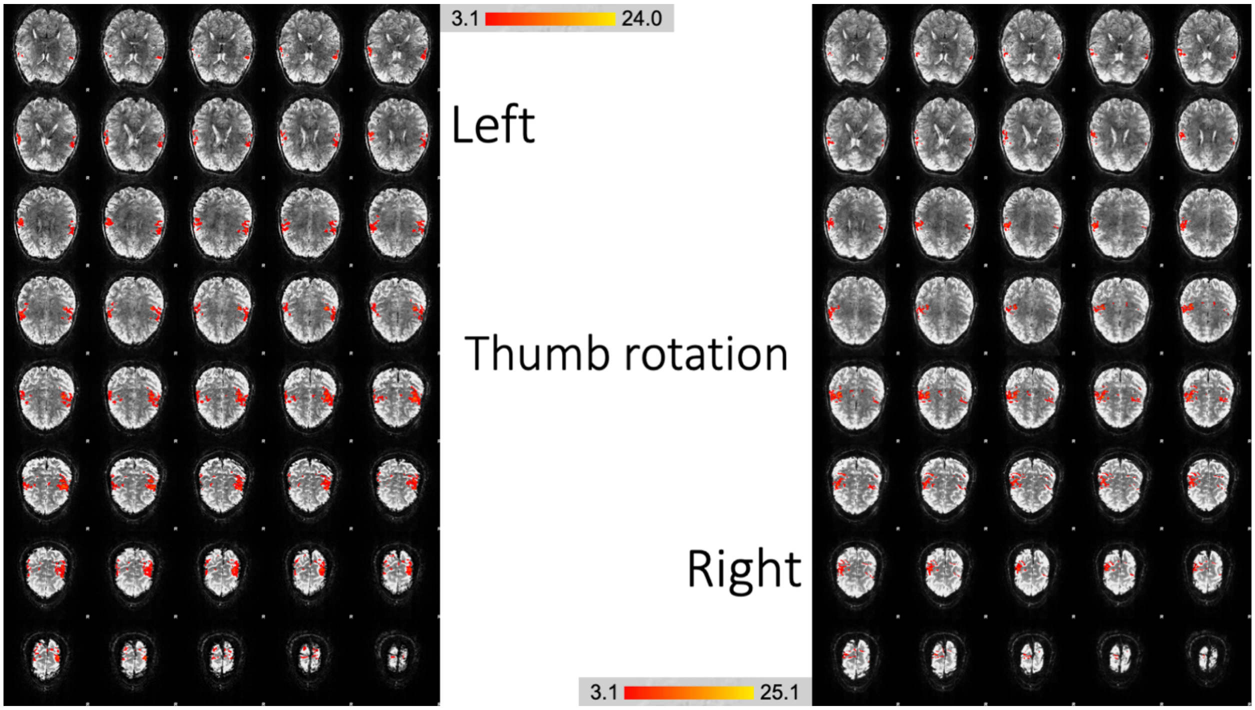


**Figure S2.2.** *BOLD activations due to thumb rotation*

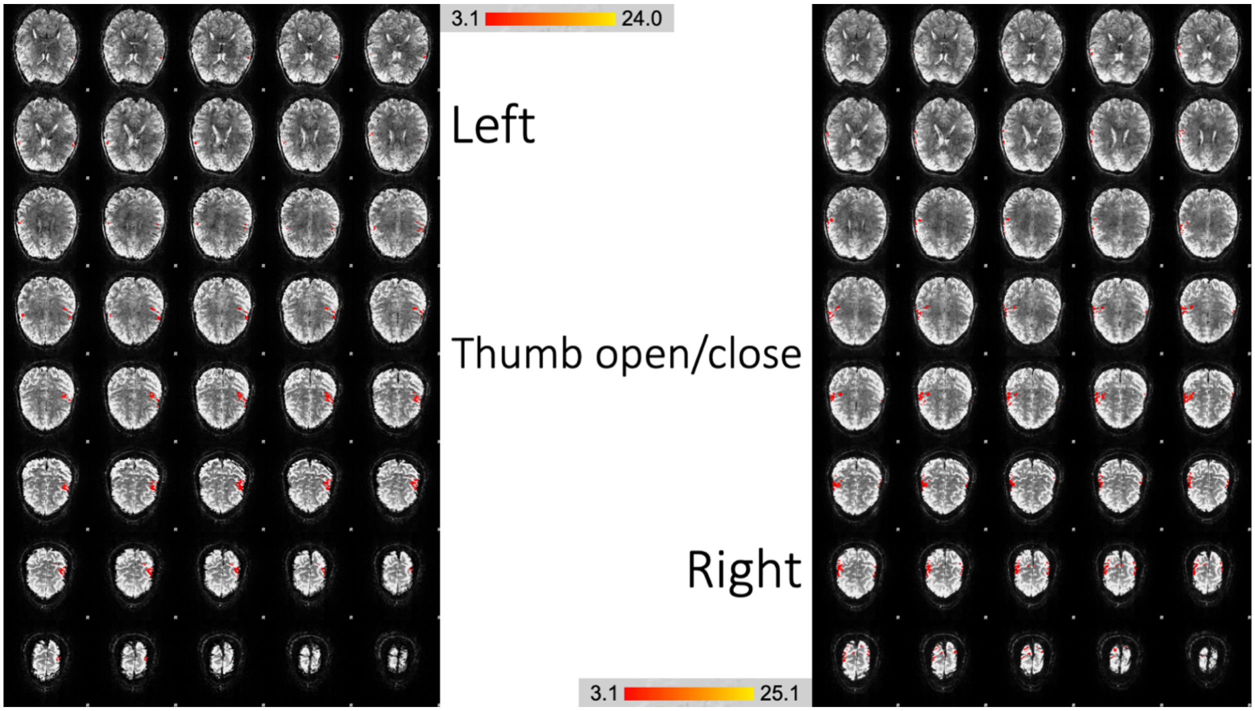


**Figure S2.3.** *BOLD activations due to thumb open/close movement*

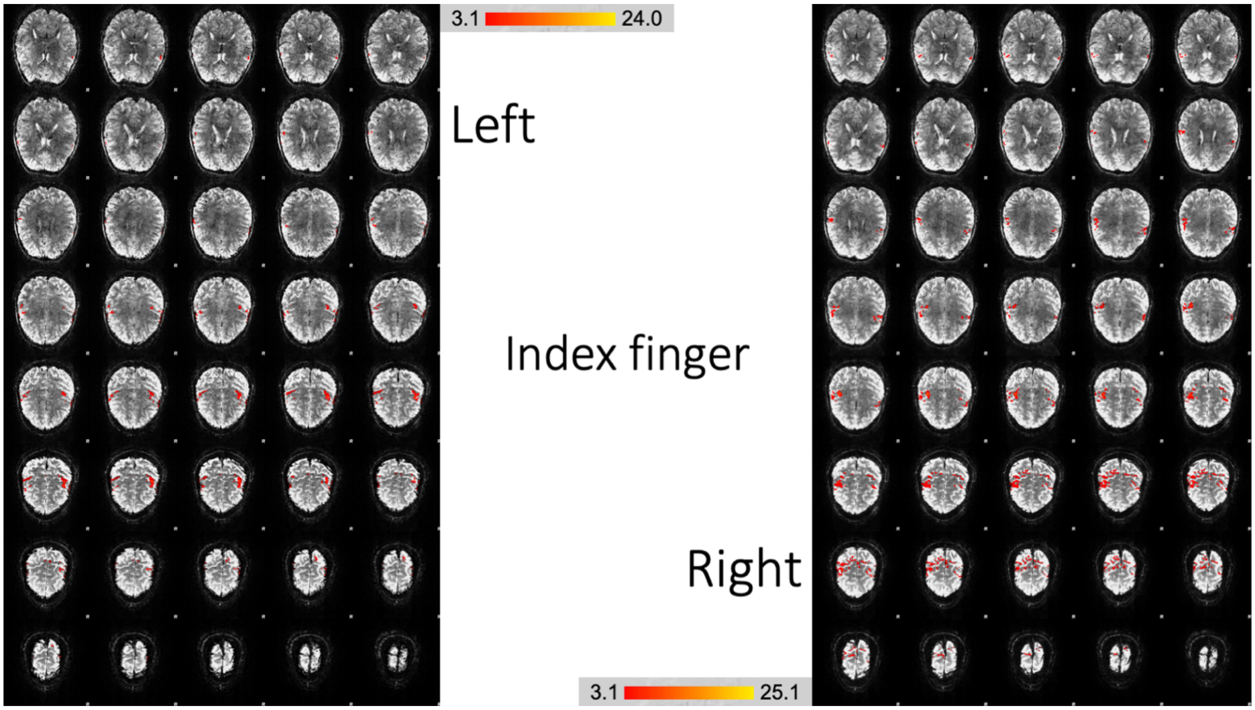


**Figure S2.4.** *BOLD activations due to index finger open/close movement*

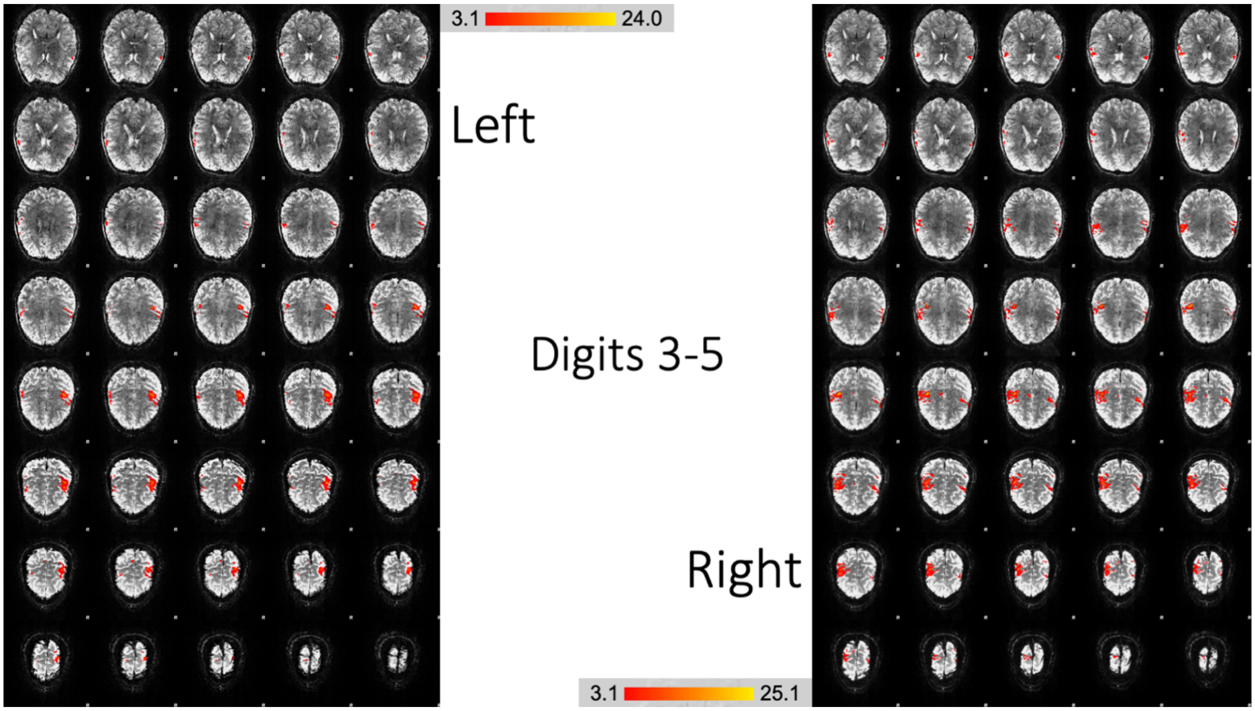


**Figure S2.5.** *BOLD activations due to digits 3 to 5 open/close movement*

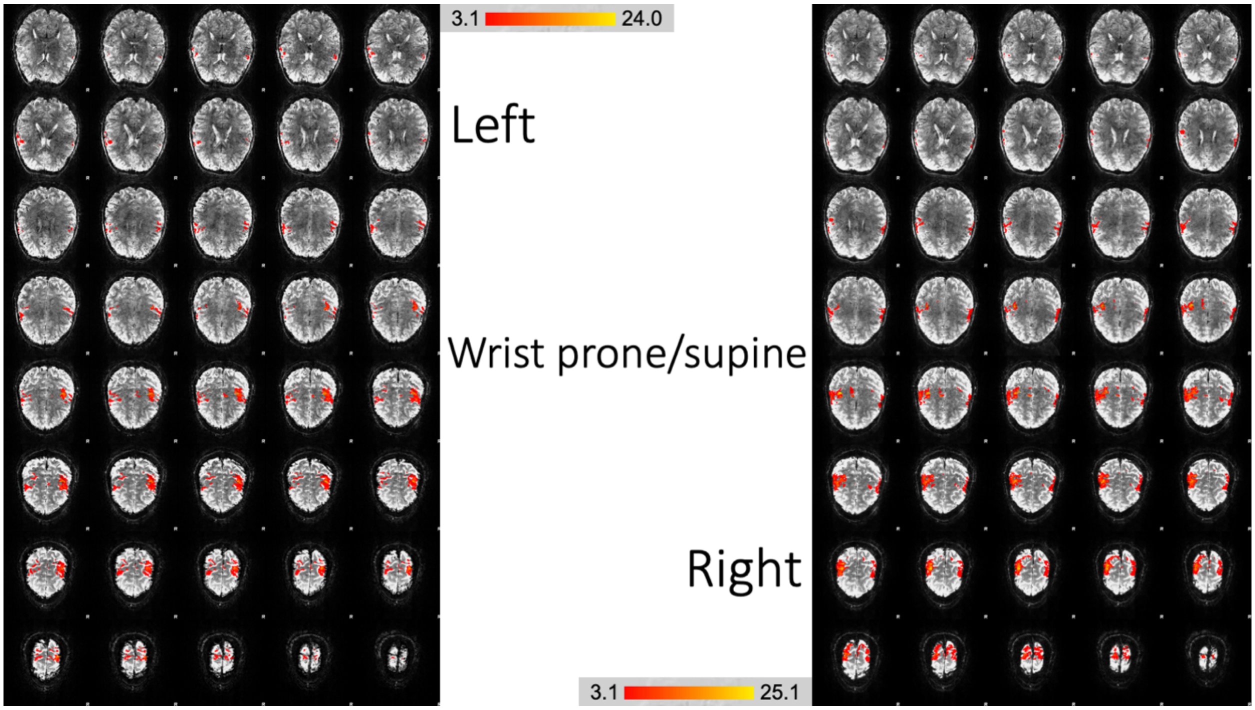


**Figure S2.6.** *BOLD activations due to wrist prone/supine movement*

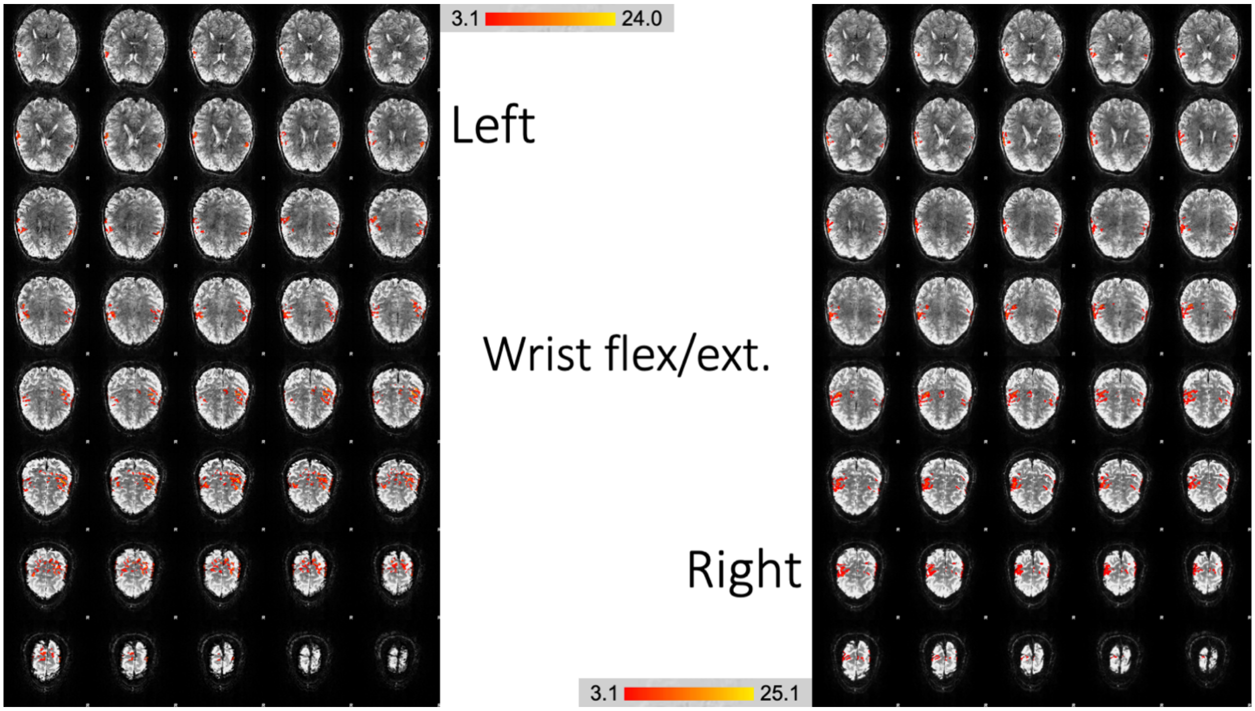


**Figure S2.7.** *BOLD activations due to wrist flexion/extension movement*

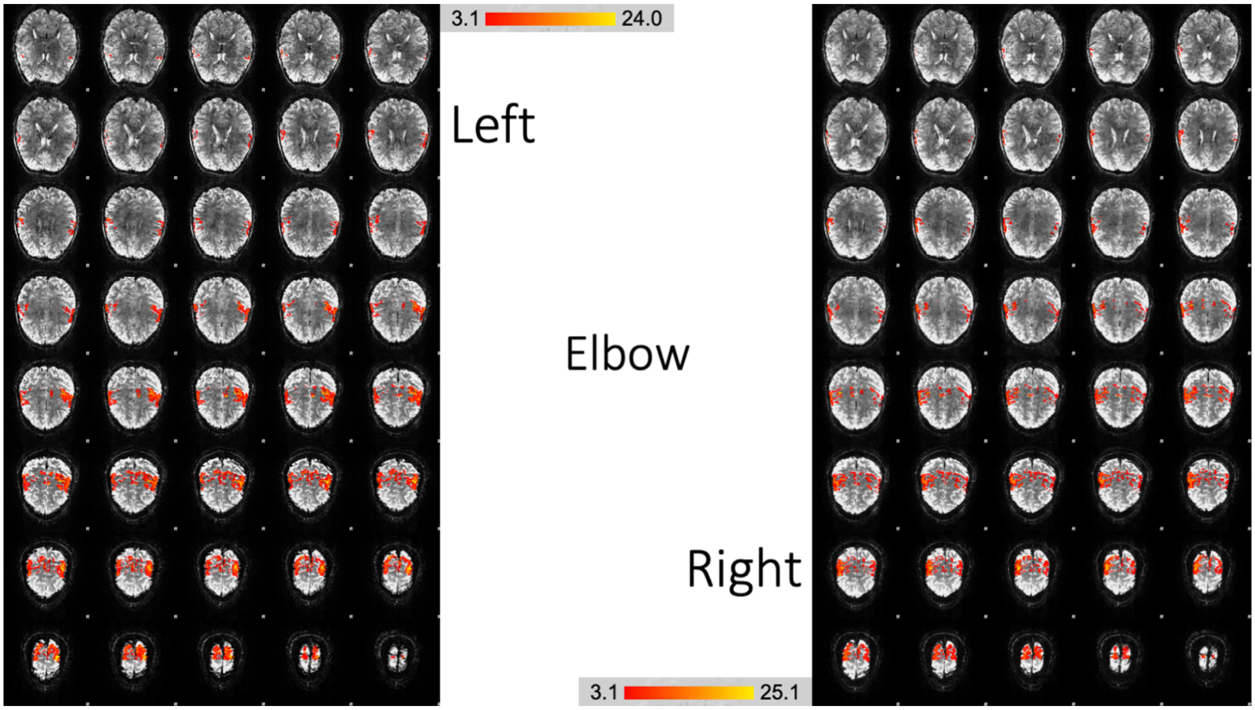


**Figure S2.8.** *BOLD activations due to elbow open/close movement*

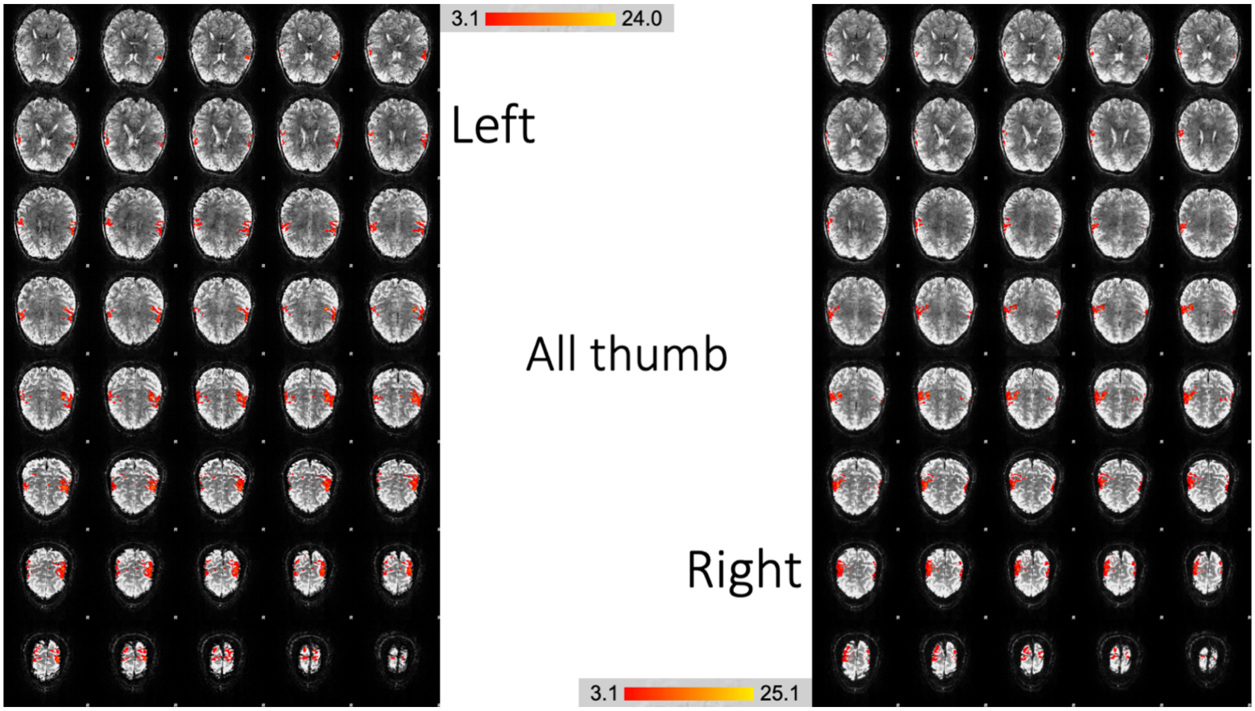


**Figure S2.9.** *BOLD activations due to all the thumb movements taken together*

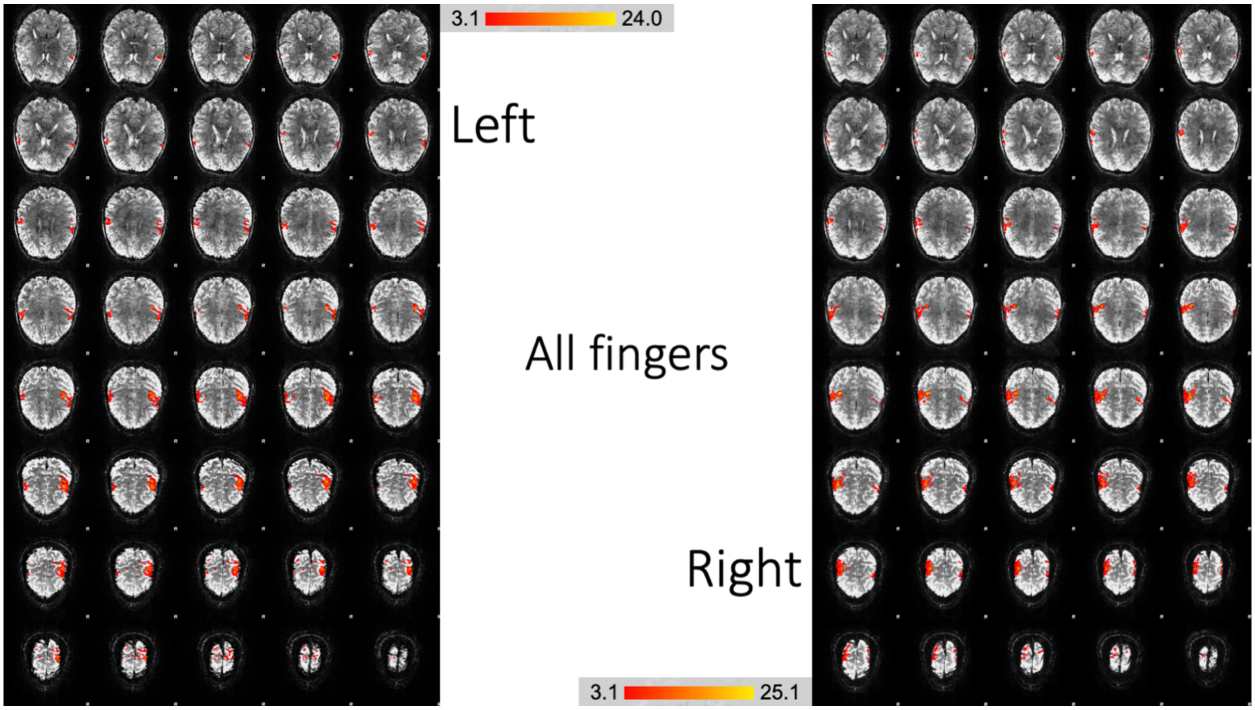


**Figure S2.10.** *BOLD activations due to all the five finger movements taken together*

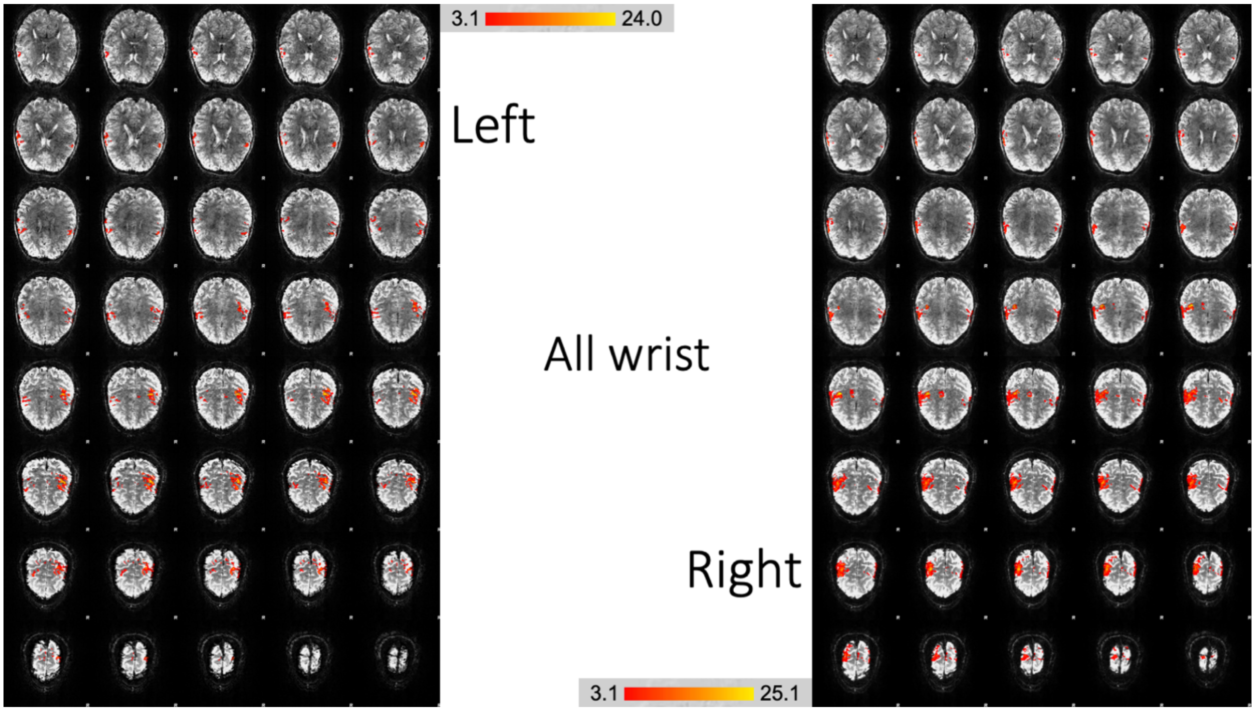


**Figure S2.11.** *BOLD activations due to all the wrist movements taken together*

### S3. 3D overlay images for different upper limb movements

For additional visual insight on how similar or different the activations were with different limb movements, we provide snapshots of activations overlaid on a 3D brain (8 runs case). These 3D overlays were generated using Mango (http://ric.uthscsa.edu/mango/) as follows. The Z-stat images were overlaid on the mean functional image, with an overlay color of red for the first overlay and green/blue for subsequent overlays. This rendered all Z values in the same color (deliberately chosen for better viewing and simplicity). The transparency of the first (red) overlay was set to 0.1 and that of subsequently overlays were set to 0.4 for green and 0.5 for blue. The 3D image was then generated using the 'Build Surface' feature under 'Image' in the menu (default settings, threshold=5000). Readers could regenerate these using the provided Z-stat NIfTI images.

As mentioned in the main text, it must be noted that fMRI in the primary somatosensory/motor cortex is mostly sensitive to the volume and location of moving muscle fibers, and not to the sequence or pattern of movements. This also pertains to the limitations of measuring brain function with fMRI, such as its limited temporal resolution and spatial specificity. This is also pronounced with limited scan duration (cannot put someone through several hours of scanning at once in a practical protocol) and task performance issues with increasing scan duration. Overlap in activations across different movements must be interpreted in this light.

For instance, compared to the thumb open/close movement, the thumb up/down and rotation movements involve more muscles (including the palm), some of which are common between the movements. In our data, thumb up/down and rotation resulted in similar activations (no. of activated voxels = 4438 and 6125, respectively), with considerable overlap (dice coefficient, $D_c$=0.53). Much fewer activated voxels (no. of activated voxels = 1396) were noticed with thumb open/close, which also exhibited lesser overlap with the other two (mean $D_c$=0.21).

Index finger and thumb open/close exhibited similar level of activation, and their activation locations were also distinguishable. Digits 3-5 showed overlapping activations with thumb up/down and rotation (all these movements involve palm muscles). Fingers also shared similarity with wrist movements, perhaps because both involve muscles of the palm. Wrist activations shared similarities with thumb up/down and rotation, perhaps because both depend on similar muscle groups for motion. Elbow showed the largest amount of activation; it involves larger number of muscle groups. Elbow also showed notable activation in the contralateral motor cortex, perhaps because muscles from the other part of the chest were also engaged.

Overall, these observations mimicked those presented in the main text.

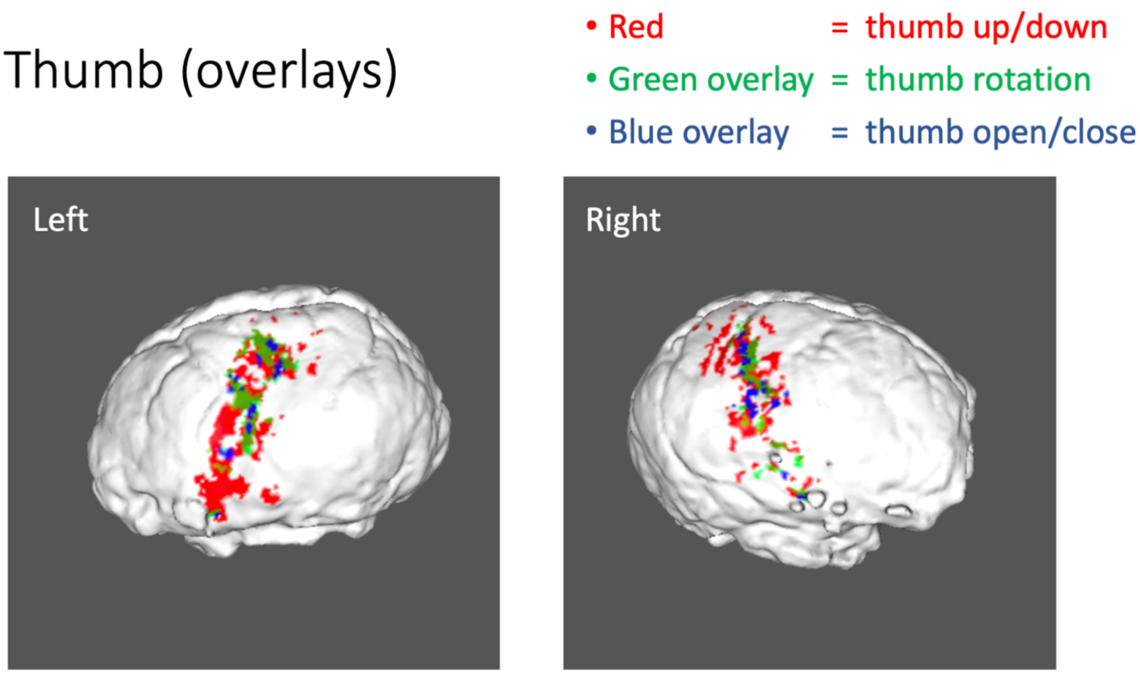


**Figure S3.1.** *3D overlays showing BOLD activations due to different thumb movements. Thumb up/down and rotation activations were similar (mean dice coefficient, $D_c$=0.51). Thumb open/close was less similar to the other two ($D_c$=0.21).*

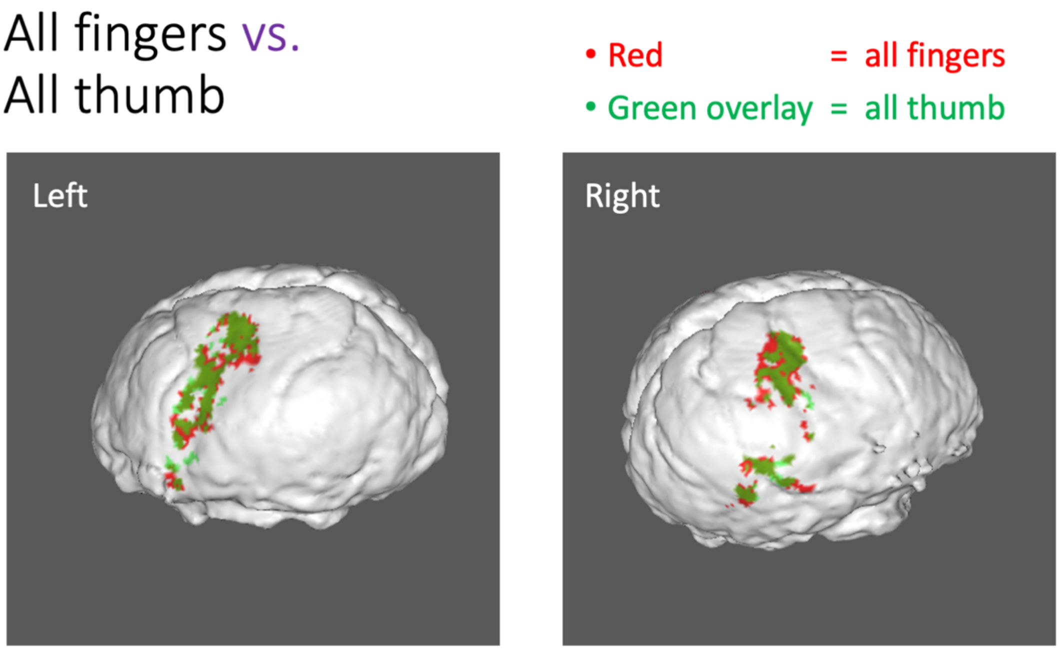


**Figure S3.2.** *3D overlays comparing 'all fingers' (including the thumb) and 'all thumb'. Robust activations with high similarity were noted ($D_c$=0.79).*

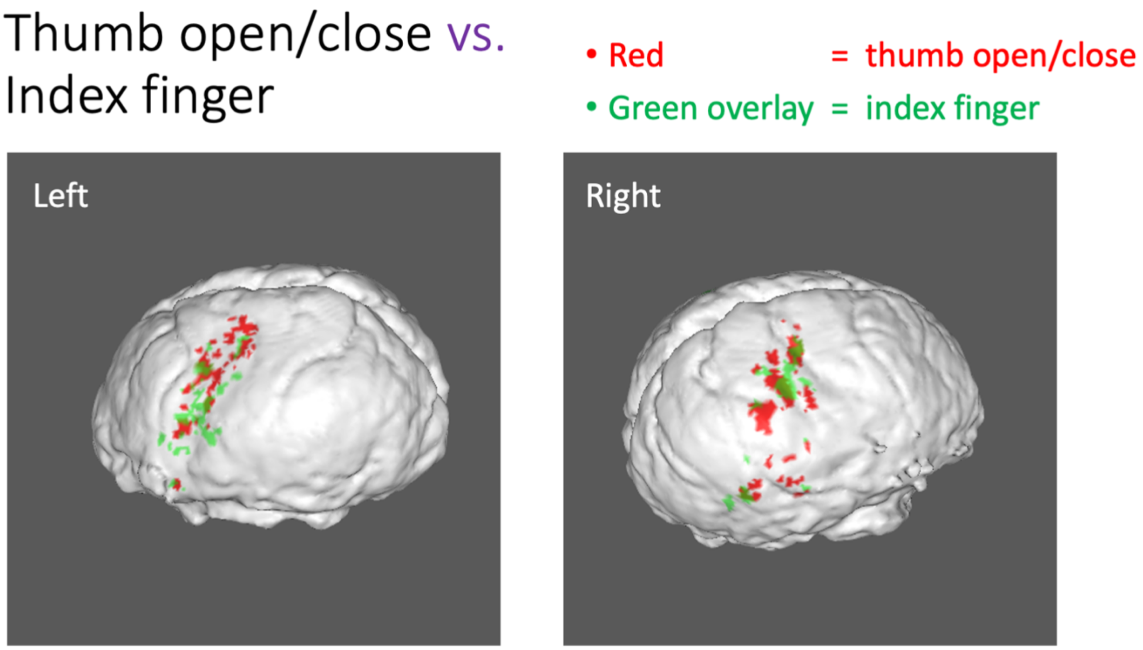


**Figure S3.3.** *3D overlays comparing thumb open/close and index finger open/close activations. Activations were weaker, more scattered and less similar ($D_c$=0.19).*

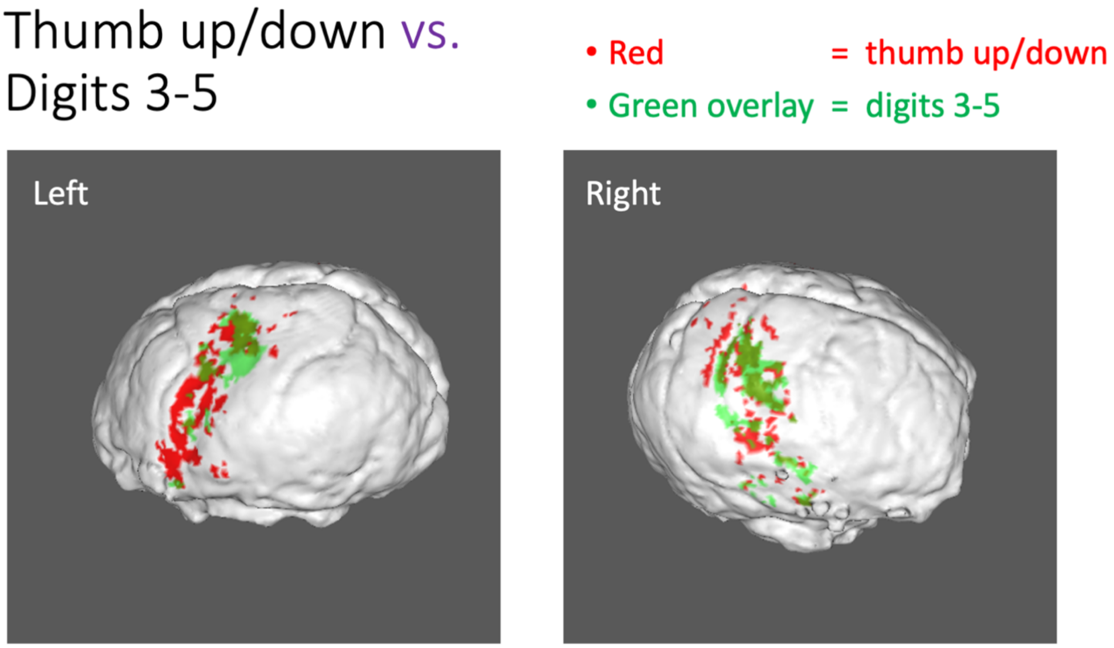


**Figure S3.4.** *3D overlays comparing thumb up/down and digits 3-5 activations with moderate similarity ($D_c$=0.38).*

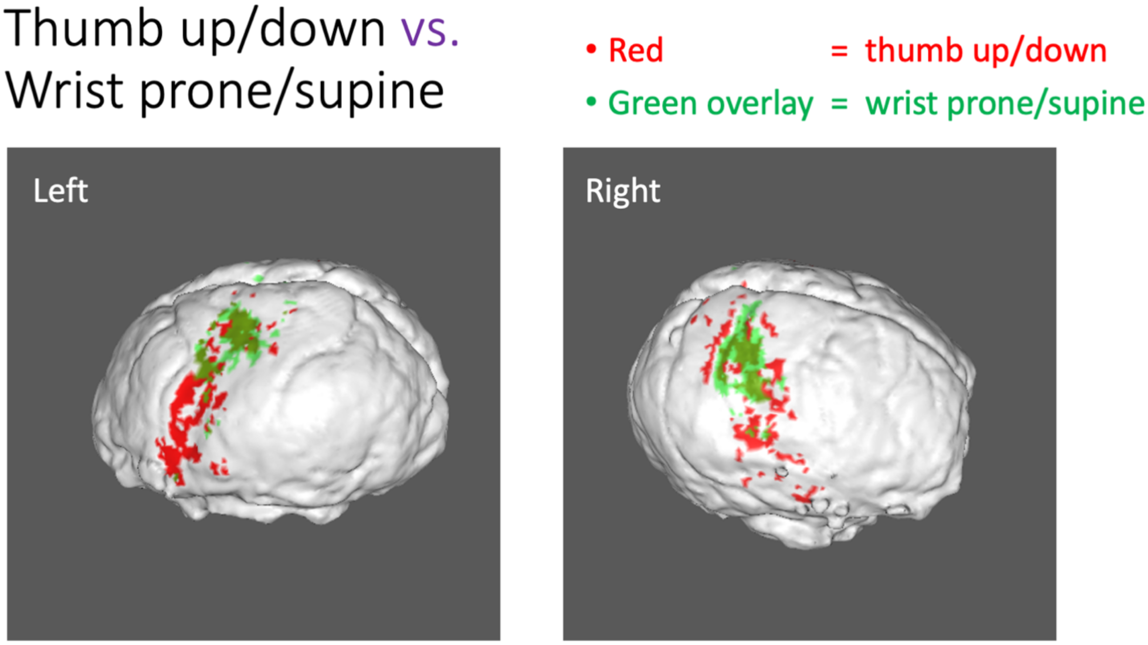


**Figure S3.5.** *3D overlays comparing thumb up/down and wrist prone/supine activations with moderate similarity ($D_c$=0.39).*

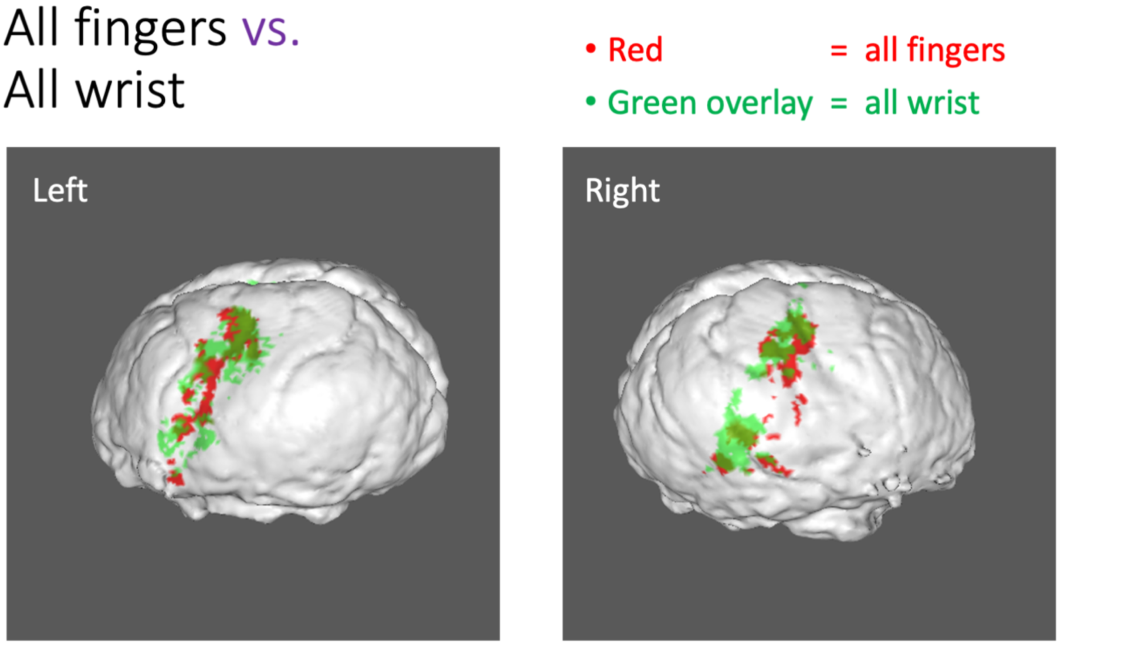


**Figure S3.6.** *3D overlays comparing 'all fingers' and 'all wrist' activations with good similarity ($D_c$=0.51).*

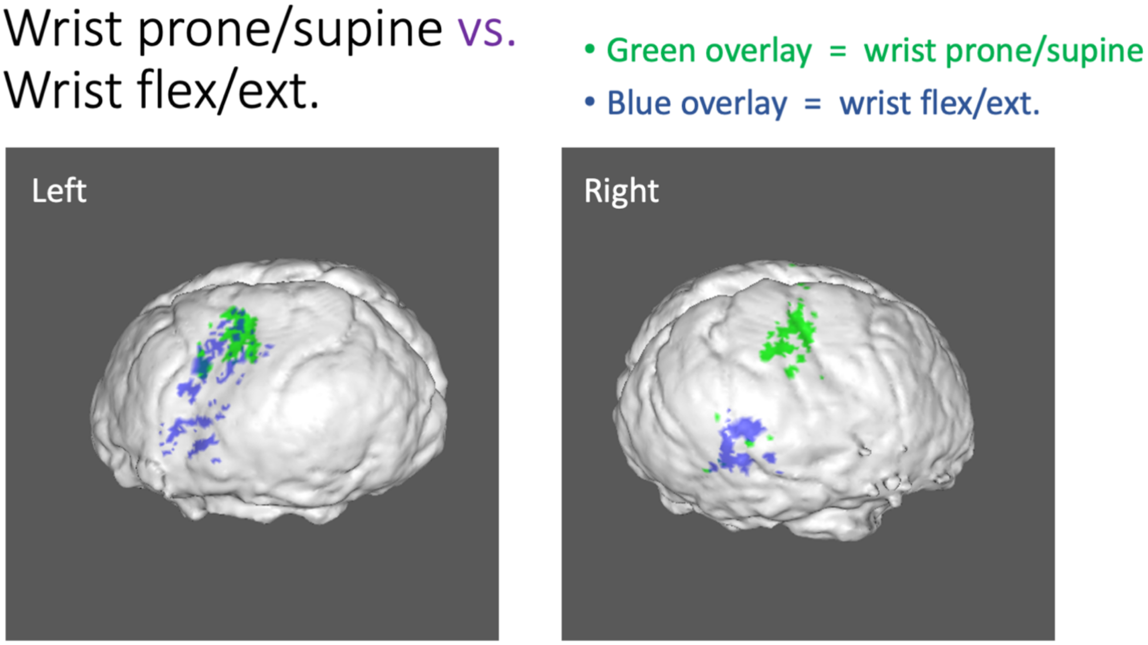


**Figure S3.7.** *3D overlays comparing wrist prone/supine and flexion/extension activations with poor similarity ($D_c$=0.21).*

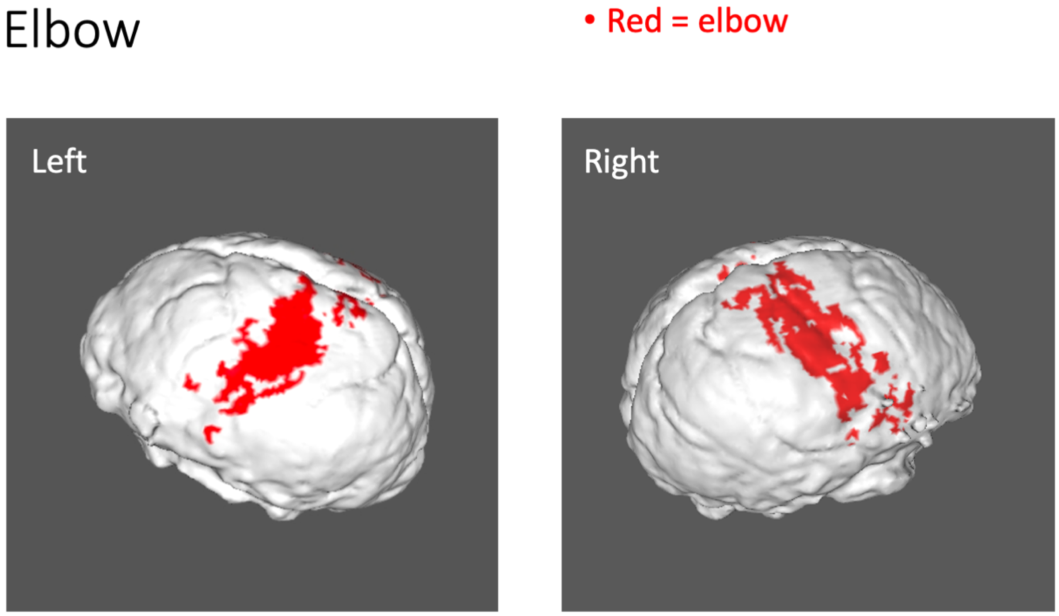


**Figure S3.8.** *3D overlays showing BOLD activations due to elbow movements.*

### S4. Activation maps for different number of runs

Here we present FSL's first-level activation images for the case of different number of runs, with both left and right arms and for all limb movements. For brevity, we present only for the cases of 7 runs, 5 runs, 3 runs and 2 runs (Figures S2.1 through S2.8 presented for the case of 8 runs). All figures (all runs) as well as corresponding NIfTI images are publicly available (DOI: 10.17632/ggyy848pxj.1). In each figure, the axial slices from top to bottom correspond to slices $z$=0 through $z$=60 in the MNI space.

Tasks with larger activations (thumb up/down and rotation, digits 3-5, wrist prone/supine and elbow) showed remarkably visible consistency of activation for 3, 5 and 7 runs cases. Tasks with weaker activations (thumb open/close, index finger) were more inconsistent across runs. Some of these movements hardly showed activation with 2 runs. Whether with 3 or higher runs, these limb movements had weaker detection power. However, it was possible to visually distinguish between tasks even in this case with just 3 runs. Overall, although identifying activations with these tasks was challenging with 3 runs, it was still sufficient to distinguish between limb movements visually.

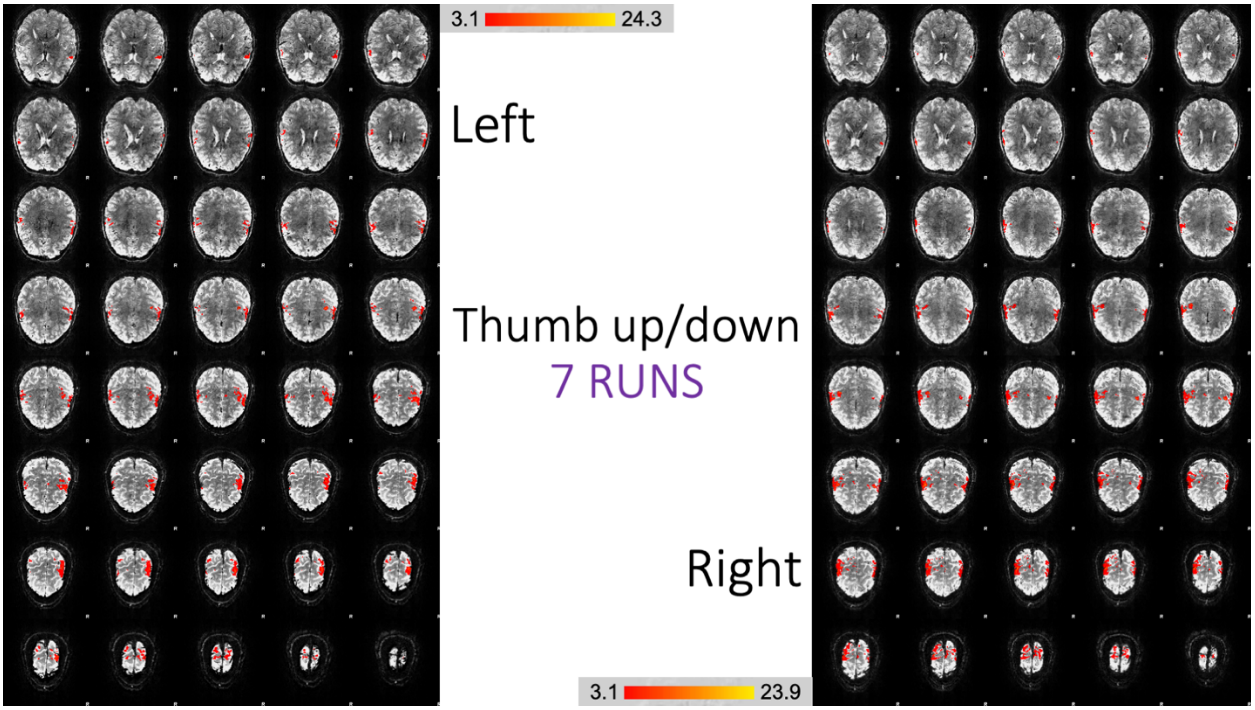
3.1
24.3
Left
Thumb up/down
7 RUNS
Right
3.1
23.9

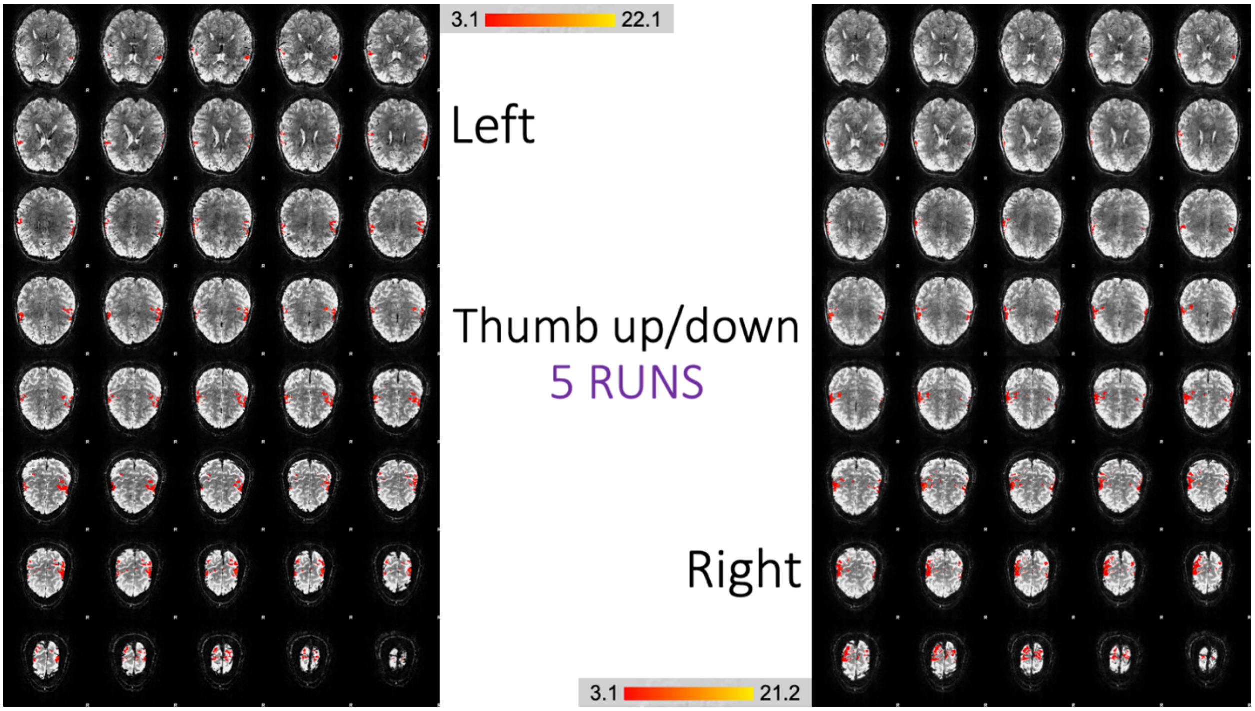
3.1
22.1
Left
Thumb up/down
5 RUNS
Right
3.1
21.2

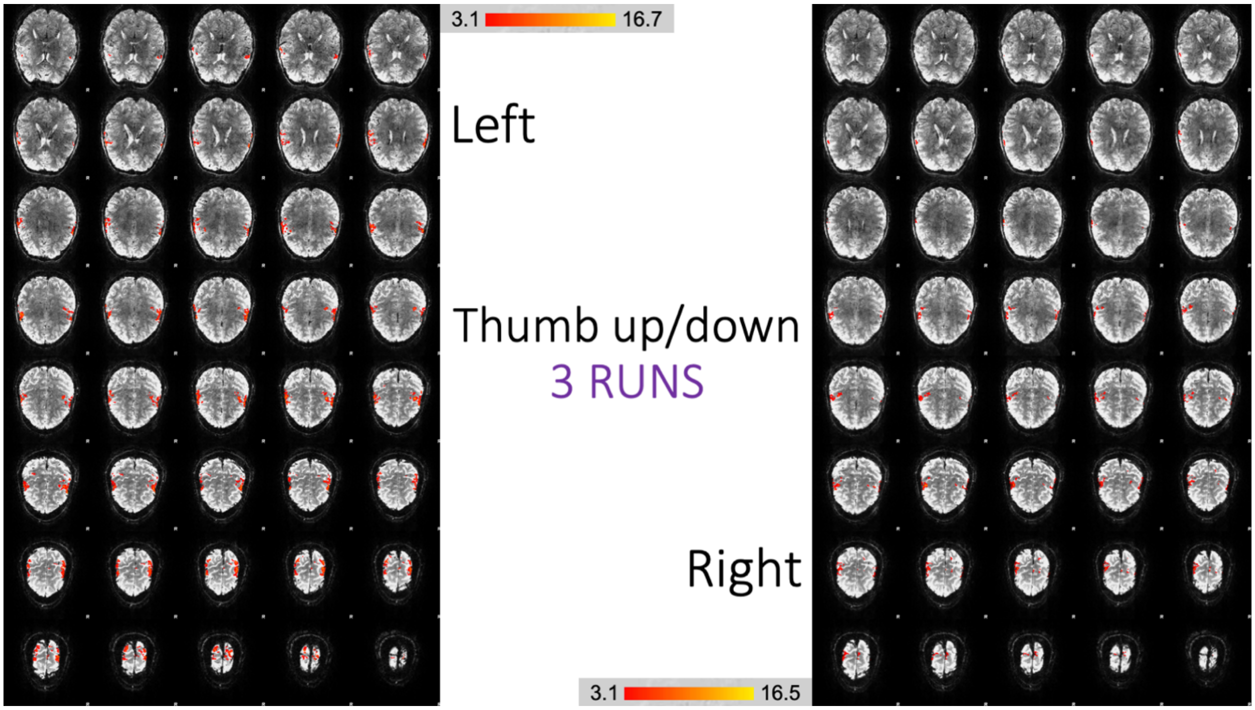


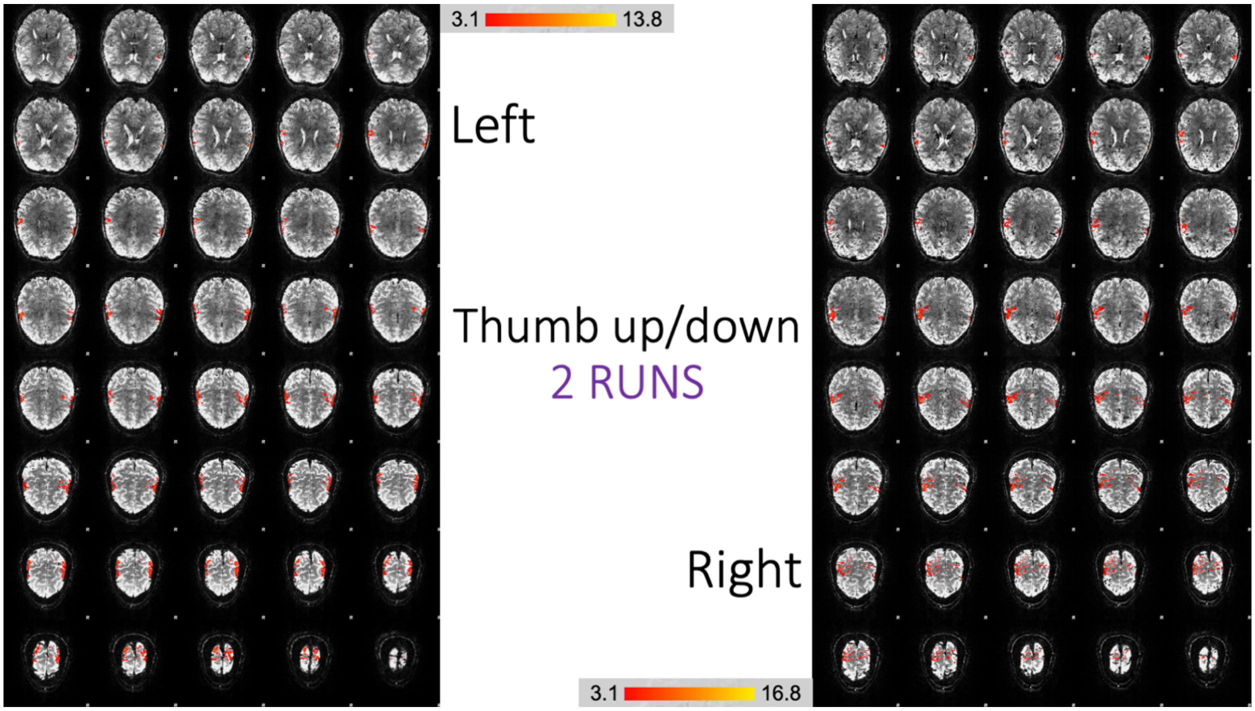


**Figure S4.1.** *Comparing 7 runs, 5 runs, 3 runs and 2 runs with the thumb up/down movement*

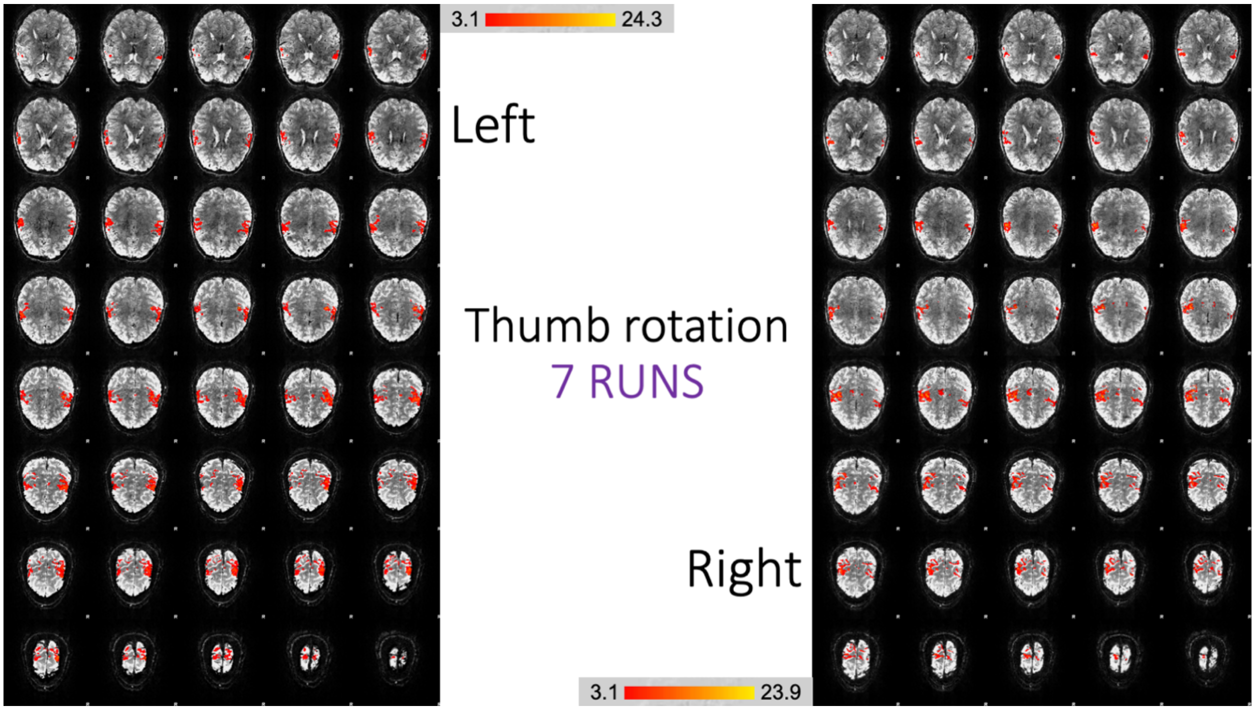
3.1
24.3
Left
Thumb rotation
7 RUNS
Right
3.1
23.9

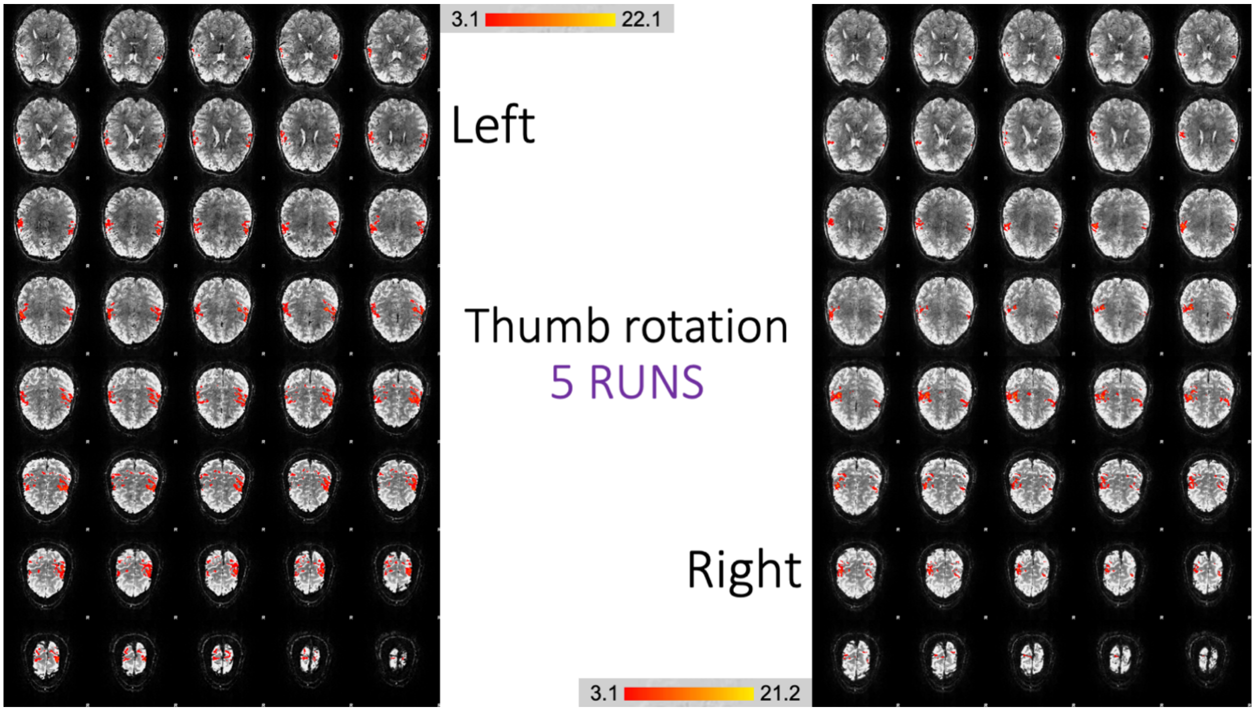
3.1
22.1
Left
Thumb rotation
5 RUNS
Right
3.1
21.2

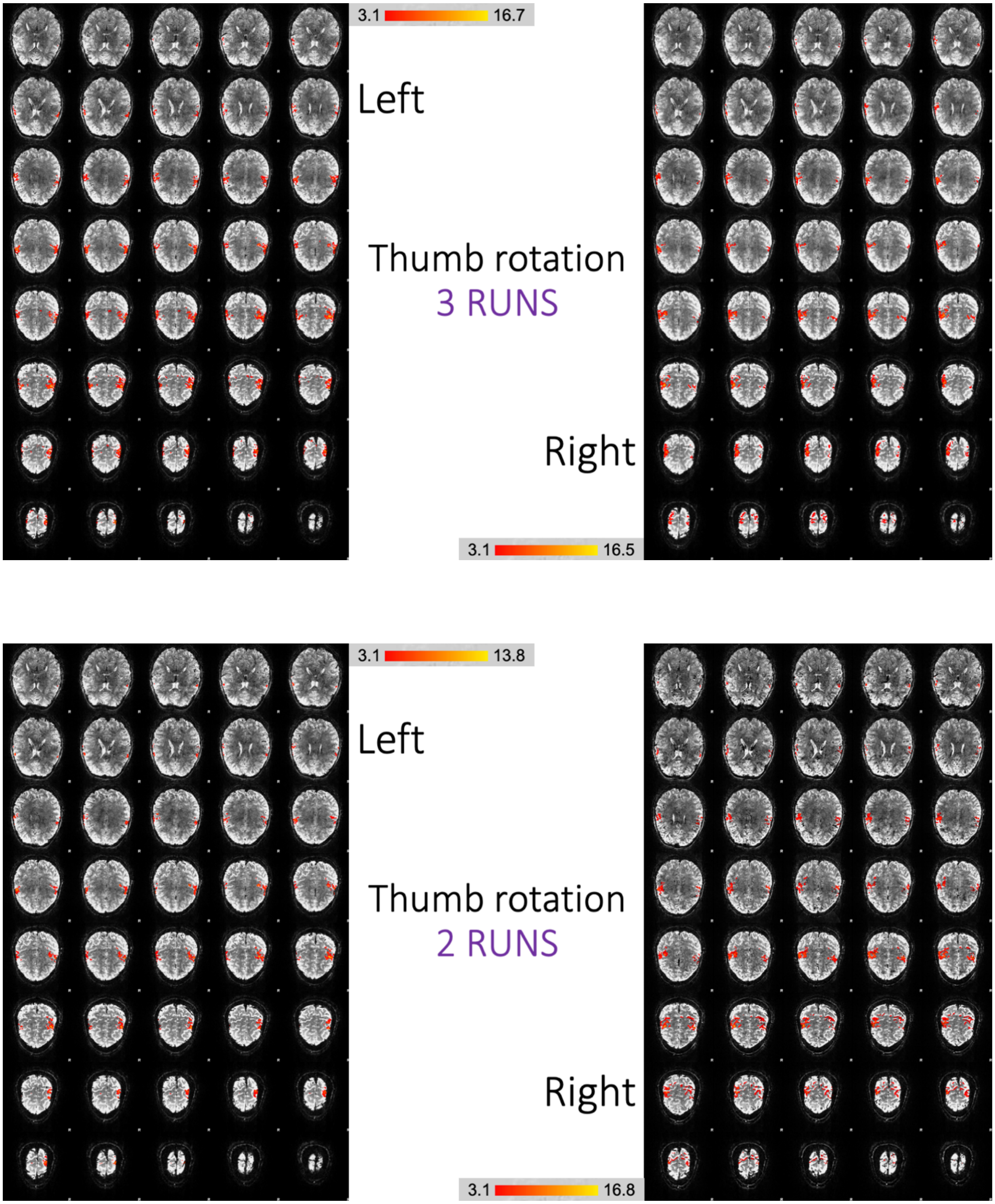


**Figure S4.2.** *Comparing 7 runs, 5 runs, 3 runs and 2 runs with thumb rotation*

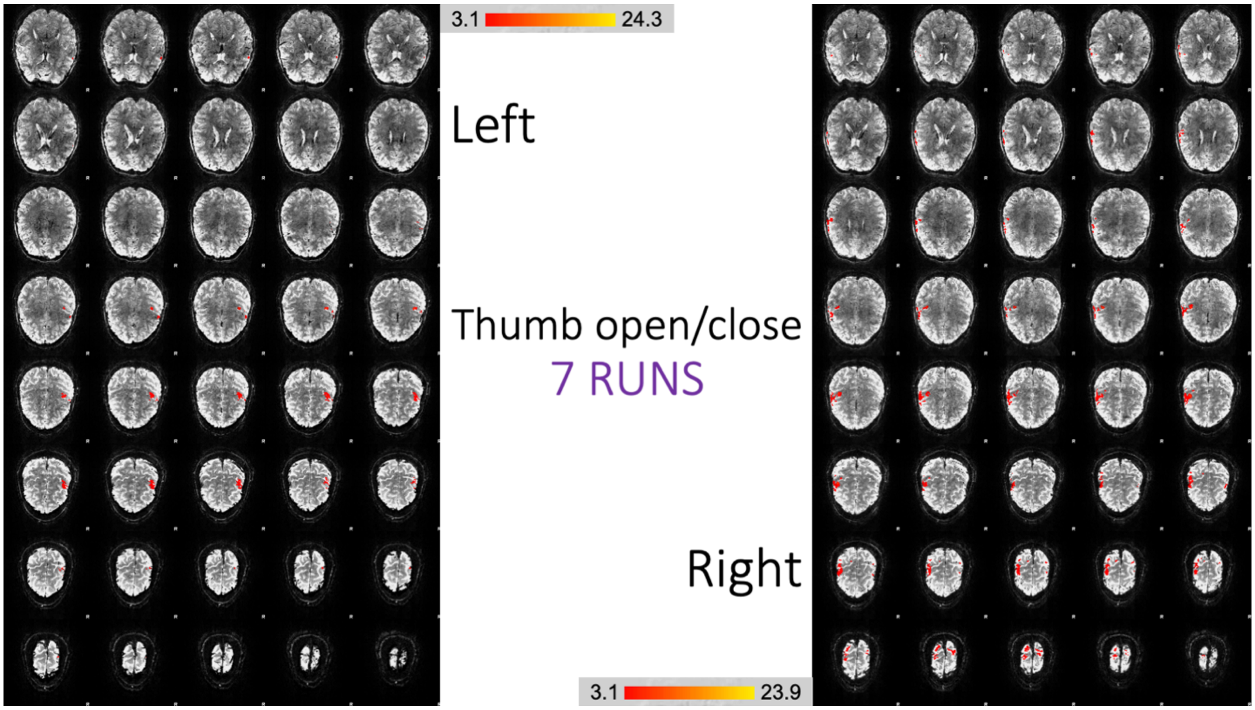
3.1
24.3
Left
Thumb open/close
7 RUNS
Right
3.1
23.9

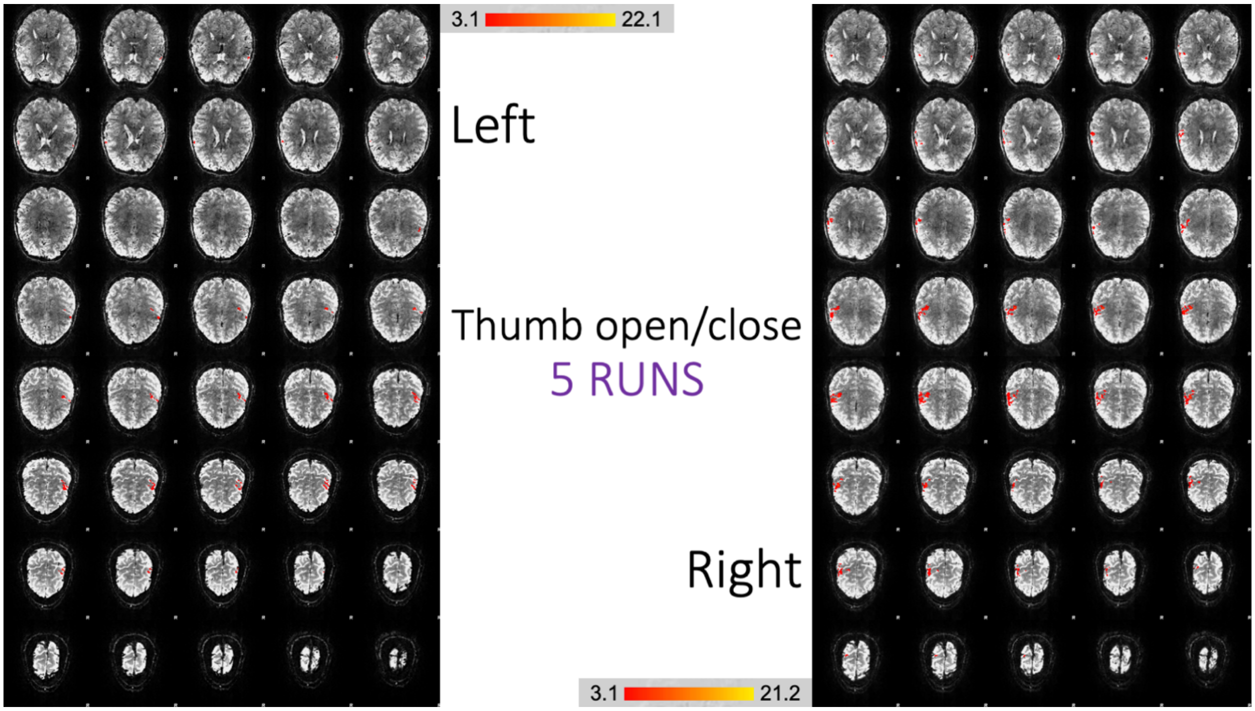
3.1
22.1
Left
Thumb open/close
5 RUNS
Right
3.1
21.2

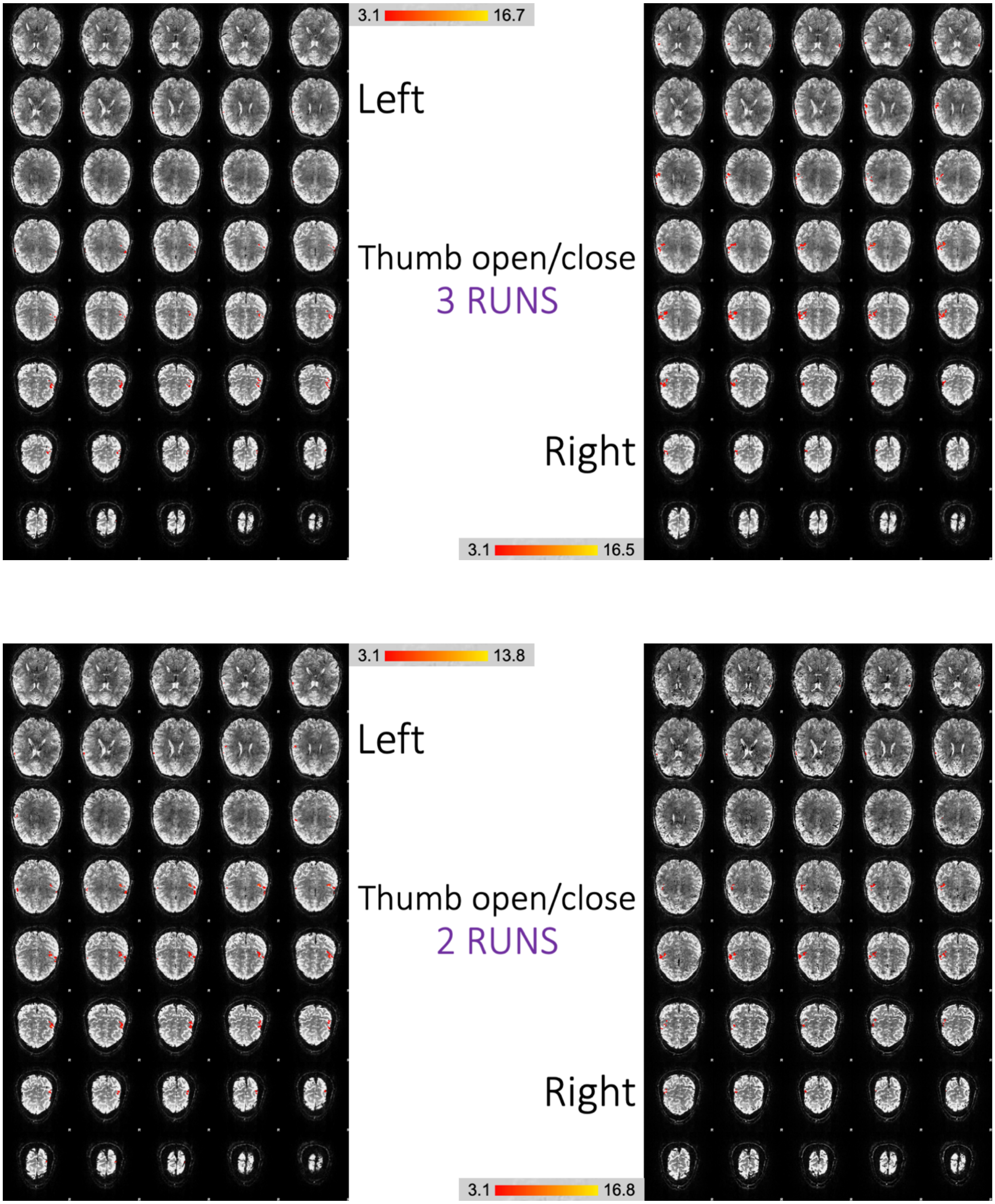


**Figure S4.3.** *Comparing 7 runs, 5 runs, 3 runs and 2 runs with the thumb open/close movement*

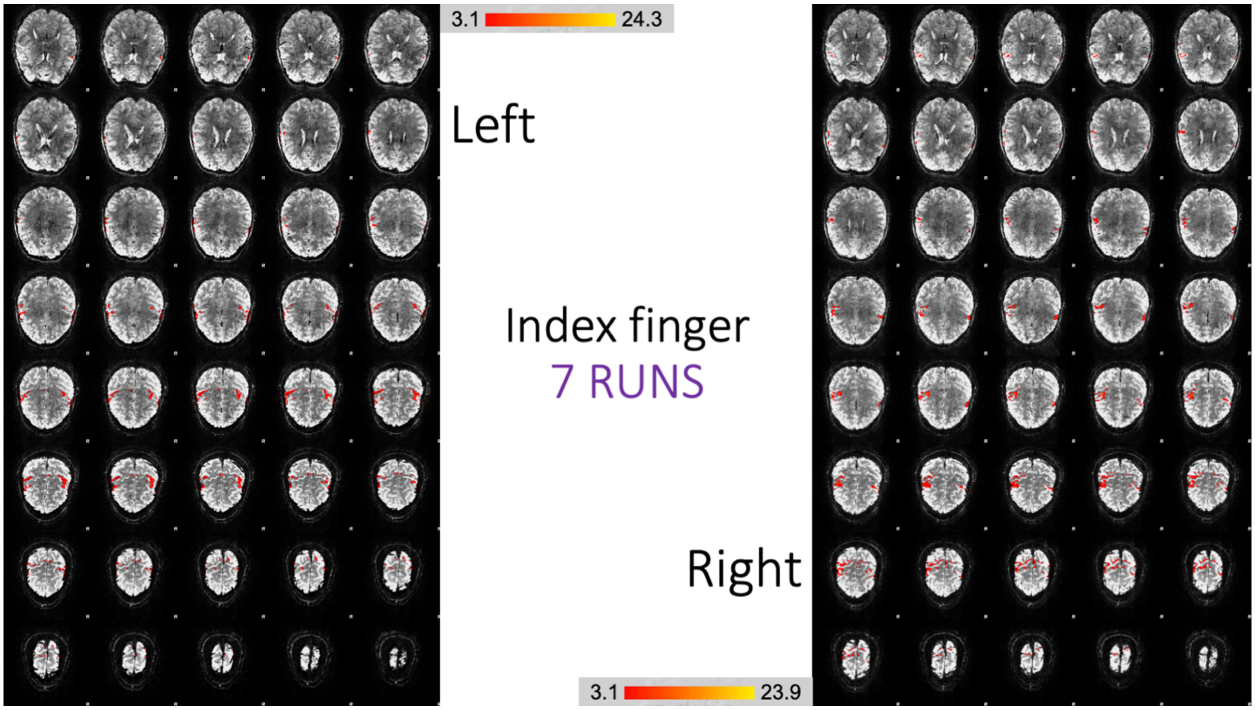
3.1
24.3
Left
Index finger
7 RUNS
Right
3.1
23.9

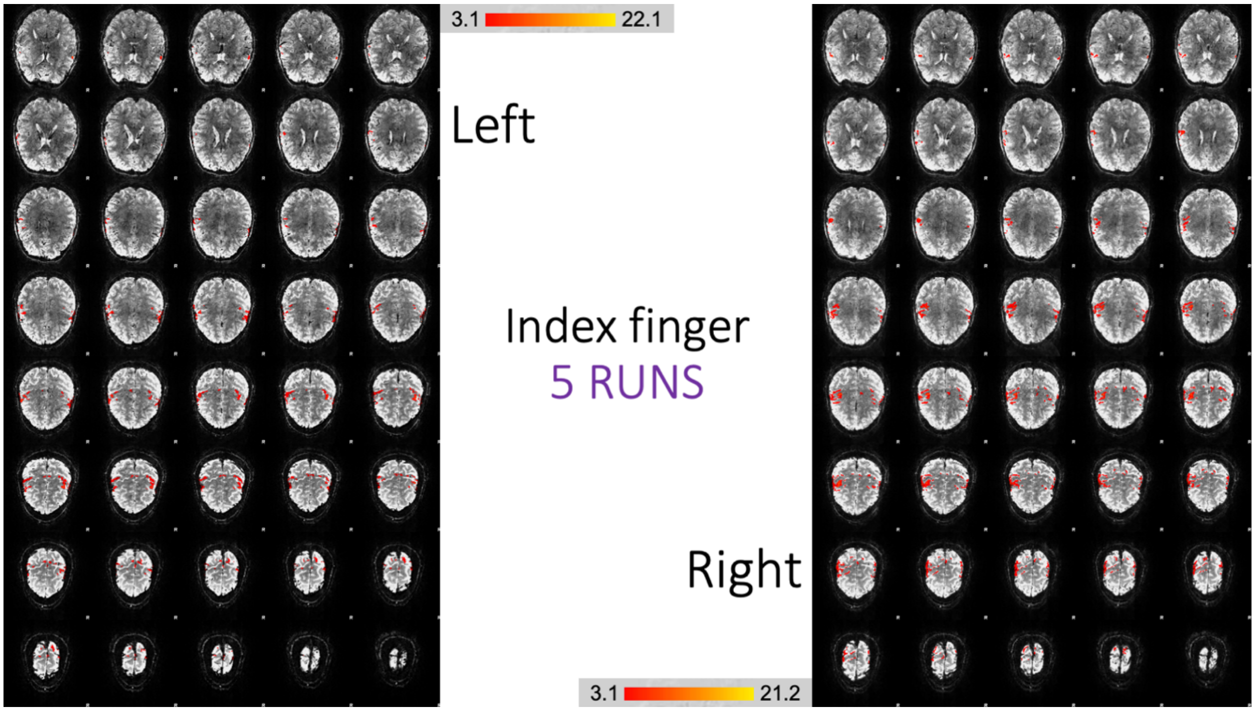
3.1
22.1
Left
Index finger
5 RUNS
Right
3.1
21.2

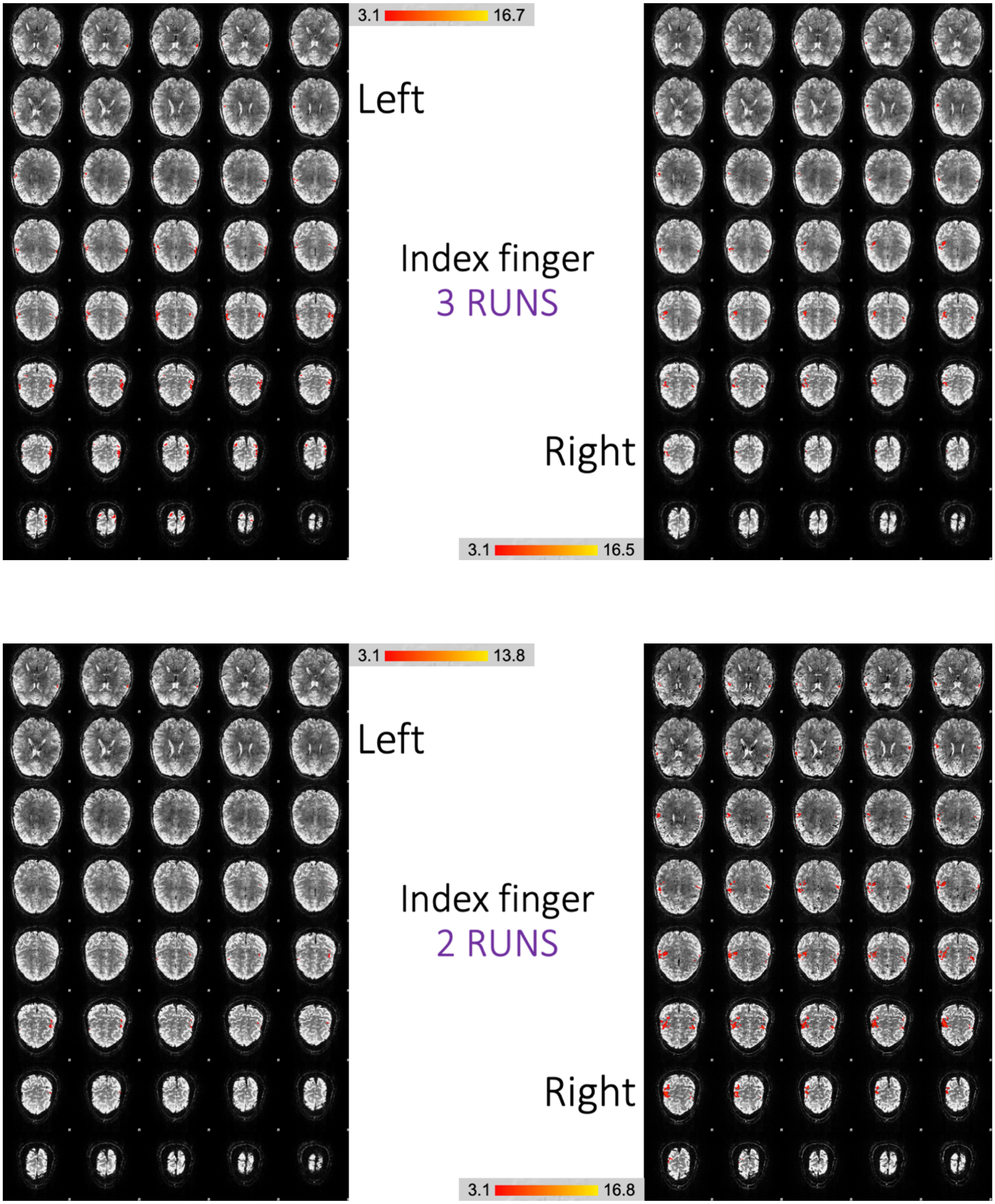


**Figure S4.4.** *Comparing 7 runs, 5 runs, 3 runs and 2 runs with the index open/close movement*

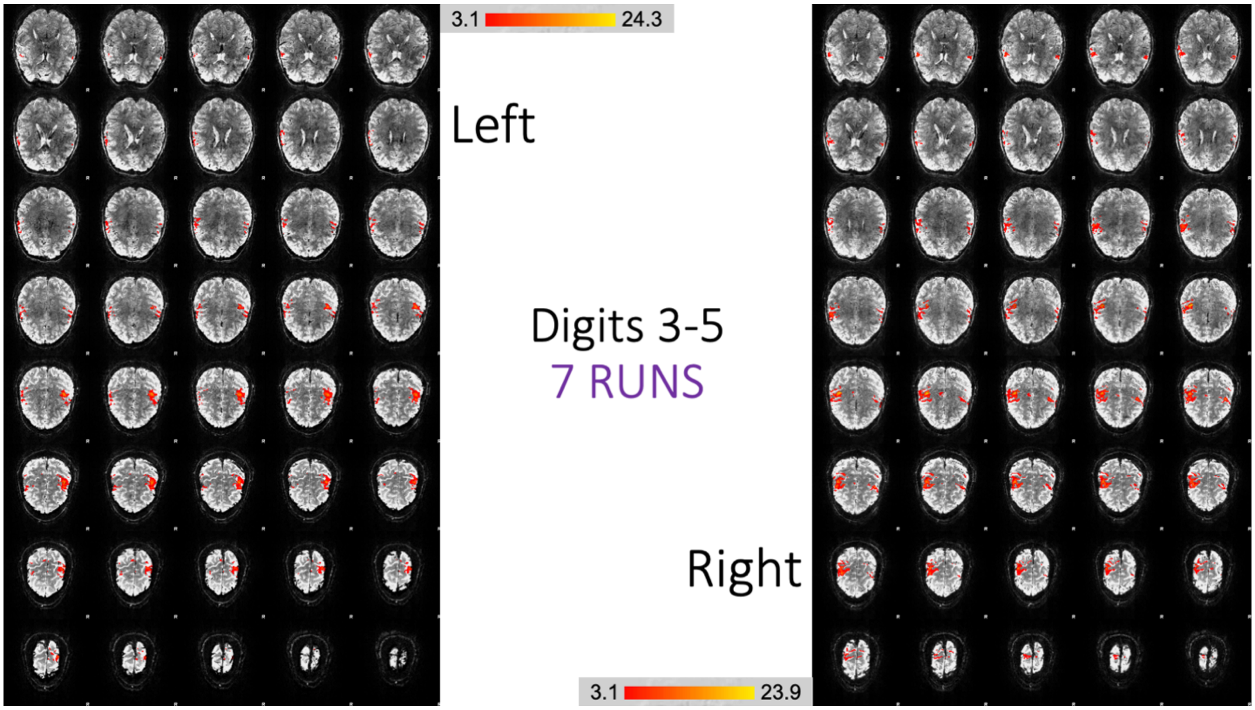
3.1
24.3
Left
Digits 3-5
7 RUNS
Right
3.1
23.9

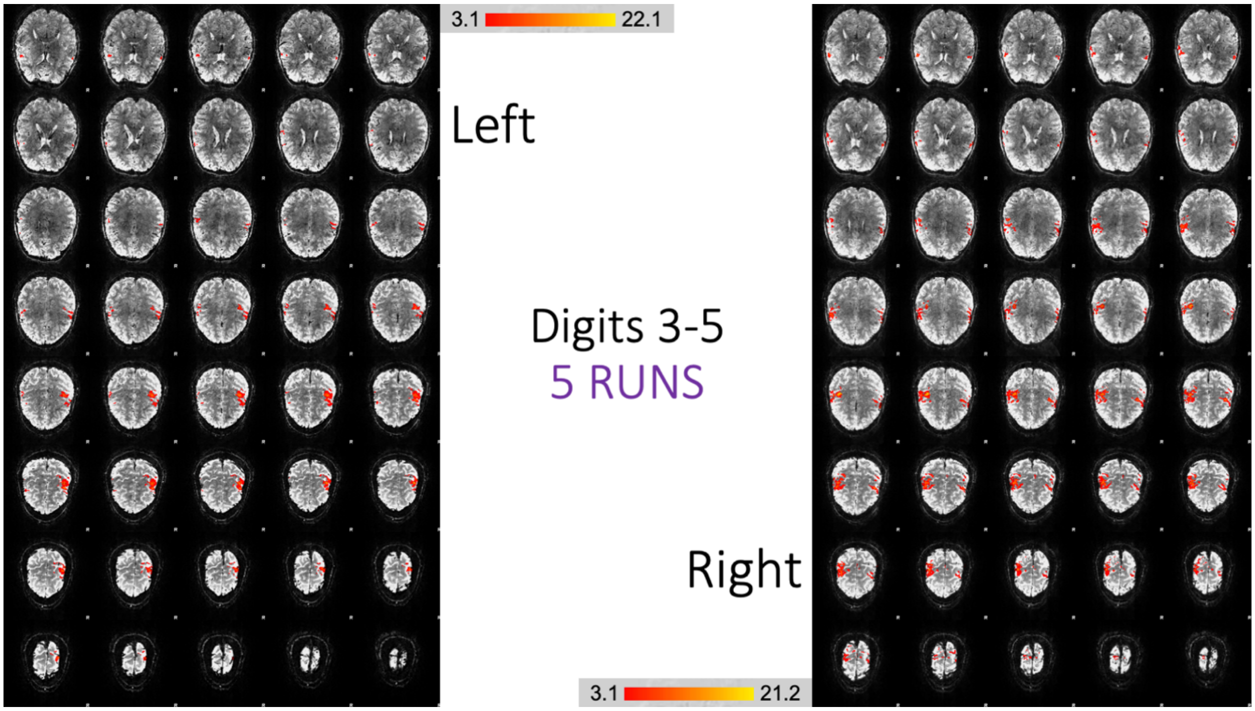
3.1
22.1
Left
Digits 3-5
5 RUNS
Right
3.1
21.2

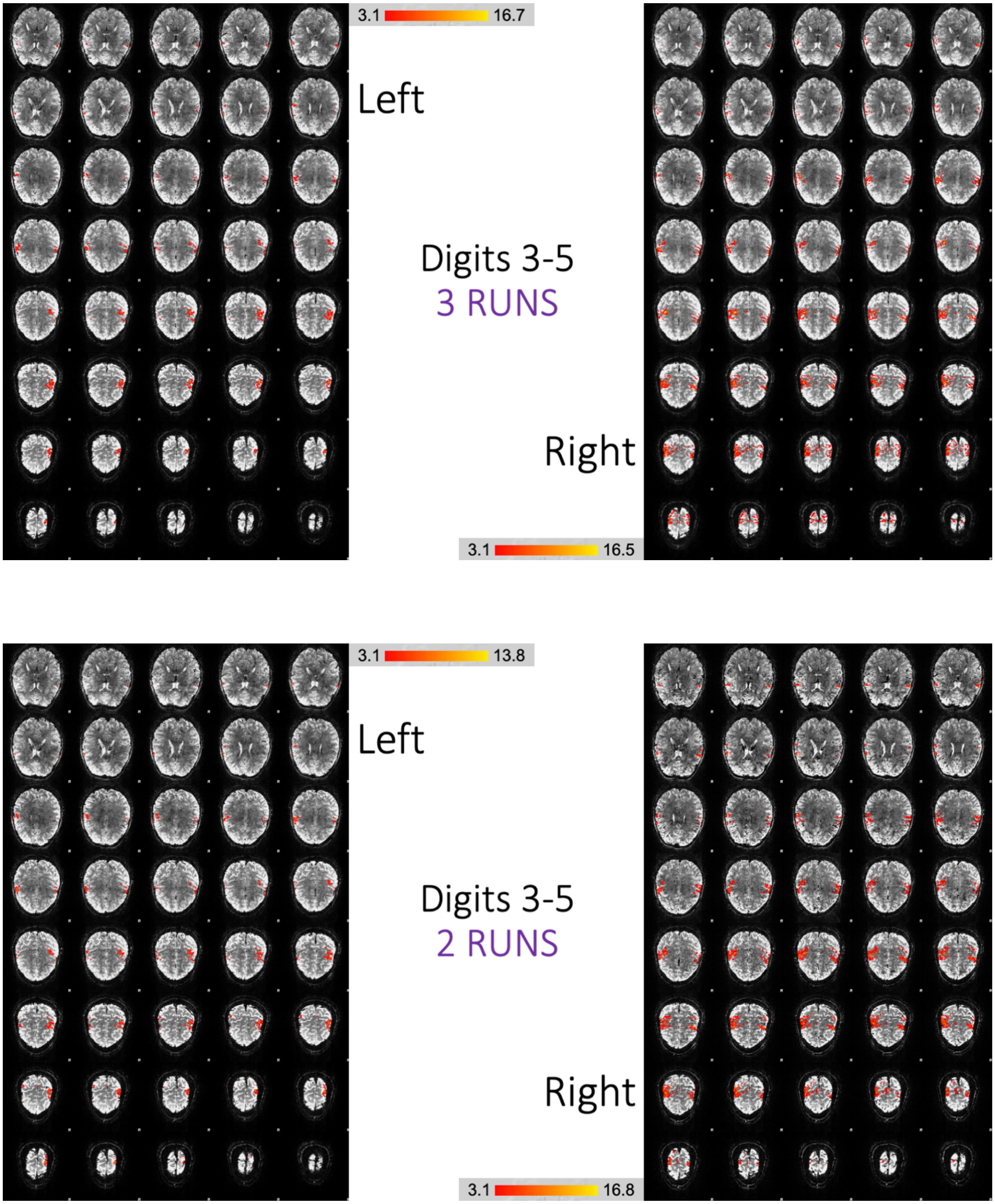


**Figure S4.5.** *Comparing 7 runs, 5 runs, 3 runs and 2 runs with digits 3-5 open/close movement*

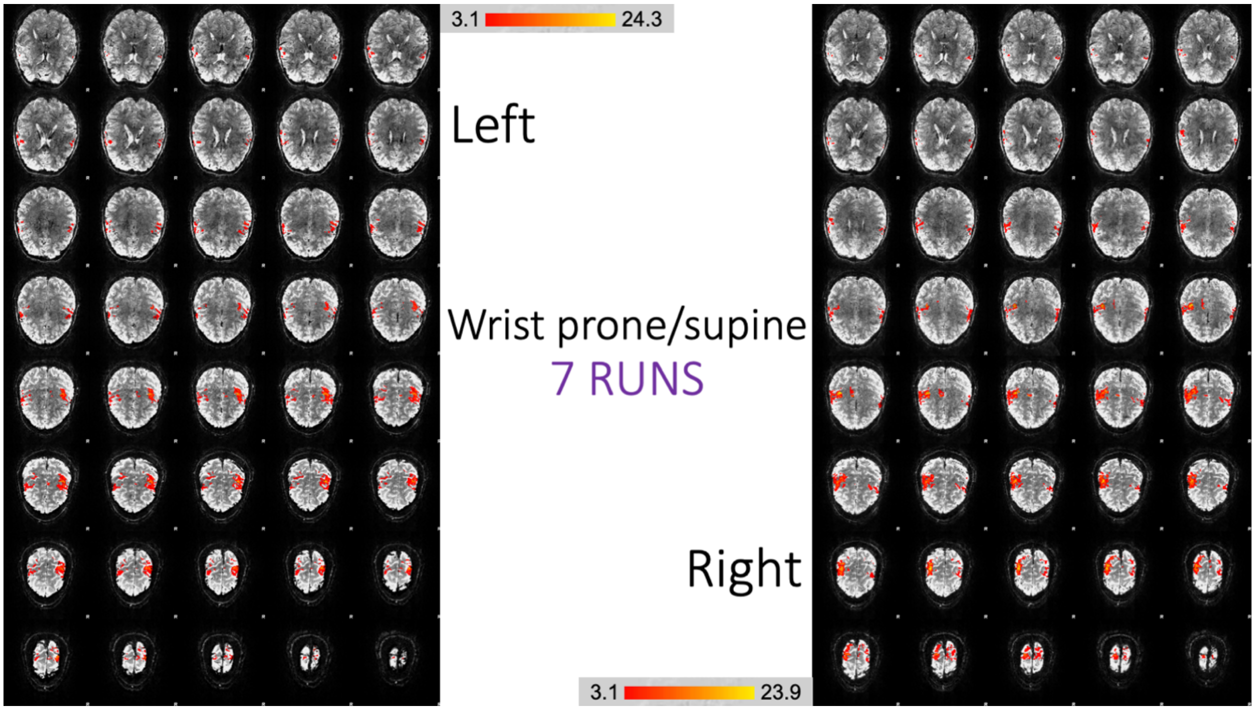
3.1
24.3
Left
Wrist prone/supine
7 RUNS
Right
3.1
23.9

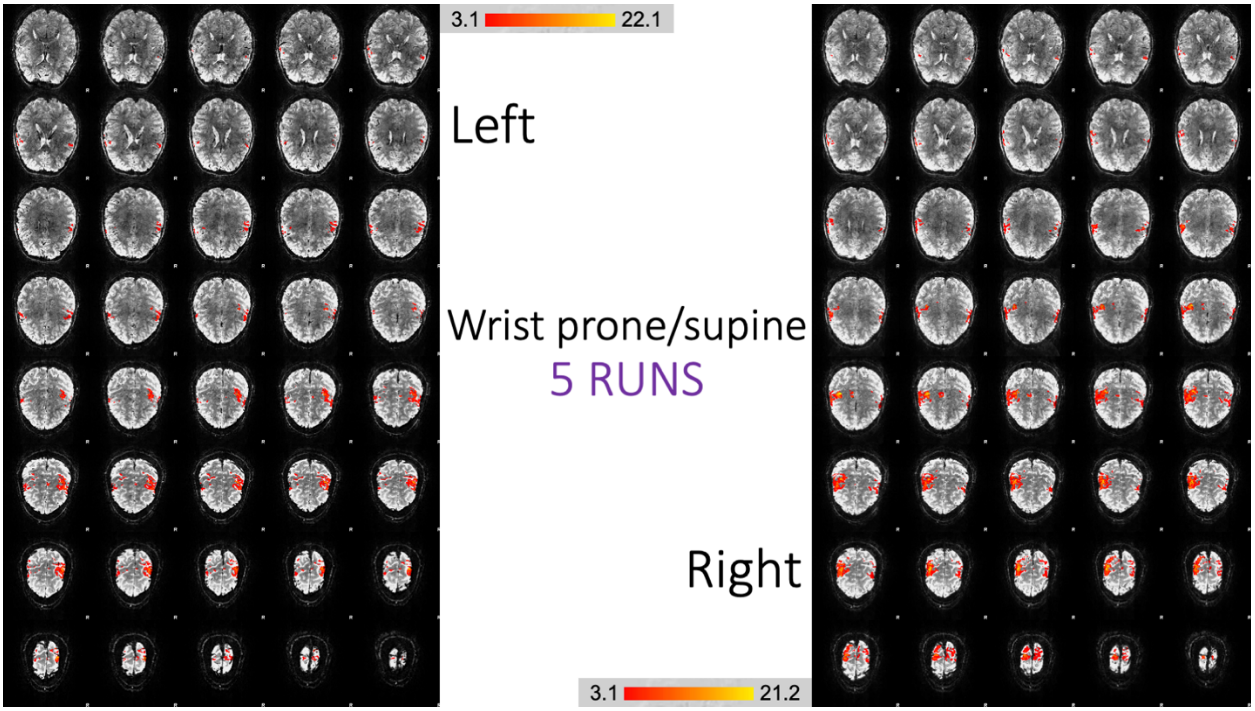
3.1
22.1
Left
Wrist prone/supine
5 RUNS
Right
3.1
21.2

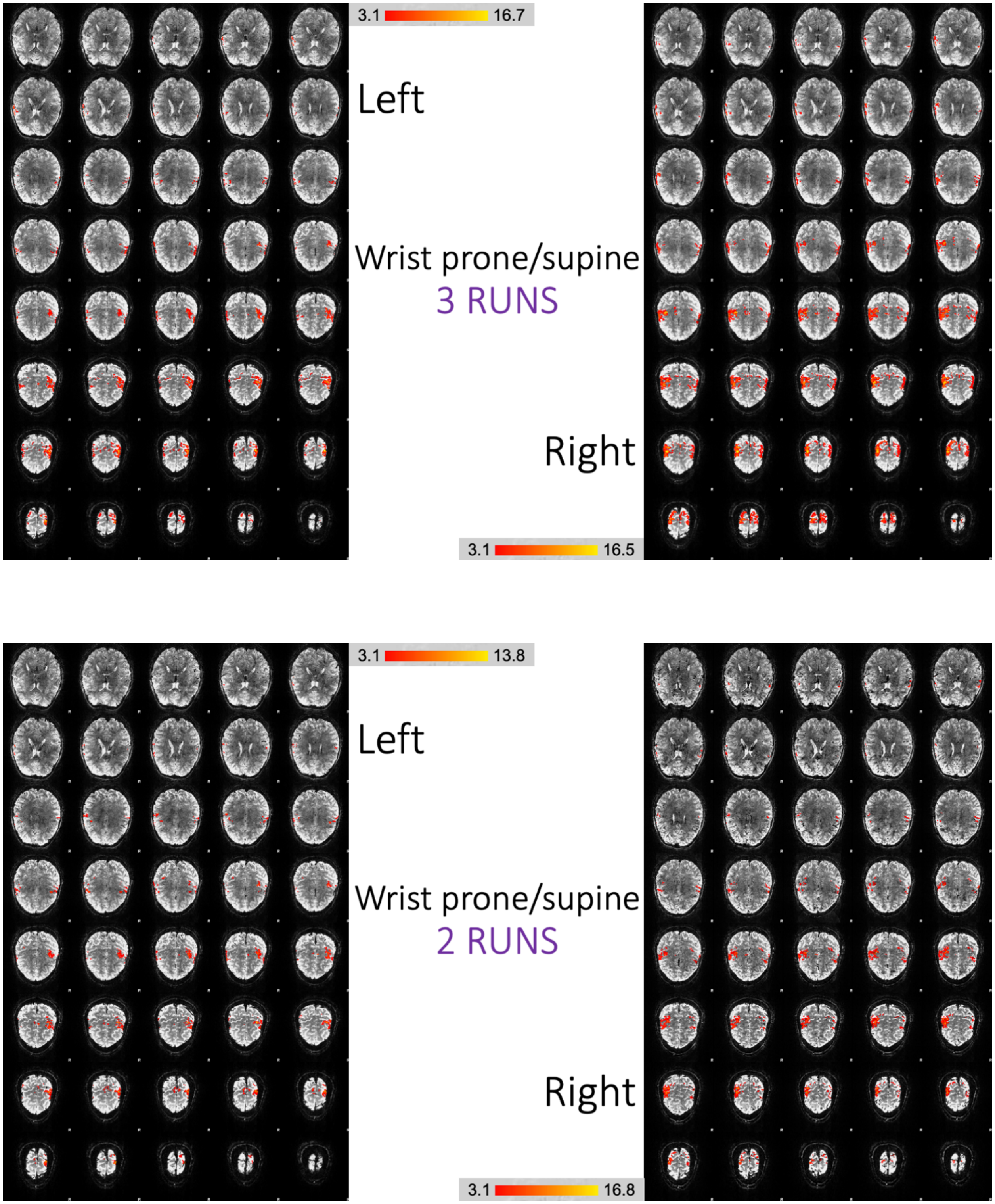


**Figure S4.6.** *Comparing 7 runs, 5 runs, 3 runs and 2 runs with the wrist prone/supine movement*

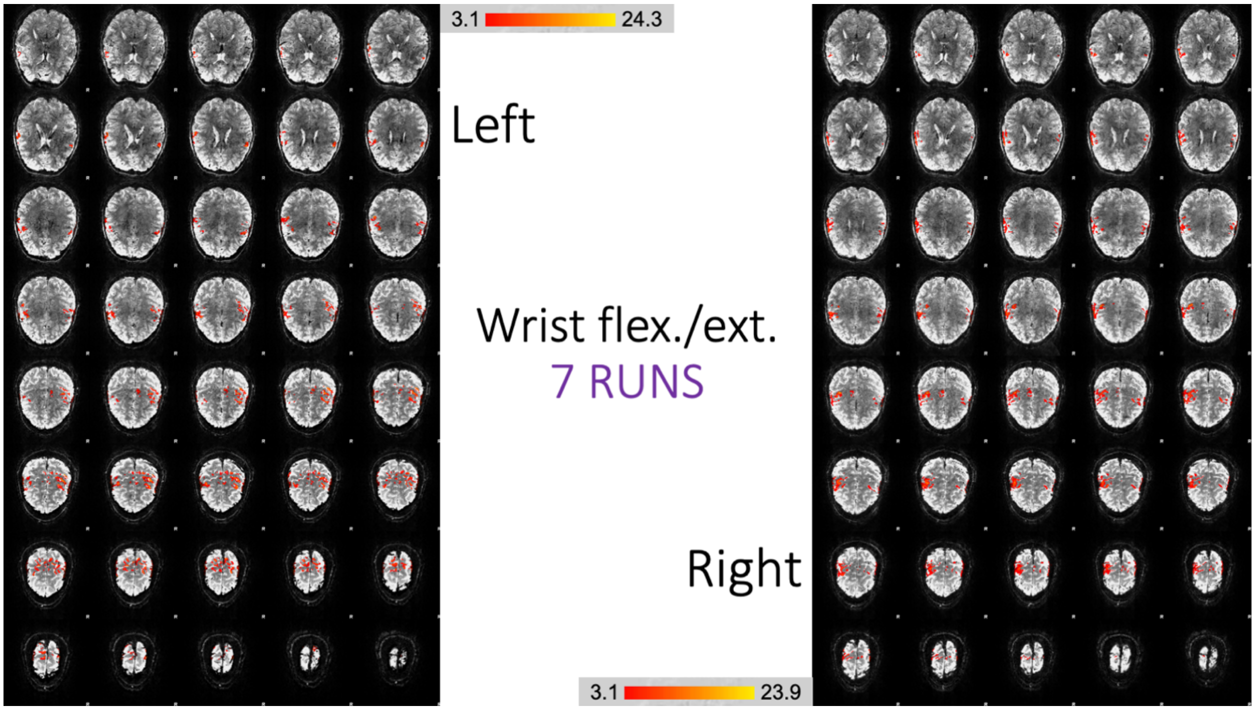
3.1
24.3
Left
Wrist flex./ext.
7 RUNS
Right
3.1
23.9

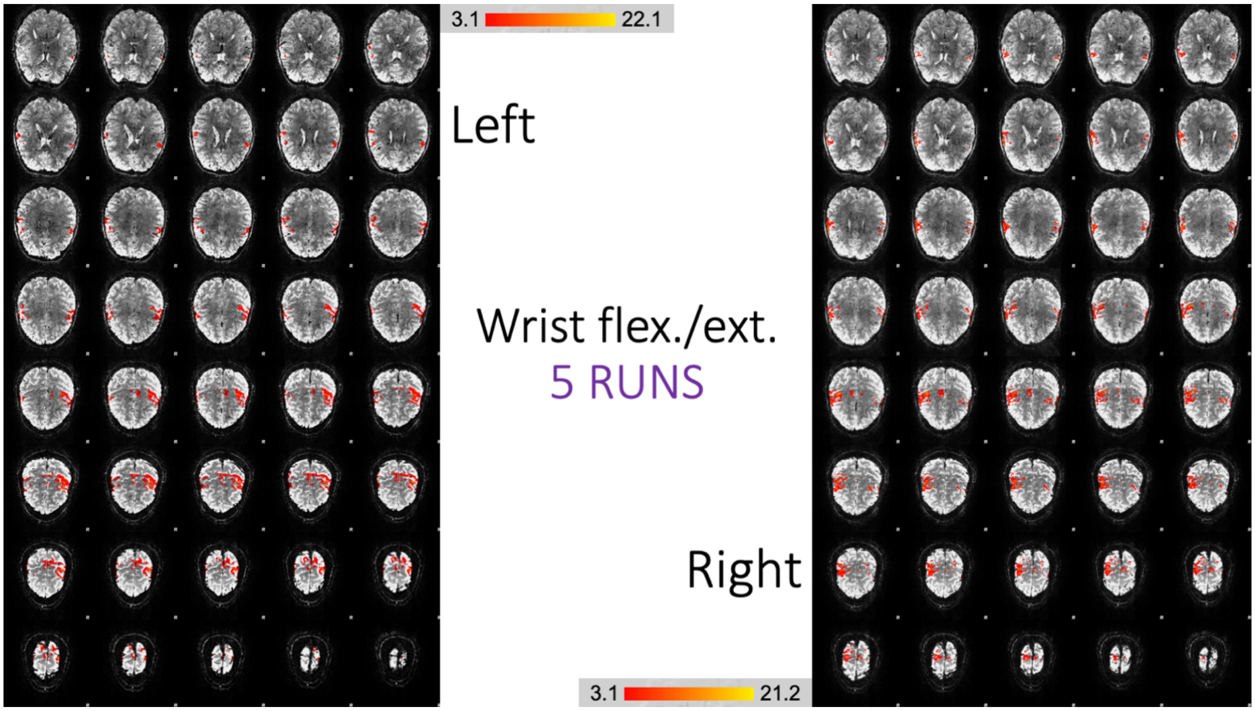
3.1
22.1
Left
Wrist flex./ext.
5 RUNS
Right
3.1
21.2

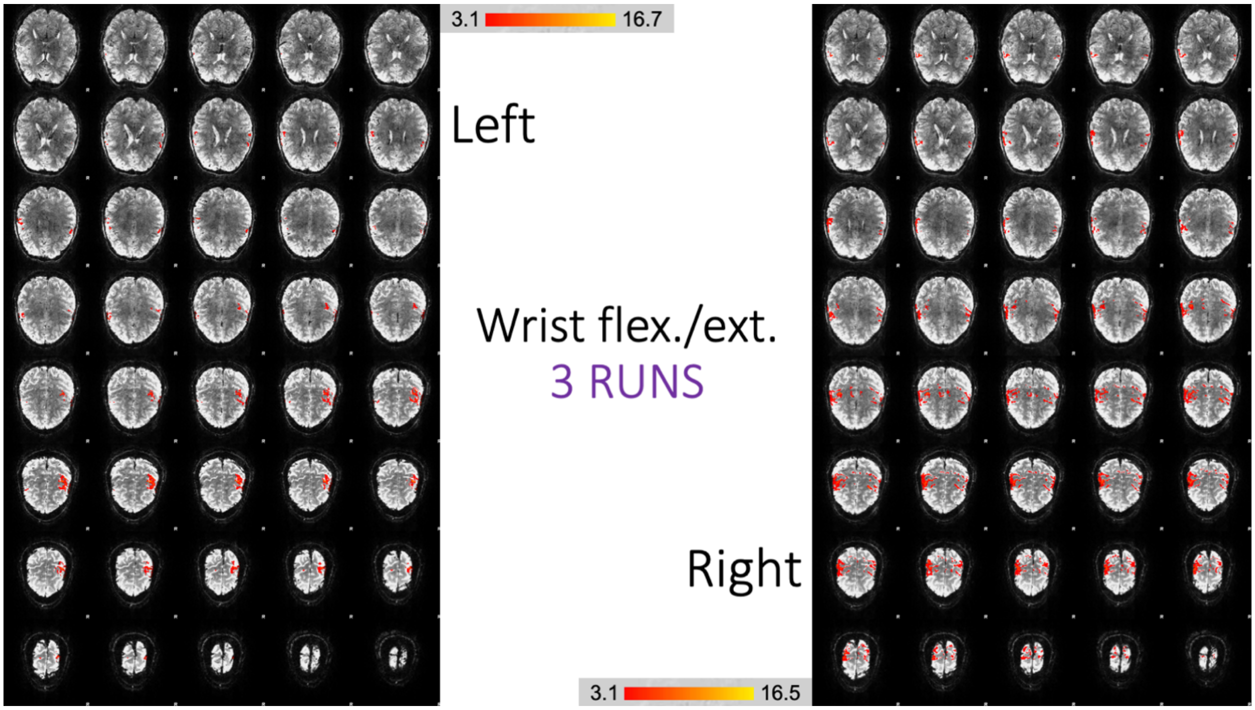


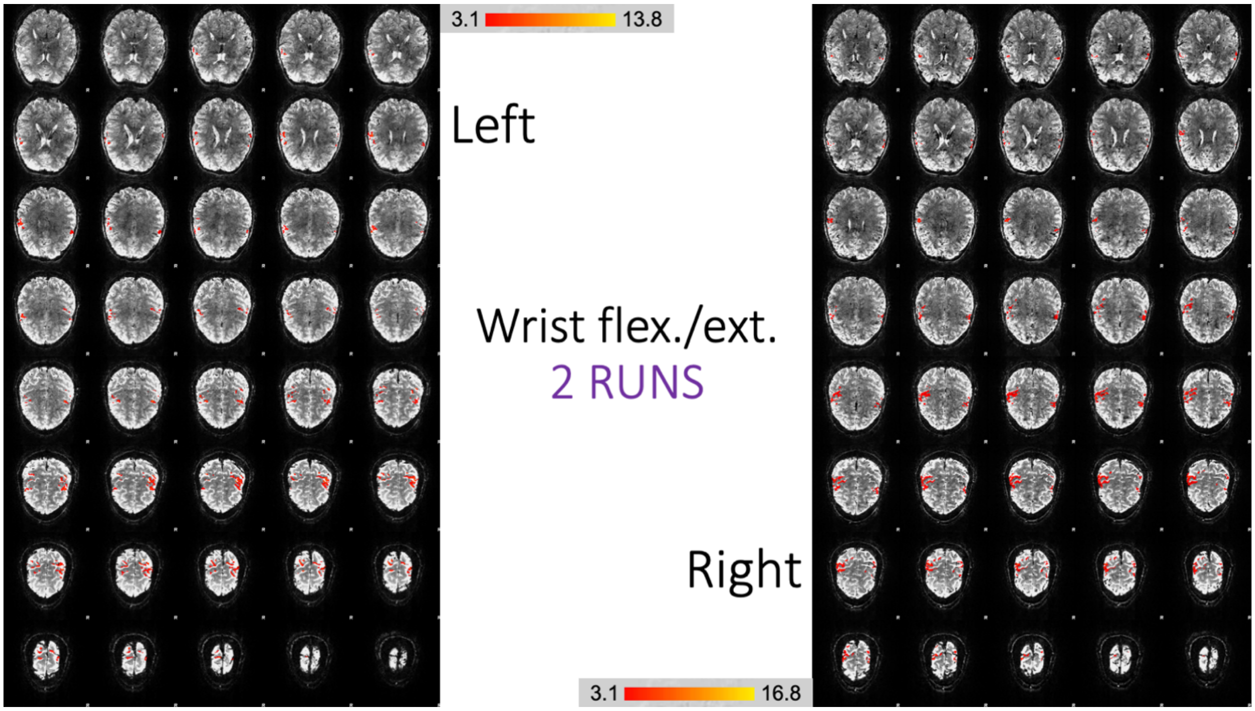


**Figure S4.7.** *Comparing 7 runs, 5 runs, 3 runs and 2 runs with the wrist flexion/ext. movement*

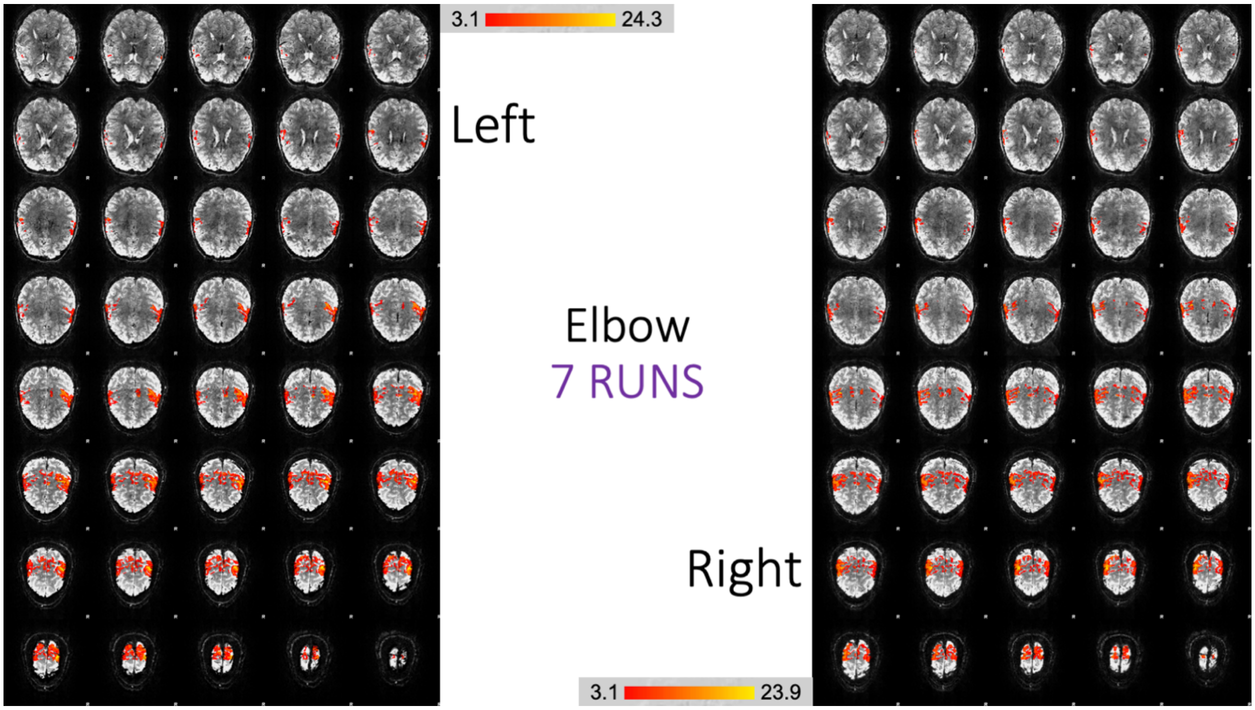
3.1
24.3
Left
Elbow
7 RUNS
Right
3.1
23.9

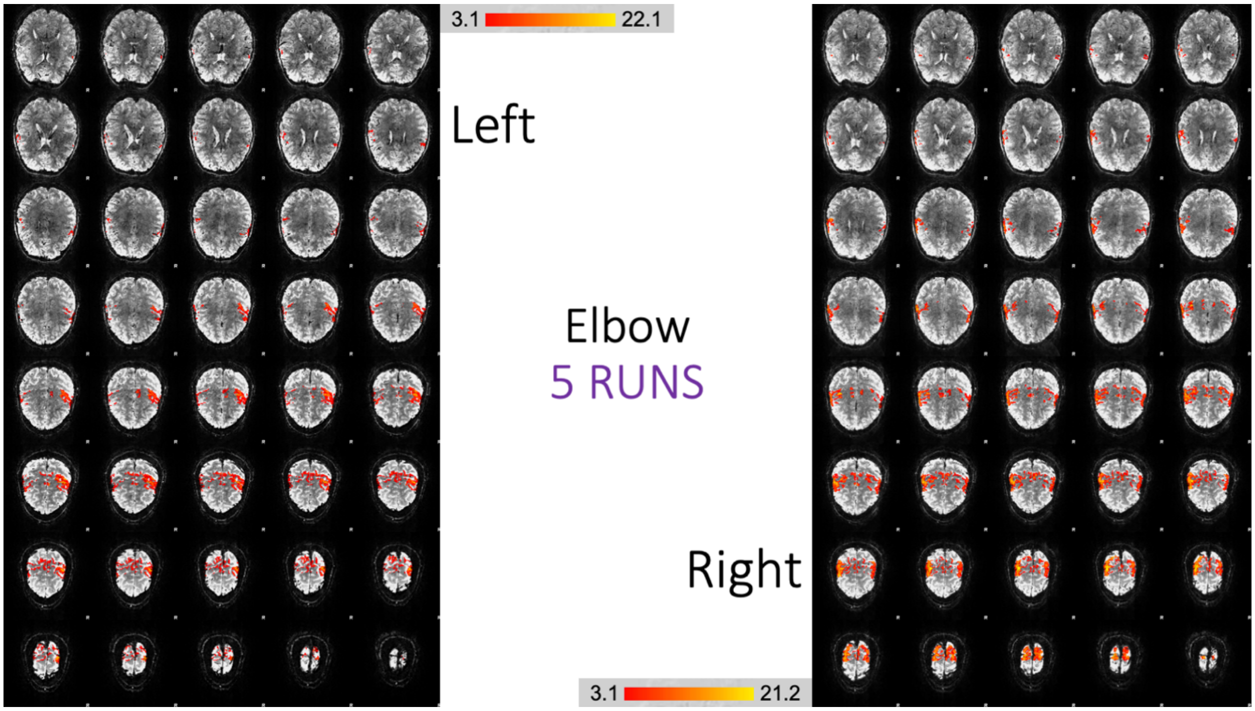
3.1
22.1
Left
Elbow
5 RUNS
Right
3.1
21.2

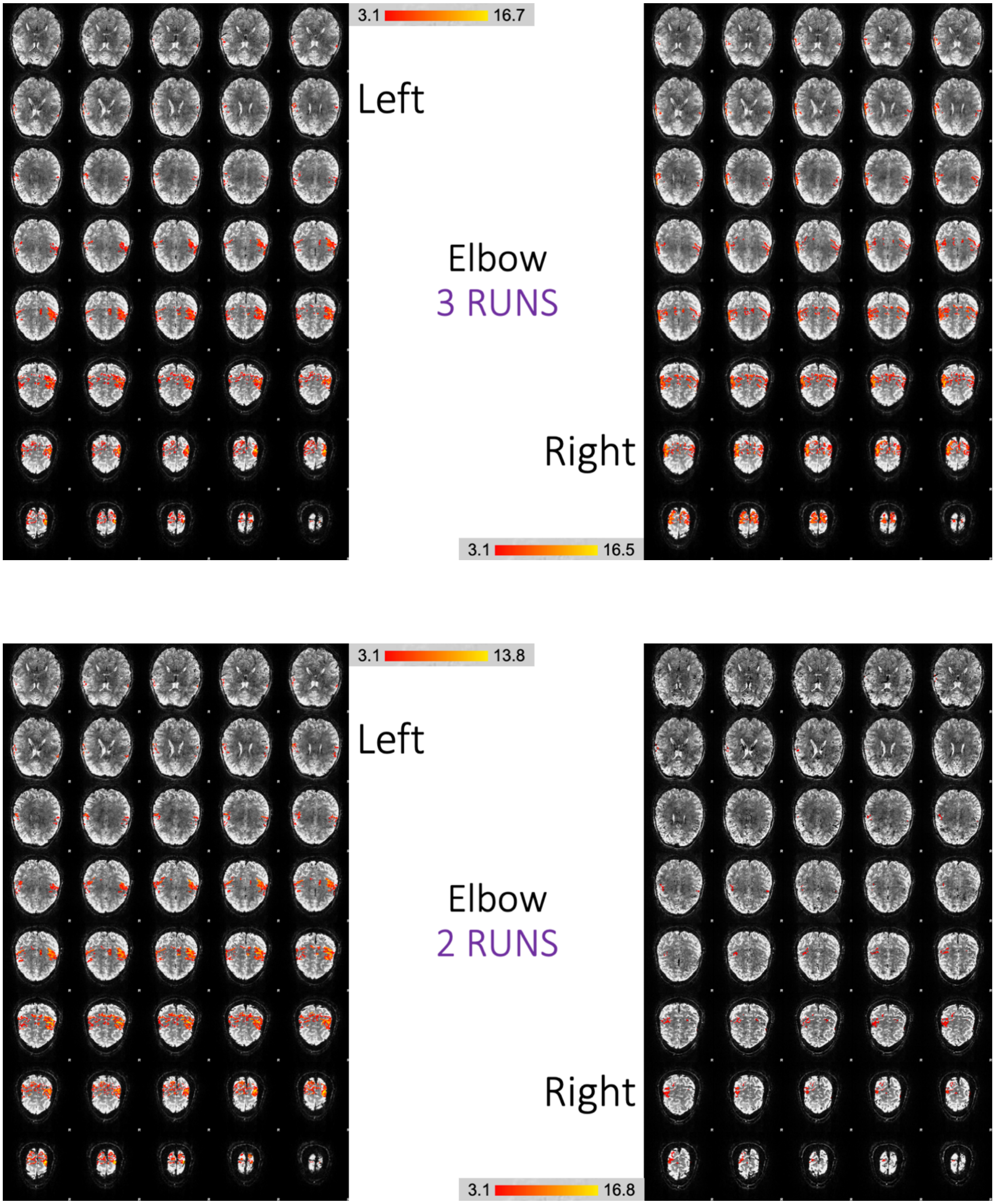


**Figure S4.8.** *Comparing 7 runs, 5 runs, 3 runs and 2 runs with the elbow open/close movement*

### S5. 3D overlay images for different number of runs

3D overlay images were generated using Mango for each of the 8 movements (left and right arms), comparing 7, 5 and 3 runs cases.

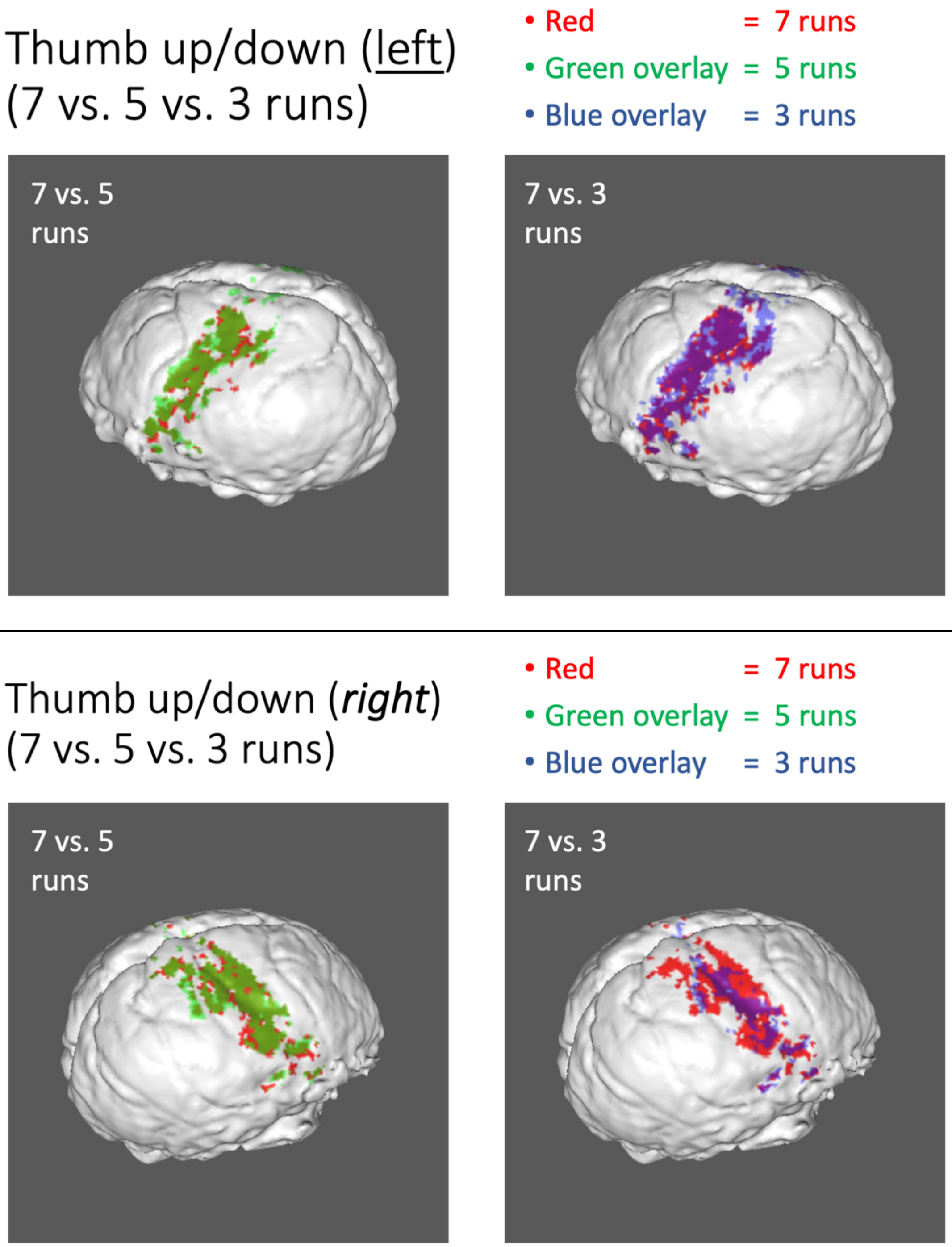


**Figure S5.1.** *3D overlays comparing runs (thumb up/down). Robust activations with high similarity were noted (left: 7 vs. 5* $D_c^{7\text{-}5}$*=0.64,* $D_c^{7\text{-}3}$*=0.55; right:* $D_c^{7\text{-}5}$*=0.74,* $D_c^{7\text{-}3}$*=0.43).*

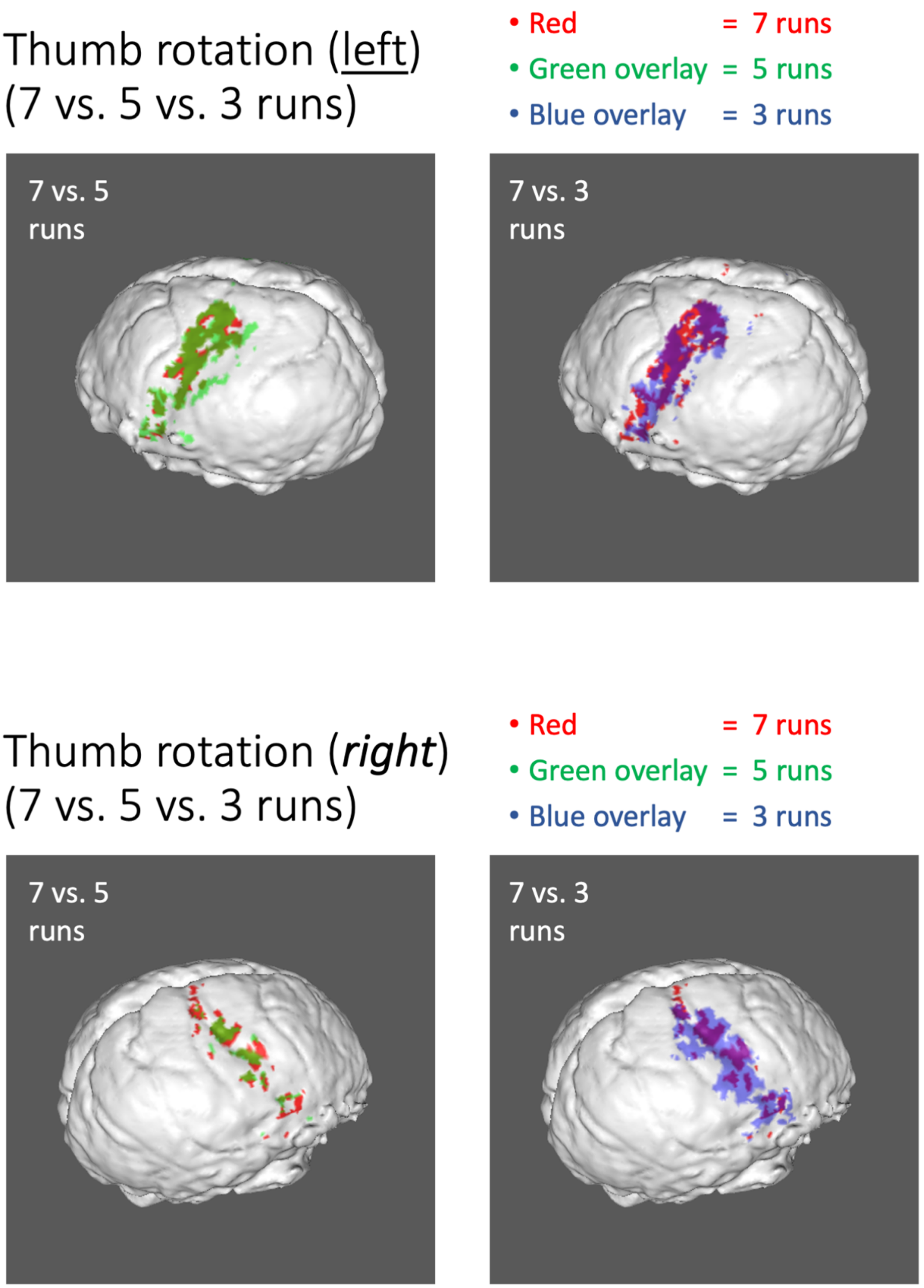


**Figure S5.2.** *3D overlays comparing runs (thumb rotation). Robust activations with high similarity were noted (left: $D_c^{7\text{-}5}=0.71$, $D_c^{7\text{-}3}=0.51$; right: $D_c^{7\text{-}5}=0.63$, $D_c^{7\text{-}3}=0.47$).*

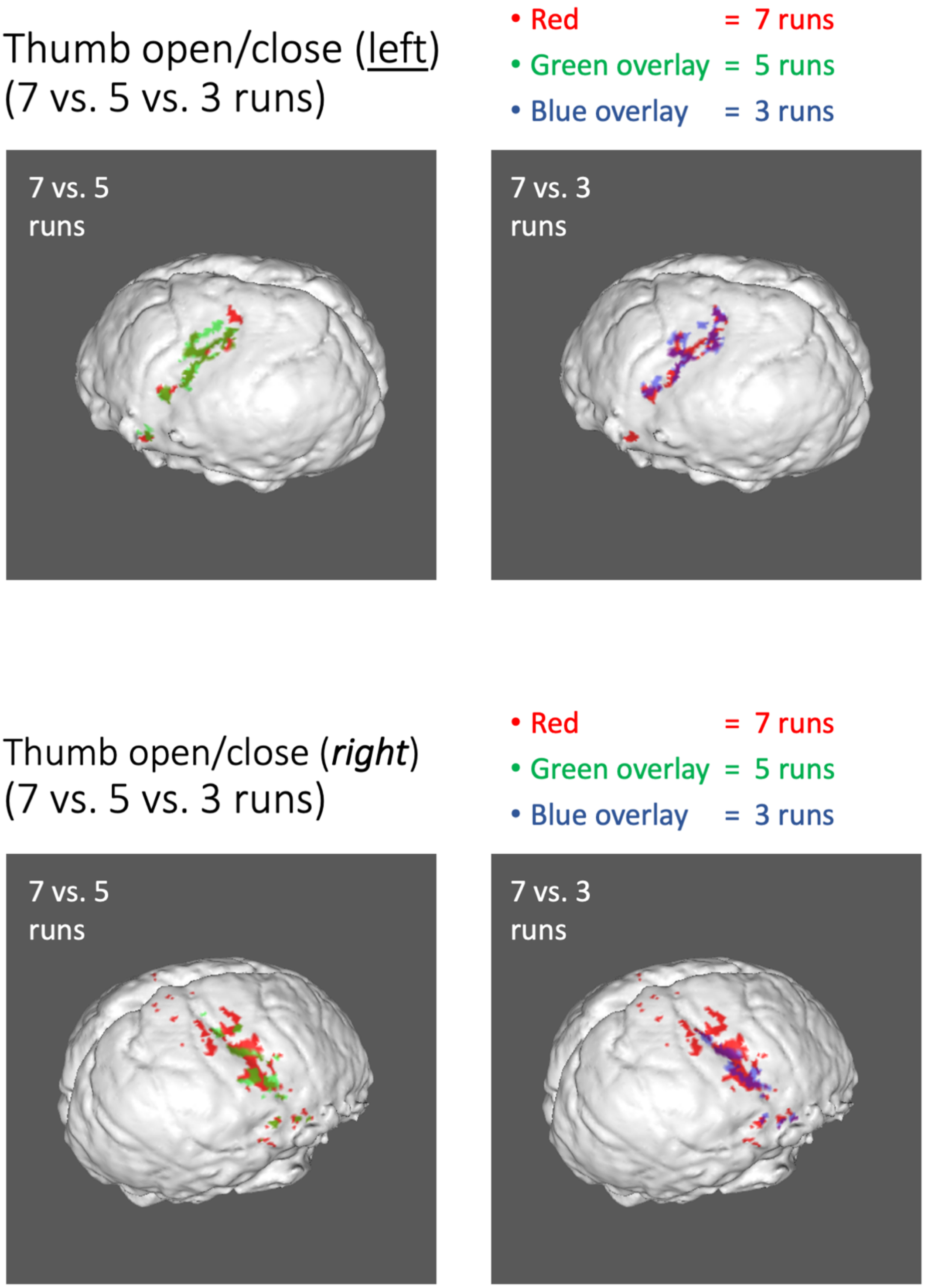


**Figure S5.3.** *3D overlays comparing runs (thumb open/close). Weaker activations with high similarity were noted (left:* $D_c^{7\text{-}5}$*=0.61,* $D_c^{7\text{-}3}$*=0.42; right:* $D_c^{7\text{-}5}$*=0.58,* $D_c^{7\text{-}3}$*=0.47).*

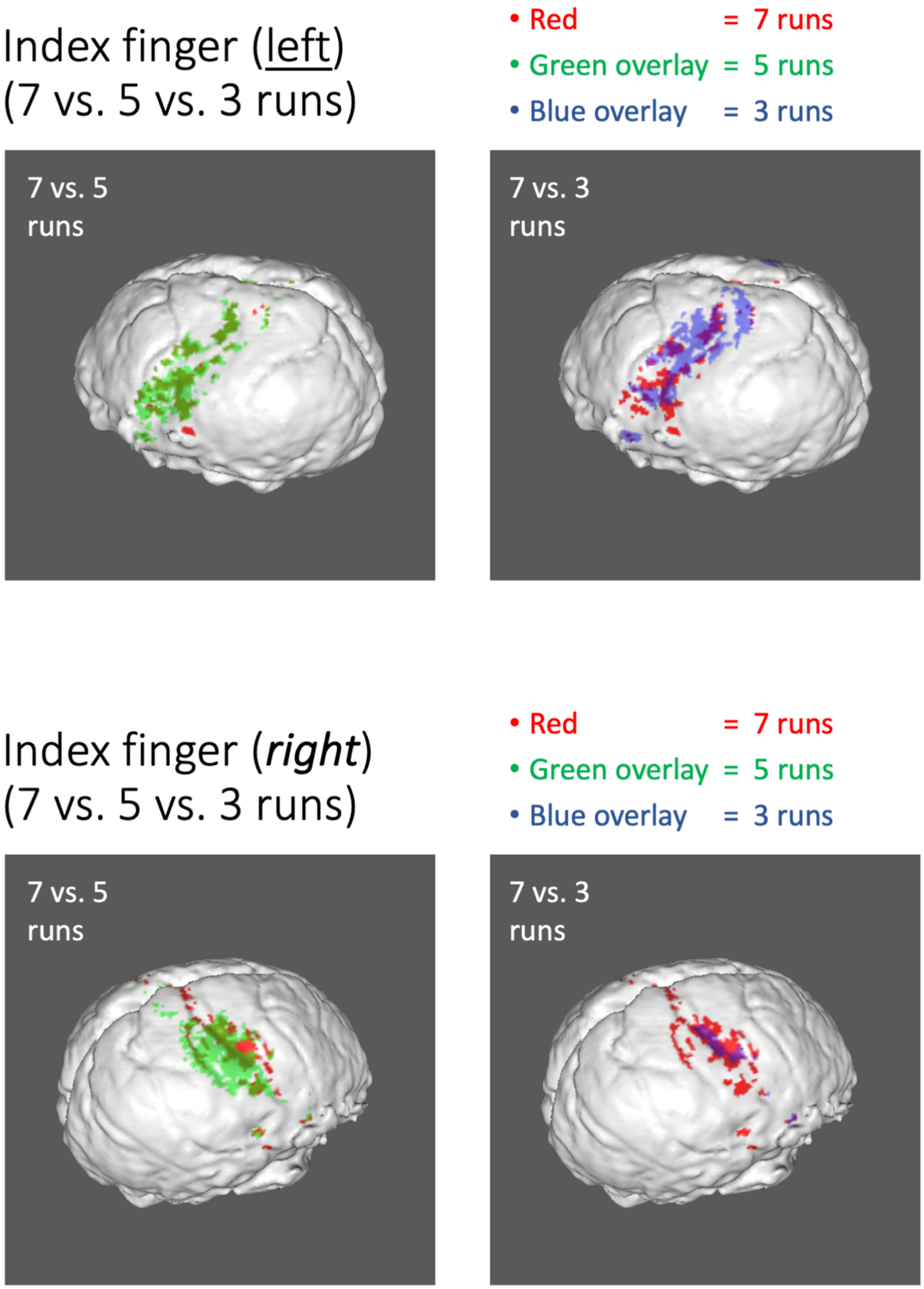


**Figure S5.4.** *3D overlays comparing runs (index finger). Weaker activations with high similarity were noted between 7 and 5 runs, but 3 runs exhibited poor similarity with 7 runs (left:* $D_c^{7\text{-}5}=0.78$*,* $D_c^{7\text{-}3}=0.23$*; right:* $D_c^{7\text{-}5}=0.41$*,* $D_c^{7\text{-}3}=0.27$*).*

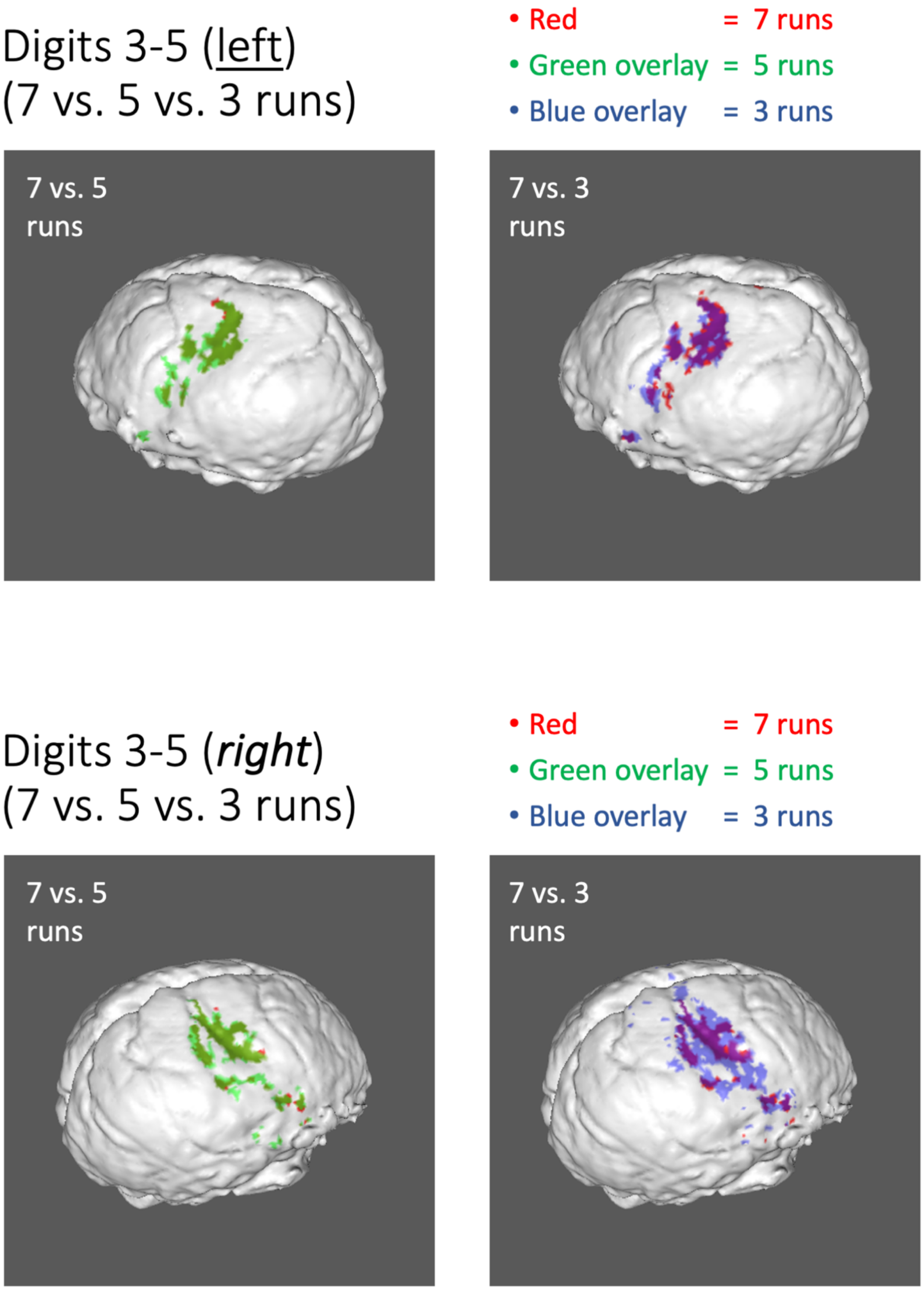


**Figure S5.5.** *3D overlays comparing runs (digits 3-5). Robust activations with high similarity were noted (left: $D_c^{7\text{-}5}$=0.69, $D_c^{7\text{-}3}$=0.50; right: $D_c^{7\text{-}5}$=0.76, $D_c^{7\text{-}3}$=0.59).*

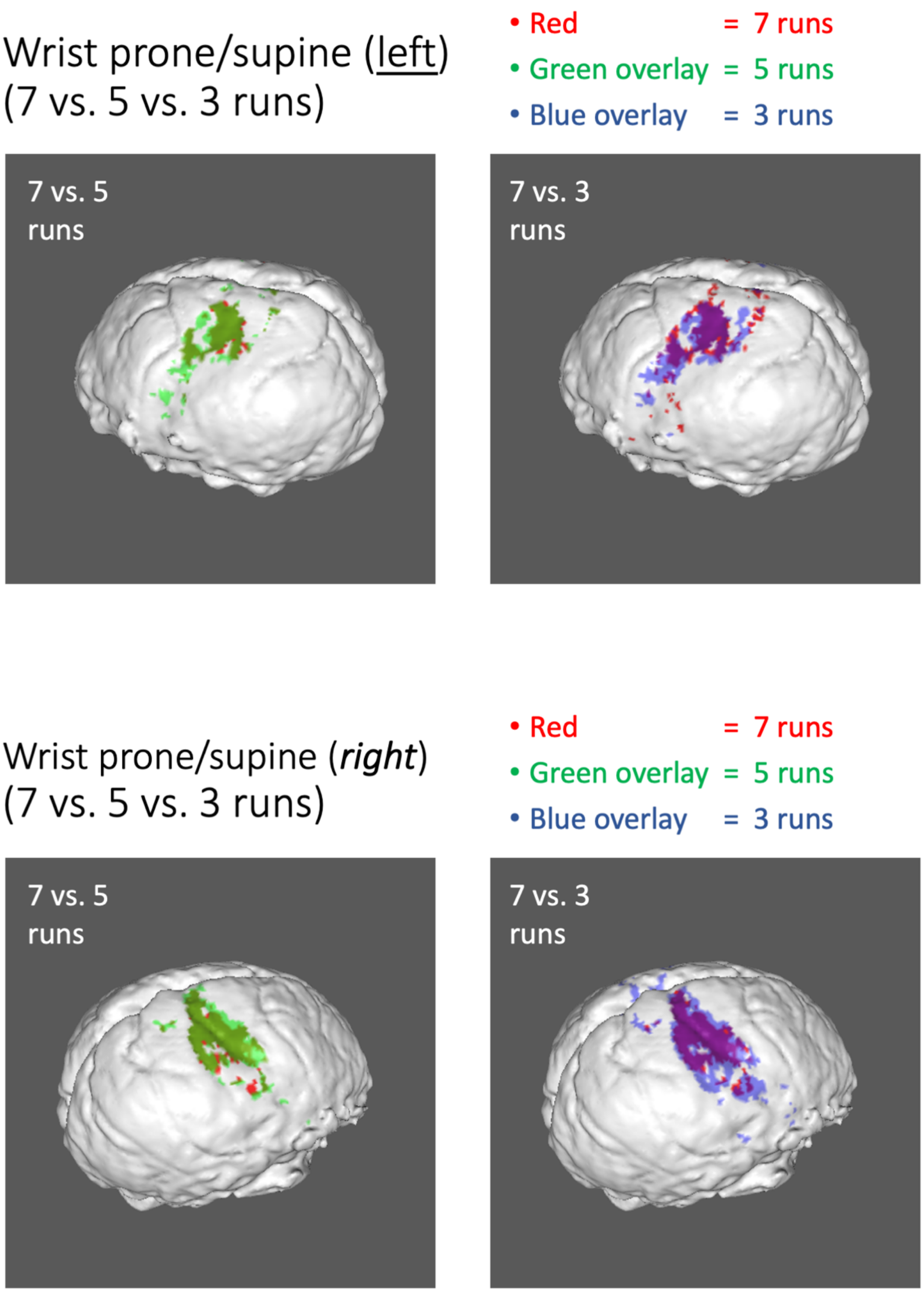


**Figure S5.6.** *3D overlays comparing runs (wrist prone/supine). Robust activations with high similarity were noted (left:* $D_c^{7-5}=0.69$*,* $D_c^{7-3}=0.56$*; right:* $D_c^{7-5}=0.78$*,* $D_c^{7-3}=0.66$*).*

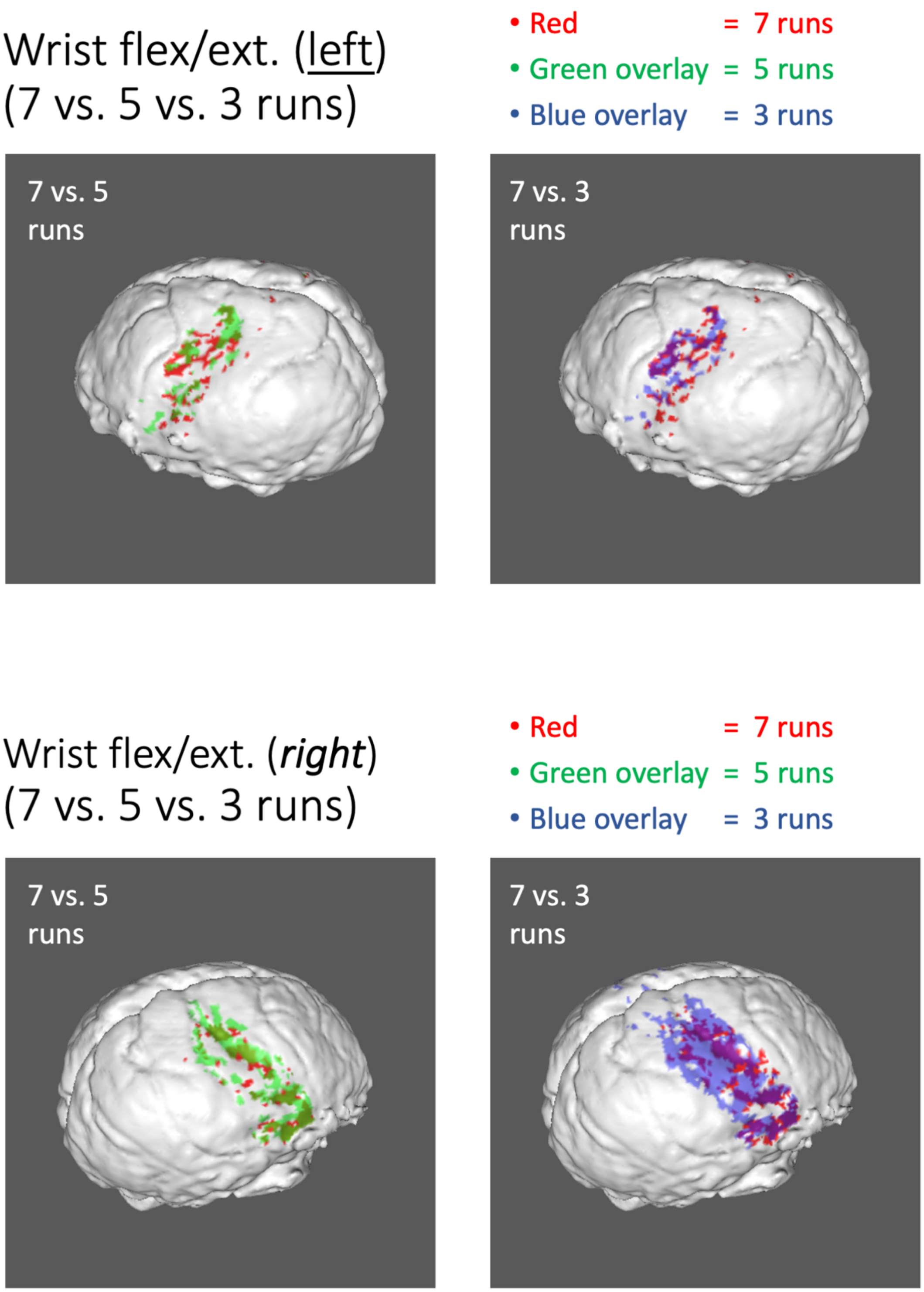


**Figure S5.7.** *3D overlays comparing runs (wrist flexion/extension). Robust activations with moderate similarity were noted (left: $D_c^{7\text{-}5}$=0.52, $D_c^{7\text{-}3}$=0.31; right: $D_c^{7\text{-}5}$=0.64, $D_c^{7\text{-}3}$=0.44).*

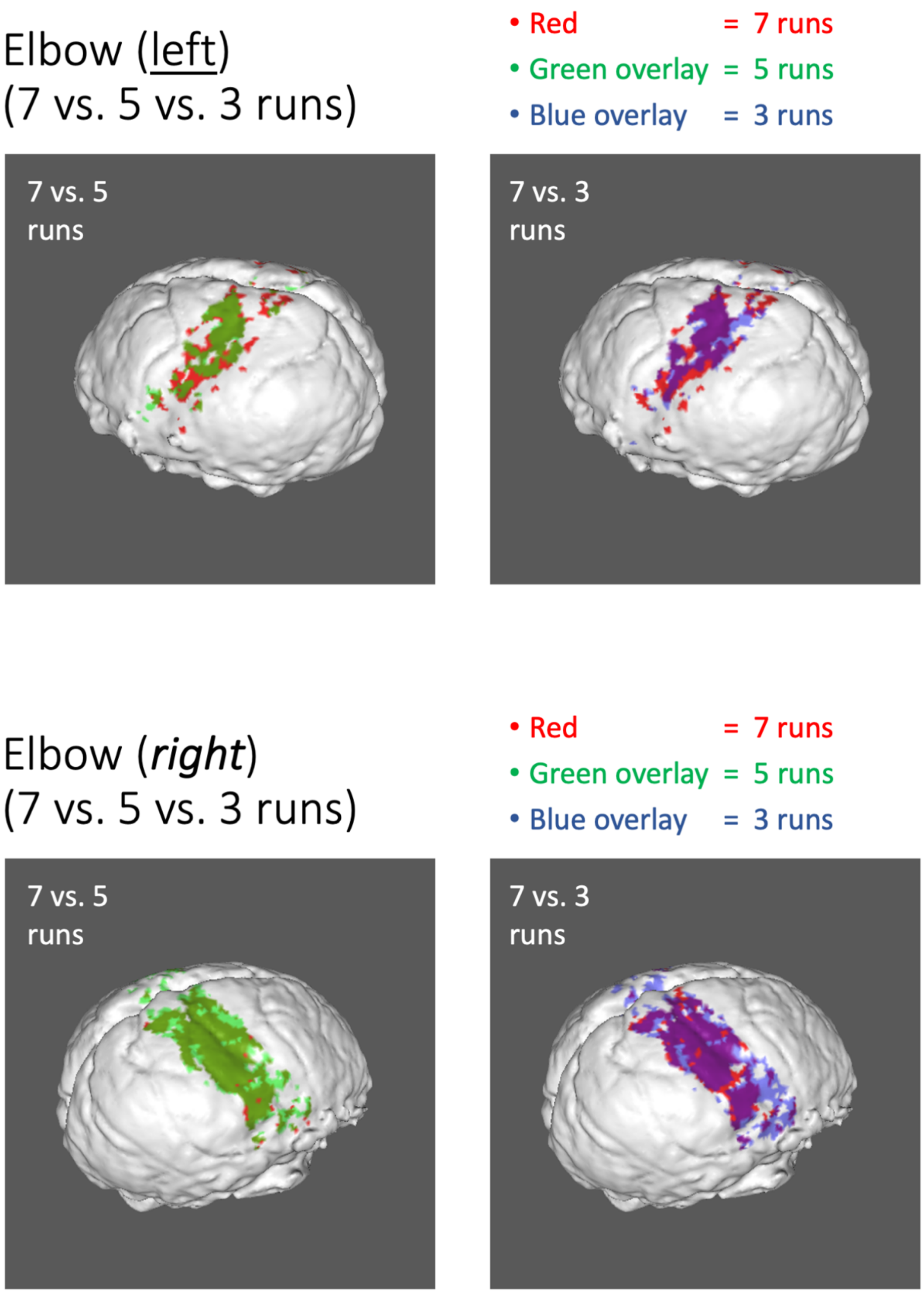


**Figure S5.8.** *3D overlays comparing runs (elbow). Robust activations with high similarity were noted (left:* $D_c^{7\text{-}5}=0.69$*,* $D_c^{7\text{-}3}=0.59$*; right:* $D_c^{7\text{-}5}=0.85$*,* $D_c^{7\text{-}3}=0.78$*).*

------- ------- -------
